# The most common human ADAR1p150 Zα domain mutation P193A is well tolerated in mice and does not activate the integrated stress response pathway

**DOI:** 10.1101/2022.06.24.497437

**Authors:** Zhen Liang, Scott Taylor, Ankita Goradia, Jacki Heraud-Farlow, Carl R Walkley

## Abstract

ADAR1 mediated A-to-I RNA editing is a self/non-self discrimination mechanism for cellular double stranded RNAs. *ADAR* mutations are one cause of Aicardi-Goutières Syndrome, an inherited paediatric encephalopathy, broadly classed as a “Type I interferonopathy”. The most common ADAR1 mutation is a proline 193 alanine (p.P193A) mutation, mapping to the ADAR1p150 isoform specific Zα domain. We report the development of an independent murine P195A knock-in mouse, homologous to the human P193A mutation. The *Adar1^P195A/P195A^* mice are largely normal and the mutation is well tolerated. Contrasting with previous reports when the P195A mutation was compounded with an ADAR1 null allele, the majority of mice have only a modest reduction in weaning weight and survived long-term. Severe runting and shortened survival of *Adar1^P195A/-^*animals are dependent on the parental genotype. The P195A mutation is well tolerated *in vivo* and the loss of MDA5 is sufficient to completely rescue the *Adar1^P195A/-^* mice.

## Introduction

One of the most common RNA modifications in mammals is the deamination of adenosine to inosine, termed A-to-I editing, in double stranded regions of RNA (dsRNA) by Adenosine Deaminase Acting on RNA 1 (ADAR1) and ADAR2 (Eisenberg and Levanon, 2018). A-to-I editing results in the non-reversible conversion from adenosine to inosine for the targeted nucleotide within the RNA. Inosine is usually interpreted as a guanosine during translation. Depending on where the editing site lies in the transcript A-to-I editing can have a variety of consequences (Eisenberg and Levanon, 2018; Solomon et al., 2017; Solomon et al., 2013; Walkley and Li, 2017). Editing within protein-coding sequences can change the amino acid codon, and therefore the protein product, from that genomically encoded (Licht et al., 2019). A-to-I editing can also impact RNA splicing, stability, translation, localization (Kapoor et al., 2020; Lev-Maor et al., 2007; Shoshan et al., 2015; Stellos et al., 2016) as well as the biogenesis of non-coding RNAs (microRNAs (Kawahara et al., 2007) and circRNAs (Ivanov et al., 2015)). Editing alters the base-pairing properties within RNA, both stabilising and destabilising the RNA secondary structure depending on context (Liddicoat et al., 2015; Solomon *et al*., 2017). A-to-I editing can be readily detected using sequencing methods, as A-to-I editing can be identified by A-to-G mismatches between the cDNA and genomic DNA, allowing genome-wide mapping (Levanon et al., 2004; Li et al., 2009; Ramaswami et al., 2013).

There are millions of A-to-I editing sites across all tissues within the human transcriptome, with the majority located in repetitive elements such as *Alu* repeats (Bazak et al., 2014; Gabay et al., 2022; Picardi et al., 2015; Tan et al., 2017). In mice there are between 50,000-150,000 editing events, also concentrated in evolutionarily related repetitive elements (SINE/LINEs) (Costa Cruz et al., 2020; Licht *et al*., 2019; Pfaller et al., 2018; Pinto et al., 2014). The key physiological function of ADAR2 is to recode the *Gria2* transcript by editing a coding region of the transcript, resulting in an amino acid substitution and a correctly functioning GLUR2 protein (Higuchi et al., 2000; Higuchi et al., 1993). In contrast to the recoding of a single relevant mRNA target for ADAR2, it has been demonstrated by multiple groups that the physiologically most important function of ADAR1 editing is to prevent the cells own RNA being mistaken as foreign RNA by the innate immune system (Liddicoat *et al*., 2015; Mannion et al., 2014; Pestal et al., 2015). ADAR1’s primary physiological function is to counteract cytoplasmic innate immune sensing by melanoma differentiation-associated protein 5 (MDA5) of endogenous RNAs (Chalk et al., 2019; Hartner et al., 2009; Heraud-Farlow et al., 2017; Liddicoat *et al*., 2015), a species conserved function (Chung et al., 2018; Pestal *et al*., 2015). ADAR1 is expressed as two isoforms, a constitutive and primarily nuclear p110 protein and an inducible and mostly cytoplasmic p150 isoform. Recent studies have focussed attention on the role of the cytoplasmic p150 isoform and how this intersects with innate immune sensing (Kim et al., 2021; Pestal *et al*., 2015; Ward et al., 2011).

*ADAR* (ADAR1) loss of function mutations have been identified as one of the genetic causes of Aicardi-Goutières Syndrome (AGS) (Rice et al., 2012; Rice et al., 2017). AGS is an inherited paediatric encephalopathy, classed as an auto-inflammatory “Type I interferonopathy”, characterised by the upregulation of interferon (IFN) and interferon stimulated gene (ISG) expression (Crow and Manel, 2015). AGS patients with *ADAR* mutations most often have compound heterozygous mutations, with one mutation impacting the p150 isoform together with a second that affects both p110 and p150 isoforms (Rice *et al*., 2012; Rice *et al*., 2017). Mutations in *ADAR* have also been identified in bilateral striatal necrosis (BSN), where patients have a dystonic or rigid movement disorder associated with symmetrical abnormalities of the brain (Livingston et al., 2014). Similar to *ADAR* mutant AGS, BSN patients with *ADAR* mutation have a characteristic “interferon signature”. Recent knock-in mouse models have begun to address how distinct ADAR1 mutations, including those reported in AGS and BSN, impact the function of ADAR1 *in vivo* (de Reuver et al., 2021; Guo et al., 2021; Inoue et al., 2021; Maurano et al., 2021; Nakahama et al., 2021; Tang et al., 2021).

One of the most frequently reported ADAR1 mutations is proline 193 to alanine (p.Pro193Ala; P193A) (Rice *et al*., 2017), which maps to the unique Zα domain of the p150 isoform (Herbert, 2021; Herbert et al., 1995; Herbert and Rich, 2001; Herbert et al., 1998; Nakahama and Kawahara, 2021). The Zα domain is found in only one other protein in the human genome, Z-DNA binding protein 1 (ZBP1), that activates cell death pathways during viral infection and in animals deficient in Receptor interaction protein kinase 1 (*Ripk1*) and caspase 8 (*Casp8*) (Jiao et al., 2020; Newton et al., 2016; Zhang et al., 2020). Proline 193 contacts the left-handed helix structure characteristic of Z-form DNA and RNA and is important in the interaction between ADAR1p150 and Z-form nucleic acid (Schwartz et al., 1999). Interestingly, unlike other *ADAR* mutations, the P193A mutation is present at an allele frequency of 0.002 of the human population globally (Herbert, 2020; Karczewski et al., 2020), however the effects of this mutation are not definitively understood. In human populations the P193A mutation is nearly always heterozygous (Karczewski *et al*., 2020). Maurano et al., recently reported development and characterisation of a murine p.Pro195Ala (P195A) mutant mouse model, homologous to the human P193A mutation (Maurano *et al*., 2021). They reported that the P195A allele was well tolerated in isolation and could be homozygous viable with a normal lifespan. When combined with a second mutation, either a p110/p150 null allele or p150 null allele, they reported a significantly shortened lifespan and reduced weaning weights, with evidence of activation of an innate immune/interferon and integrated stress response gene expression program in the *Adar1^P195A/p150-^*. They report rescue of the phenotypes by loss of MDA5 consistent with prior genetic rescue of ADAR1 loss of function alleles (de Reuver *et al*., 2021; Liddicoat *et al*., 2015; Nakahama *et al*., 2021; Pestal *et al*., 2015). However, unlike ADAR1 null/editing dead models (Liddicoat et al., 2016b; Mannion *et al*., 2014; Wang et al., 2004), they also reported normalisation of both survival and weaning weight by concurrent loss of LGP2, IFNAR or PKR.

Here, we report generation and analysis of an independent *Adar1^P195A^* mutant mouse model. Consistent with the published model (Maurano *et al*., 2021), we find that the P195A allele is well tolerated when heterozygous or homozygous. When the P195A mutation was combined with either a p110/p150 null allele or a point mutation that renders both p110 and p150 editing deficient (E861A), we observed long term survival of the majority of mice across all genotypes. Adult *Adar1^P195A/−^* and *Adar1^P195A/E861A^* animals had evidence of a mild ISG signature but not of activation of a PKR dependent integrated stress response. However, when healthy *Adar1^P195A/−^* mice were used for breeding with *Adar1^P195A/+^*, the next generation of *Adar1^P195A/−^*were severely runted with shortened lifespan. The loss of *Ifih1* (MDA5), the primary sensor of unedited cellular dsRNA, prevented activation of the ISG response and rescued long term survival of all genotypes tested. These data do not support the conclusion that the P195A allele leads to fully penetrant PKR-dependent pathology and shortened lifespan in mice.

## Materials and Methods

### Ethics Statement

All animal experiments conducted for this study were approved by the Animal Ethics Committee of St. Vincent’s Hospital, Melbourne, Australia (Protocol number #016/20). Animals were euthanized by cervical dislocation or CO_2_ asphyxiation.

### Animals

*Adar1^P195A^* mice were generated using CRISPR/Cas9 targeting in C57BL/6 zygotes by the Monash Genome Modification Platform (Monash University, Clayton, Australia). A repair oligo containing a <u>C</u>CT><u>G</u>CT point mutation resulting in a p.P195A (Proline>Alanine) mutation. The repair oligo also had a silent point mutation (CC<u>T</u>>CC<u>C</u>; Proline>Proline, p.P194P) immediately upstream and a second silent mutation (<u>T</u>TG><u>C</u>TG; Leucine>Leucine, p.L196L) immediately downstream of the P195A mutation, respectively, that created a *BsrBI* restriction enzyme site and prevented Cas9 from cutting the repaired locus and to be used for genotyping using restriction digest of the genotyping PCR product. Introduction of the mutation was confirmed by Sanger sequencing of the region in both the founders and subsequent generations.

The *Adar^E861A/+^* (*Adar1^E861A/+^*; MGI allele: *Adar^tm1.1Xen^*; MGI:5805648) (Liddicoat *et al*., 2015), *Adar^−/−^* (*Adar1^−/−^*; exon 2-13 deleted; MGI allele: *Adar^tm2Phs^*; MGI:3029862) (Hartner et al., 2004), *Adar^fl/fl^* (*Adar1^fl/fl^*; exon 7-9 floxed; MGI allele: *Adar^tm1.1Phs^*; MGI:3828307) (Hartner *et al*., 2004; Hartner *et al*., 2009), *Ifih1*^−/−^ (*Ifih1^tm1.1Cln^*) (Gitlin et al., 2006), and *Rosa26*-CreER^T2^ (Gt(ROSA)26Sor^tm1(cre/ERT2)Tyj^) (Ventura et al., 2007) mice have all been previously described and were on a backcrossed C57BL/6 background. All animals were housed at the BioResources Centre (BRC) at St. Vincent’s Hospital. Mice were maintained and bred under specific pathogen-free conditions with food and water provided *ad libitum.* For acute somatic deletion model (*R26*-CreER^T2^), all animals were ≥8 weeks of age at tamoxifen initiation; Tamoxifen containing food was prepared at 400mg/kg tamoxifen citrate (Selleckchem) in standard mouse chow (Specialty Feeds, Western Australia).

### Genotyping

Genotyping of the P195A mutants was determined by PCR. The repair oligo carrying the P195A mutation also introduced a silent *BsrB1* immediately upstream of the P195A mutation. The digestion of the genomic DNA PCR product was used to determine presence of the P195A allele and can discriminate heterozygous and homozygous mutants. The following primers: Primer P1 (5’-ACCATGGAGAGGTGCTGACG-3’) and P2 (5’-ACATCTCGGGCCTTGGTGAG-3’) were used to obtain a 489bp product from the wild-type allele, which would yield two fragments (265bp and 224bp product) when digested with *BsrB1* (NEB) for the P195A mutant. The P1 primer was used for Sanger sequencing (Australian Genome Research Facility (AGRF), Melbourne) of the purified PCR product as required. Genotyping of all other lines used was performed as previously described (Heraud-Farlow *et al*., 2017; Liddicoat *et al*., 2015).

### Histology

Three wild-type (*Adar1^+/+^*, 2 female (F)/1 male (M)), three *Adar1^P195A/+^* (2F/1M) and four *Adar1^P195A/E861A^*(3F/1M) animals at 6-7 months of age were used for histopathology examination as previously described (Chalk *et al*., 2019; Heraud-Farlow *et al*., 2017). The wild-type animals were littermate controls of the mutant bearing animals. Tissue collection and histology was performed by the Phenomics Australia Histopathology and Slide Scanning Service, University of Melbourne. The wild-type animals were identified to the pathologists as “controls”, the remaining samples were genotype blinded to the staff and pathologist assessors. The following organs were assessed: adrenal glands, bladder, bone marrow, brain, cecum, cervix, colon, duodenum, epididymes, eyes, gall bladder, harderian glands, head, heart, hind leg (long bone, bone marrow, synovial joint, skeletal muscle), ileum, jejunum, kidney, liver, lungs, mammary tissue, mesenteric lymph node, ovaries, oviducts, pancreas, penis, preputial gland, prostate glands, salivary glands and regional lymph nodes, seminal vesicles, skin, spinal cord, spleen, stomach, tail, testes, Thymus, thyroids, trachea, uterus, vagina. The full pathology report and genotypes is available in Supplemental Dataset 1.

Additional pathology was performed on 13-19 week old brain, liver, kidney and spleen isolated from wild-type (bred and housed in the same facility, not littermates of mutant animals; n=3), *Adar1^P195A/+^* (n=3), *Adar1^P195A/−^* (n=4), *Adar1^P195A/−^ Ifih1^−/−^* (n=3). Tissue was fixed overnight in 2% paraformaldehyde, transferred to 70% ethanol and then processed, sectioned and stained by the Phenomics Australia Histopathology and Slide Scanning Service, University of Melbourne. The wild-type animals were identified to the pathologists as “controls”, the remaining samples were genotype blinded to the staff and pathologist assessors. The full pathology report and genotypes is available in Supplemental Dataset 2.

### Mouse Embryonic Fibroblasts

Mouse embryonic fibroblasts (MEFs) were generated from E13.5 embryos of the indicated genotypes. The embryos were dissected, removing the head (used for genotyping), heart and foetal liver, and the remaining tissue was used to generate MEFs. The tissue was placed in 1mL of 0.025% Trypsin-EDTA (Gibco/ThermoFisher) then drawn through an 18G needle/1mL syringe, then placed at 37°C in a 10cm^2^ tissue culture plate for 30 minutes. After 30 minutes 10mL of media (High glucose DMEM (Sigma), 10% FBS (not heat inactivated, Assay Matrix), 1% Penicillin/Streptomycin (Gibco), 1% Glutamax (Gibco), 1% amphotericin B (Sigma; 250ug/mL stock)) was added to the plate and the contents dispersed. The MEFs were incubated in a hypoxia chamber flushed with 5% oxygen/5% carbon dioxide in nitrogen at 37°C. Once the cells were 70% confluent, the cells were trypsinized and passaged onto 10cm plates in normoxic conditions for all further culture. MEFs of the indicated genotypes were treated with recombinant murine interferon beta (PBL Assay Science; PBL-12405) at 250U/mL for 24hrs in normal growth media. After 24 hours, cells were collected by trypsinization and pellets washed in cold PBS and resuspended in RIPA buffer (20mM Tris·HCl, pH8.0, 150mM NaCl, 1mM EDTA, 1% sodium deoxycholate, 1% Triton X-100, 0.1% SDS) supplemented with 1x HALT protease inhibitor and 1x PhosSTOP phosphatase inhibitor (ThermoFisher). Lysates were used for western blotting as described below.

### Western blotting

Protein was quantified using the Pierce BCA protein assay kit (ThermoFisher) on an Enspire multimode plate reader (Perkin Elmer). Lysates from MEFs: indicated genotypes +/− interferon treatment were used and 40μg of protein extract per sample was loaded. Lysates from kidney: the whole kidney was homogenised in RIPA buffer containing 1x Halt Phosphatase inhibitor (ThermoFisher) and 1x Halt Protease inhibitor (ThermoFisher) and 5ug of protein was loaded. All samples were loaded on pre-cast NuPAGE™ 10%, Bis-Tris (1.5 mm, 10-well) polyacrylamide gels (Invitrogen) and transferred onto Immobilon-P PVDF membranes (Merck Millipore). Membranes were blocked with 5% milk in Tris-buffered saline with tween (TBST) and incubated at 4°C overnight with rat monoclonal anti-mouse ADAR1 antibody (clone RD4B11) (Liddicoat *et al*., 2015), phospho-eIF2α (Anti-EIF2S1 (phospho S51), Abcam, ab32157), total eIF2α (Cell Signalling Technology, 5324), mouse anti-Actin (Sigma, A1978). Membranes were then probed with HRP-conjugated goat anti-rat (ThermoFisher, 31470) or anti-mouse (ThermoFisher, 31444) secondary antibodies and visualized using ECL Prime Reagent for chemiluminescent detection on Hyperfilm ECL (Amersham).

### Peripheral blood analysis

Peripheral blood (approximately 100μl) was obtained via retro-orbital bleeding. The blood was red blood cell-depleted using hypotonic lysis buffer (150mM NH_4_Cl, 10mM KHCO_3_, 0.1mM Na_2_EDTA, pH7.3) and resuspended in 50μl of FACS buffer for flow cytometry analysis.

### Flow cytometry analysis

Antibodies against murine B220 (conjugated to APC-eFluor780), CD11b/Mac-1 (PE), Gr1 (PE-Cy7), F4/80 (APC), CD4 (eFluor450) and CD8a (PerCP-Cy5.5) were used to measure granulocyte, macrophage, B lymphocyte, and T cell populations. The measurement of erythroid cells used antibodies against murine Ter119 (PE), CD71 (APC), CD44 (PE-Cy7). Sca-1(PerCP-Cy5.5),c-Kit (APC-eFluor780), CD150 (PE), CD48 (PE-Cy7), CD34(eFluor660), CD16/32 (eFluor450) and biotinylated antibodies (CD2, CD3e, CD4, CD5, CD8a, B220, Gr-1, CD11b/Mac1) were used to quantify the lineage negative fraction of bone marrow as described (Heraud-Farlow *et al*., 2017; Liddicoat *et al*., 2015; Singbrant et al., 2011; Smeets et al., 2014). Biotinylated antibodies were detected with streptavidin conjugated Brilliant Violet 786. Antibodies were purchased from eBioscience, BioLegend or BD Pharmingen. Cells were acquired on a BD LSRIIFortessa and analyzed with FlowJo software Version 9 or 10.0 (Treestar).

### RT-qPCR

Whole bone marrow and myeloid cells +/− tamoxifen treatment were collected, and RNA isolated using the RNeasy kit with on-column DNase digestion (Qiagen). Mouse tissues were collected (one brain hemisphere, liver, kidney; additional histology performed on tissue from these same animals) from independent biological replicates from 14-19 week old C57BL/6 (n=3; 1F/2M; bred and housed in same facility), *Adar1^P195A/+^* (n=3; 1F/2M), *Adar1^P195A/−^* (n=4; 2F/2M), *Adar1^P195A/+^ Ifih1^−/−^* (n=3; 2F/1M) and *Adar1^P195A/−^ Ifih1^−/−^* (n=3; 2F/1M). Additional tissue was collected from a separate cohort of runted *Adar1^P195A/−^*and controls [*Adar1^+/−^* (1F; 3 wks old), *Adar1^P195A/+^* (2M; one 3 wks old, one 4 wks old), *Adar1^P195A/P195A^* (2F; one 3 wks old, one 10 wks old), non-runted *Adar1^P195A/−^*(2M; both 4 wks old), runted *Adar1^P195A/−+^* (2M/1F; two 3 wks old, one 10 wk old)]. Tissues were collected and immediately snap frozen in liquid nitrogen and stored at −80C. Frozen tissues were homogenized in Trisure reagent (Bioline) using IKA T10 basic S5 Ultra-turrax Disperser. RNA was extracted using Direct-Zol columns (Zymo Research) as per manufacturer’s instruction. cDNA was synthesized using Tetro cDNA synthesis kit (Bioline; used for all cDNA described).

Real-time PCR was performed using two methods: SYBR green method (for myeloid cell lines and *in vivo R26*-CreER experiments, normalised to *Ppia*; Fig 5 and Fig 6’ primer sequences in Supplementary Table 1) as previously described (Heraud-Farlow *et al*., 2017). Alternatively predesigned Taqman probes for *Oas1a, Ifi27, Irf7, Asns, Cdkn1a, Hmox1* were used and normalised to *Ppia* and *Hprt* (for mouse organs from germ-line mutant animals in Fig 4). Duplicate reactions per sample were measured using an AriaMx Real-time PCR machine (Agilent) using TaqMan Fast Advanced Master Mix (Applied Biosystems, ThermoFisher) and predesigned/inventory FAM conjugated Taqman primer/probe sets (Applied Biosystems, ThermoFisher) against murine genes: *Oas1a* (assay ID Mm00836412_m1); *Ifi27* (Mm00835449_g1); *Irf7* (Mm00516788_m1); *Asns* (Mm00803785_m1); *Cdkn1a* (Mm04205640_g1); *Hmox1* (Mm00516005_m1) and control gene *Hprt* (Mm00446968_m1). *Hprt* was used as reference genes for relative quantification using the ΔΔCt method (similar results using *Ppia* as reference gene; data not shown).

### Immortalized myeloid cells

HOXA9 immortalized myeloid cell lines (Wang et al., 2006) were established by retroviral infection of ficoll depleted bone marrow isolated from three independent adult (>8week old) *R26*-CreER^T2^ *Adar1^fl/+^*and *R26*-CreER^T2^ *Adar1^fl/P195A^* donor animals. The cells were cultured in IMDM (Sigma) containing 10% FBS (Assay Matrix; non-heat inactivated), 1% Penicillin/Streptomycin (Gibco/ThermoFisher), 1% Glutamax (Gibco/ThermoFisher) supplemented with 50ng/mL recombinant mouse stem cell factor (rmSCF, Peprotech), 10ng/mL recombinant mouse interleukin 3 (rmIL-3, Peprotech) and 10ng/mL recombinant human interleukin 6 (rhIL-6, Amgen) for 48 hrs. After 48hr in culture, 1×10^6^ cells were spin-infected at 1,100g for 90 minutes with ecotrophic packaged HOXA9 retrovirus (HOXA9 plasmid was generously provided by Dr Mark Kamps, University of California San Diego) and 8ug/mL hexadimethrine bromide (Polybrene; Sigma). At 48 hrs post-infection, the cells were passaged into IMDM containing 10% FBS, 1% Penicillin/Streptomycin, 1% Glutamax supplemented with 1% granulocyte-macrophage colony-stimulating factor (GM-CSF) conditioned medium (from BHK-HM5 cell conditioned medium). Cells were maintained in GM-CSF containing media after this point. Cell lines established after 3-4 weeks of culture.

Once stable cell lines were established, they were treated with 200nM 4-hydroxy tamoxifen (Merck Millipore) to activate CreER recombination which results in cells becoming *Δ/+* (*fl/+* cells become *Adar1* heterozygous) or *⊗/P195A* (*fl/P195A* cells only retain expression of P195A after tamoxifen treatment). Cells were counted with Trypan blue using a Countess II automated counter (ThermoFisher) and then passaged every 2-3 days. Viability was assessed by trypan blue staining and counted using a Countess II and gene expression on cDNA made from cells of the indicated genotypes and assessed by SYBR green based qPCR as described previously (Chalk *et al*., 2019; Heraud-Farlow *et al*., 2017).

To test if there is an aberrant response in P195A mutant cells toward non-cellular cytosolic dsRNA, *Adar1^11/+^* and *Adar1^11/P195A^* cells were treated with high molecular weight polyinosinic:polycytidylic acid (poly(I:C); InvivoGen). *R26-*CreER *Adar1^fl/+^* and *R26-*CreER *Adar1^fl/P195A^* cells were treated for 14 days with tamoxifen, genotyped, and then *Adar1^11/+^* and *Adar1^11/P195A^* cells were transferred to non-tamoxifen supplemented IMDM media for poly(I:C) testing. Three independent cell lines were used for each genotype. The cells were treated with a dose range of high molecular weight poly(I:C) (1.5-8kb; 1mg/mL stock) by nucleofection following manufacturer’s instructions (Lonza 4D-Nucleofector^TM^ Kits). Gene expression of the indicated genotypes was assessed by SYBR green based qPCR as described previously.

### Statistical analysis

To determine statistical significance, Kaplan-Meier survival plots and ordinary one-way ANOVA tests were conducted in GraphPad Prism software version 9 (GraphPad; San Diego, CA, USA). Throughout this study, significance is indicated using the following convention: *P<0.05; **P<0.01; ***P<0.001; ****P<0.0001, and data is presented as mean ± S.E.M. The number of samples used for each experiment is described in the corresponding figure legends.

## Results

### Generation of an *Adar1* P195A knock-in mutant allele

The P193A mutation in human ADAR1 maps to the Zα domain that is unique to the ADAR1 p150 isoform. The proline 193 residue of human ADAR1 is homologous to murine proline 195 (Fig 1A). We introduced a C to G point mutation to generate a proline to alanine substitution at amino acid 195 (p.P195A) into murine *Adar1* using CRISPR/Cas9 on a C57BL/6 background (Fig 1B). This resulted in the desired mutation that was confirmed by Sanger sequencing and through restriction digest of genomic PCR products, utilising a silent unique restriction site that was introduced during targeting to the modified locus (Fig 1C). After identification of heterozygous founder mice (*Adar1^P195A/+^*), these were bred to C57BL/6 mice to confirm germ-line transmission of the mutant allele. Second generation animals were subsequently bred for all experiments.

**Figure 1.**
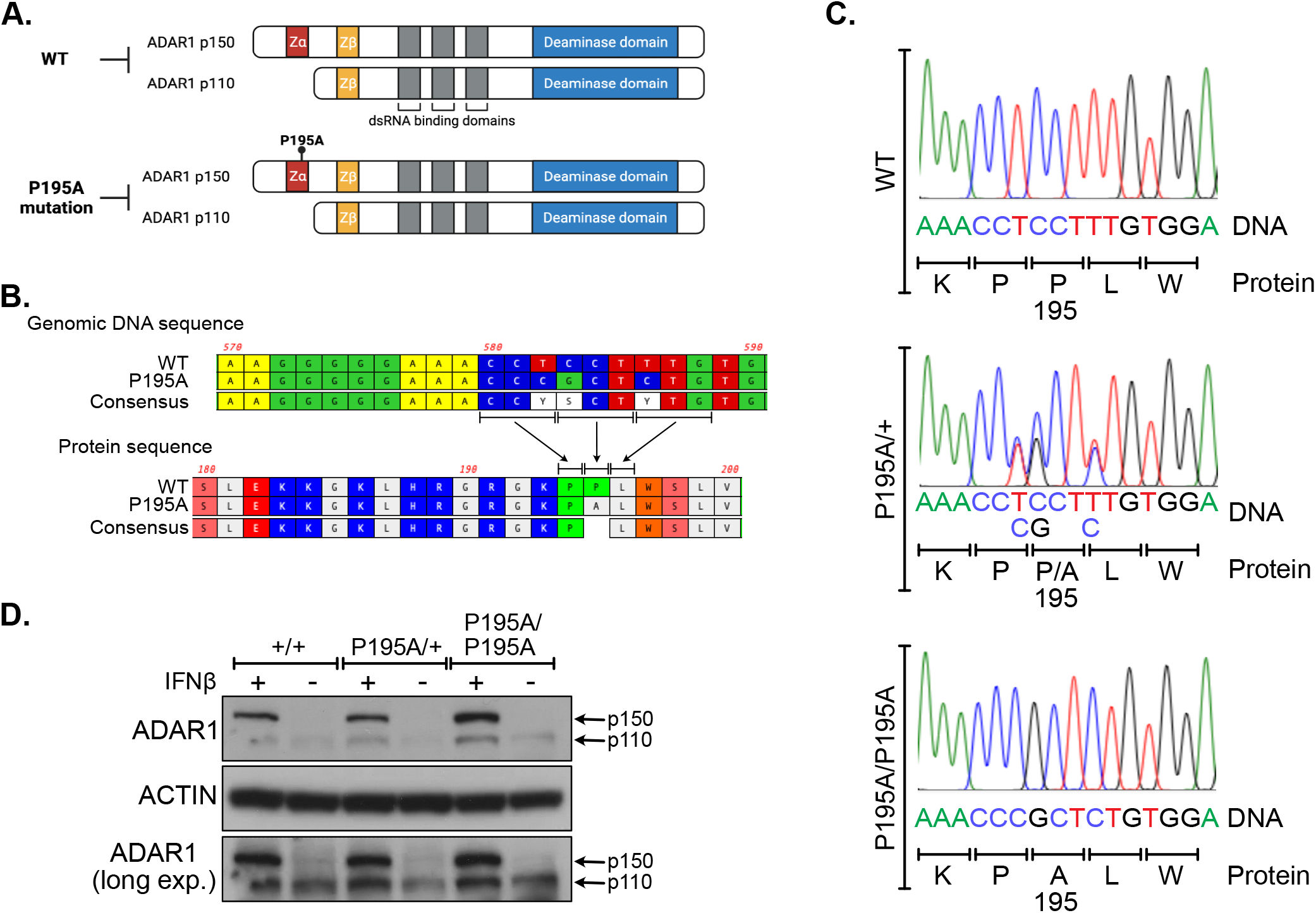
Generation of an *Adar1^P195A^* knock-in allele. **A)** Schematic of the P195A mutation. **B)** Genomic alignment and translation of the WT and P195A allele. During introduction of the P195A mutation, two additional silent mutations were introduced immediately upstream and downstream, respectively. These were to allow restriction digestion of the mutant allele and to prevent re-cutting by CAS9 during targeting. **C)** Sanger sequencing traces and alignments of genomic DNA isolated from animals of the indicated genotypes. **D)** Western blot analysis of ADAR1 expression in E13.5 mouse embryonic fibroblasts (MEFs) of the indicated genotypes +/− interferon-β (IFNβ) treatment. IFNβ was used to induce expression of the p150 isoform of ADAR1.

To assess expression of the ADAR1p150 protein from the mutant allele, we isolated E13.5 mouse embryonic fibroblasts (MEFs) and treated these with murine interferon beta (IFNβ). We tested MEFs from two independent *Adar1^+/+^*, *Adar1^P195A/+^* and *Adar1^P195A/P195A^* littermates. This demonstrated that the induction of the p150 isoform by interferon was intact and equivalent between *Adar1^+/+^*, *Adar1^P195A/+^* and *Adar1^P195A/P195A^* cells (Fig 1D).

### ADAR1 P195A mutation is well tolerated

The heterozygous *Adar1^P195A/+^* mice were intercrossed and we recovered viable homozygous animals at the expected frequency, consistent with a previous report (Fig 2A, Supplemental Figure 1B) (Maurano *et al*., 2021). To understand the consequences of a P195A mutation compounded with a second mutation, more similar to the mutational spectrum reported in AGS (Rice *et al*., 2017), we crossed the *Adar1^P195A/+^* mice to an *Adar1^+/−^* (*Adar1* null allele, exon 2-13 deleted) (Hartner *et al*., 2004) and, in parallel, to an editing deficient point mutant model (*Adar1^E861A/+^*) (Liddicoat *et al*., 2015) (Supplemental Fig 1). We recovered mice with all possible genotypes from these intercrosses (Fig 2B and data not shown).

**Figure 2.**
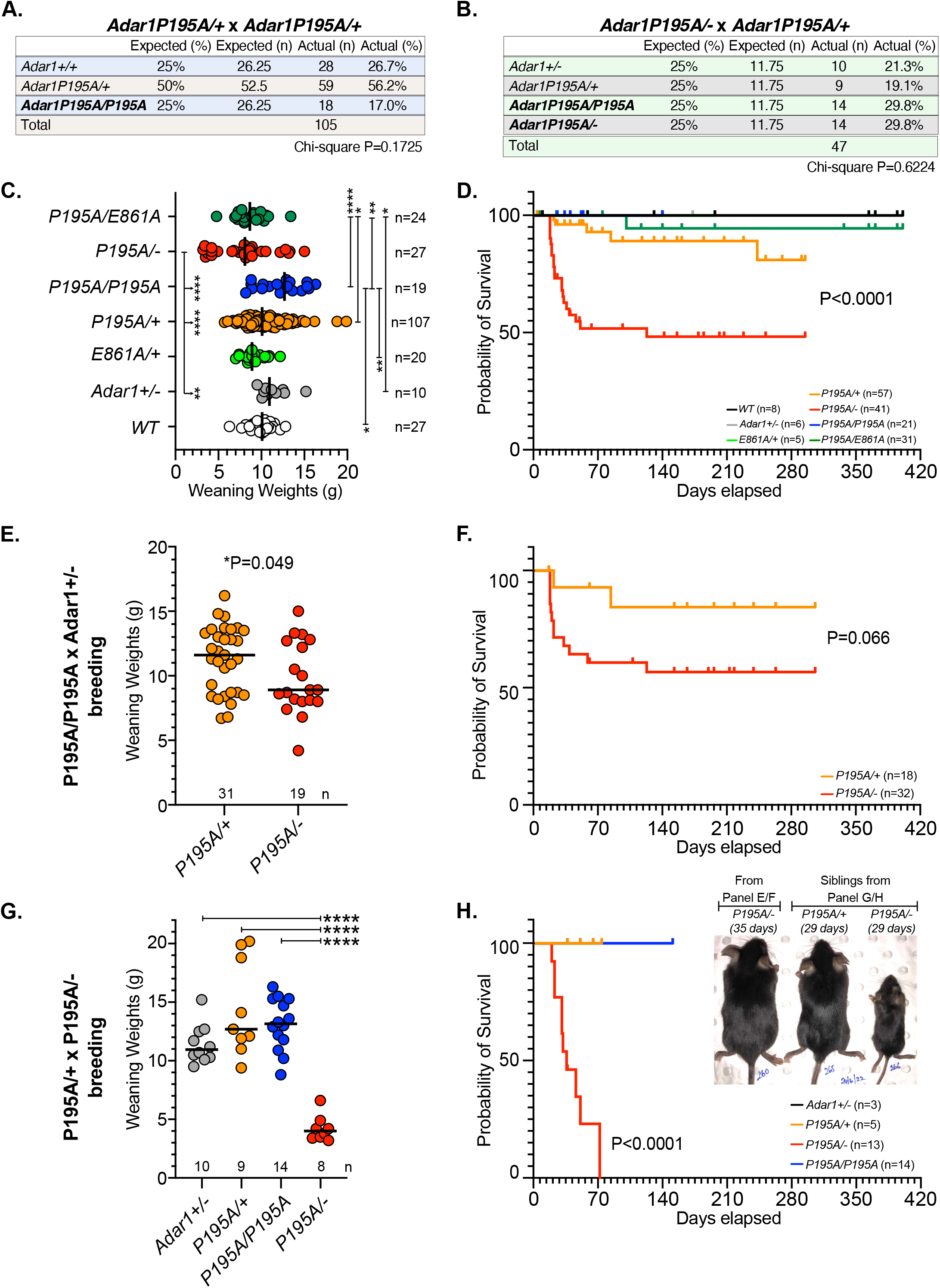
Long term survival of *Adar1^P195A/−^* animals. **A)** Results from inbreeding of *Adar1^P195A/+^* animals. **B)** Results from breeding of *Adar1^P195A/+^* animals with *Adar1^P195A/−^* animals. **C)** Weaning weights of mice of the indicated genotypes. *P<0.05; **P<0.01; ***P<0.001; ****P<0.0001; Ordinary one-way ANOVA with Tukey’s multiple comparison test (adjusted P value). **D)** Survival analysis of mice of the indicated genotypes; number as indicated for each genotype. Statistical difference determined by Log-rank (Mantel-Cox) test. **E)** Weaning weights and **F)** survival analysis of pups derived from breeding of a *Adar1^P195A/P195A^* to a *Adar1^+/−^* (*Adar1* germ-line deficient heterozygous) animal; number as indicated for each genotype. Statistical analysis of weight using unpaired t test (two sided); of survival using Log-rank (Mantel-Cox) test. **G)** Weaning weights and **H)** survival analysis of pups derived from breeding of a *Adar1^P195A/−^* to a *Adar1^P195A/+^* animal; number as indicated for each genotype. Inset photo – 35 day old *Adar1^P195A/−^*male bred from *Adar1^P195A/P195A^* x *Adar1^+/−^* parents (data in Panel E/F); sibling 29 day old *Adar1^P195A/+^* and *Adar1^P195A/−^*male bred from an *Adar1^P195A/−^* x *Adar1^P195A/+^*(data in Panel G/H). ****P<0.0001; for weight analysis Ordinary one-way ANOVA with Tukey’s multiple comparison test (adjusted P value); survival using Log-rank (Mantel-Cox) test. Data displayed as individual animals with mean indicated. Weaning weights not available if animal was found dead prior to weaning; all animals where genotype was confirmed are included in the survival analysis (where possible any found dead prior to weaning were genotyped post-mortem).

We assessed the weaning weights of the different mutants compared to controls genotypes. The *Adar1^P195A/P195A^* animals had a higher average weaning weight than wild-type *Adar1^+/+^* mice (*Adar1^P195A/P195A^*mean weight = 12.2g, n=19; *Adar1^+/+^* mean weight = 9.9g, n=27; Fig 2C) and higher weaning weights compared to the *Adar1^P195A/−^* (mean = 8.12g), *Adar1^E861A/+^*(mean = 9.13g), and *Adar1^P195A/E861A^* (mean = 8.72g) cohorts (Fig 2C). The *Adar1^P195A/−^*had significantly lower weaning weights compared to the *Adar1^+/−^*, *Adar1^P195A/+^* and *Adar1^P195A/P195A^*. The *Adar1^P195A/E861A^* mice, where the P195A mutation is paired with the editing deficient E861A mutant p110 and p150 protein, were not significantly different from the *Adar1^P195A/−^* cohort and were lighter than the *Adar1^+/−^*, *Adar1^P195A/+^*and *Adar1^P195A/P195A^* animals (Fig 2C).

Maurano et al., reported fully penetrant runting of their *Adar1^P195A/−^* animals, which contrasted with the variability in weights that we observed in our *Adar1^P195A/−^* cohort (Maurano *et al*., 2021). While the *Adar1^P195A/−^*cohort as a whole had significantly lower weaning weights than most other genotypes, they appeared to be clustered into two distinct groups of low or approximately normal weaning weight (Fig 2C). The mean weaning weights across all genotypes in our study (mean range from 8.12 to 12.2g; sexes combined) are consistent with the reference data for 3-4 week old animals of C57BL/6 background mice from both the Jackson Labs and the International Mouse Phenotyping Consortium (Dickinson et al., 2016) (Supplemental Table 2). The wildtype controls reported by Maurano et al. weighed more than 20g at 23 days of age, indicating that they differed from strains we and others have utilised. Upon further analysis, the *Adar1^P195A/−^* animals derived from crosses where a *Adar1^P195A/−^* was used as a parent were consistently runted (mean weaning weight of *Adar1^P195A/−^* 4.23g; n=8; Fig 2E, 2G). In contrast, this was largely ameliorated when a *Adar1^P195A/P195A^* was bred to a *Adar1^+/−^* animal (mean weaning weight of *Adar1^P195A/−^*9.76g; n=19; Fig 2E, 2G). This result suggested that the breeding pair genotype influenced the weaning weight of the compound mutant pups.

Next, we compared the long-term survival of the different genotypes. Cohorts of mice were allowed to age and monitored for any signs of illness or change in health status. The *Adar1^P195A/+^* and *Adar1^P195A/P195A^* mice did not have any change in long term survival (Fig 2D), consistent with that reported (Maurano *et al*., 2021). Similarly, when we assessed the survival of the *Adar1^P195A/E861A^* animals we found that these mutants survived normally long term (>450 days). The initially reported *Adar1^P195A/−^*model described the P195A allele crossed with a germ-line *Adar1* allele, derived from the germ-line deletion of the floxed Exon 7-9 allele, that was originally reported as null, but later found to express a truncated, unstable, mislocalized editing deficient protein (Bajad et al., 2020; Hartner *et al*., 2004; Hartner *et al*., 2009; Pestal *et al*., 2015). Around 90% of these compound heterozygote *Adar1^P195A/−^* mice died by one month of age and only a single animal survived to day 84 (Maurano *et al*., 2021). In our facility and with our *Adar1* null allele (exon 2-13 deletion), the *Adar1^P195A/−^* mice have survived long term, with >50% of the *Adar1^P195A/−^* animals surviving past 120 days of age, and the current oldest >350 days of age (Fig 2D). Since it appeared that there were two distinct groups of *Adar1^P195A/−^* mice based on both weaning weight (low or normal, Fig 2C) and survival (die by 60 days or long-term survival, Fig 2D) we further assessed survival based on the breeding pair genotypes. We noted poor survival and outcomes for pups bred from breeding pairs including a *Adar1^P195A/−^*parent (2 female and one male *Adar1^P195A/−^* used for breeding to date; Fig 2F, 2H). This result suggested that a significant contributor to the phenotype of the pups was the genotype of the parent, with a *Adar1^P195A/−^* parent resulting in runting and poor survival of *Adar1^P195A/−^* pups. The *Adar1^P195A/−^*genotype more closely approximates that of an AGS patient, and we cannot find literature to indicate if humans with similar genotypes can successfully rear children. Collectively these data indicate that the P195A mutation is well tolerated and is compatible with long term survival, even when compounded with either an ADAR1 null allele or editing deficient mutation. The data further indicate that the increased post weaning mortality of the *Adar1^P195A/−^*animals can be fully prevented by provision of an editing deficient, protein expressing allele.

When assessed as adult mice (>8 weeks old), both the *Adar1^P195A/−^*and *Adar1^P195A/E861A^* were macroscopically normal. To determine if there were any microscopic changes, a histopathological assessment of a cohort of *Adar1^+/+^* (wild-type littermates; n=3), *Adar1^P195A/+^*(n=3) and *Adar1^P195A/E861A^* (n=4) at 6-7 months of age was undertaken as we have previously described (Chalk *et al*., 2019; Heraud-Farlow *et al*., 2017). From this analysis there were no genotype specific differences between the groups, nor evidence for pathological changes within the tissues assessed (Supplemental Dataset 1). We also assessed the brain, kidney, liver and spleen from a cohort of 13-19 week old animals containing C57BL/6 mice (n=3; bred and housed in the same facility), *Adar1^P195A/+^*(n=3), *Adar1^P195A/−^* (n=4), and *Adar1^P195A/−^Ifih1^−/−^*(n=3; described below, Supplemental Dataset 2). This did not find any genotype specific defects or pathological changes.

### Loss of MDA5 is sufficient to prevent innate immune activation

Based on the demonstrated role of MDA5 in sensing and responding to unedited cellular dsRNA (Liddicoat *et al*., 2015), we sought to determine if loss of MDA5 (gene *Ifih1*) would prevent the lethality we had observed in the *Adar1^P195A/−^* in the first 100 days of life. We generated cohorts of animals on both an *Ifih1^+/−^* or *Ifih1^−/−^*background. Strikingly, both heterozygosity for MDA5 or full deletion was able to prevent the death we had observed in a subset of *Adar1^P195A/−^*animals (Fig 3A-3B, Supplemental Fig 2A-B). Furthermore, heterozygous or homozygous loss of MDA5 normalised the weaning weights across all genotypes assessed. This was especially the case for the *Adar1^P195A/−^*where the *Adar1^P195A/−^ Ifih1^−/−^* weights were less variable than on an *Ifih1^+/+^* background (*Adar1^P195A/−^*mean weight = 8.12g, n=27; *Adar1^P195A/−^Ifih1−/−* mean weight = 10.63g, n=16; Fig 3A-3B, Supplemental Fig 2A-B). The *Adar1^P195A/−^Ifih1^+/−^* and *Adar1^P195A/−^Ifih1^−/−^*weights were not significantly different from any other cohort on an MDA5 heterozygous or null background. (Fig 3A-3B, Supplemental Fig 2A-B). This demonstrates that loss of MDA5 alone, even as a heterozygous mutation, is sufficient to restore viability and long-term survival of the *Adar1^P195A/−^* and *Adar1^P195A/E861A^* animals.

**Figure 3.**
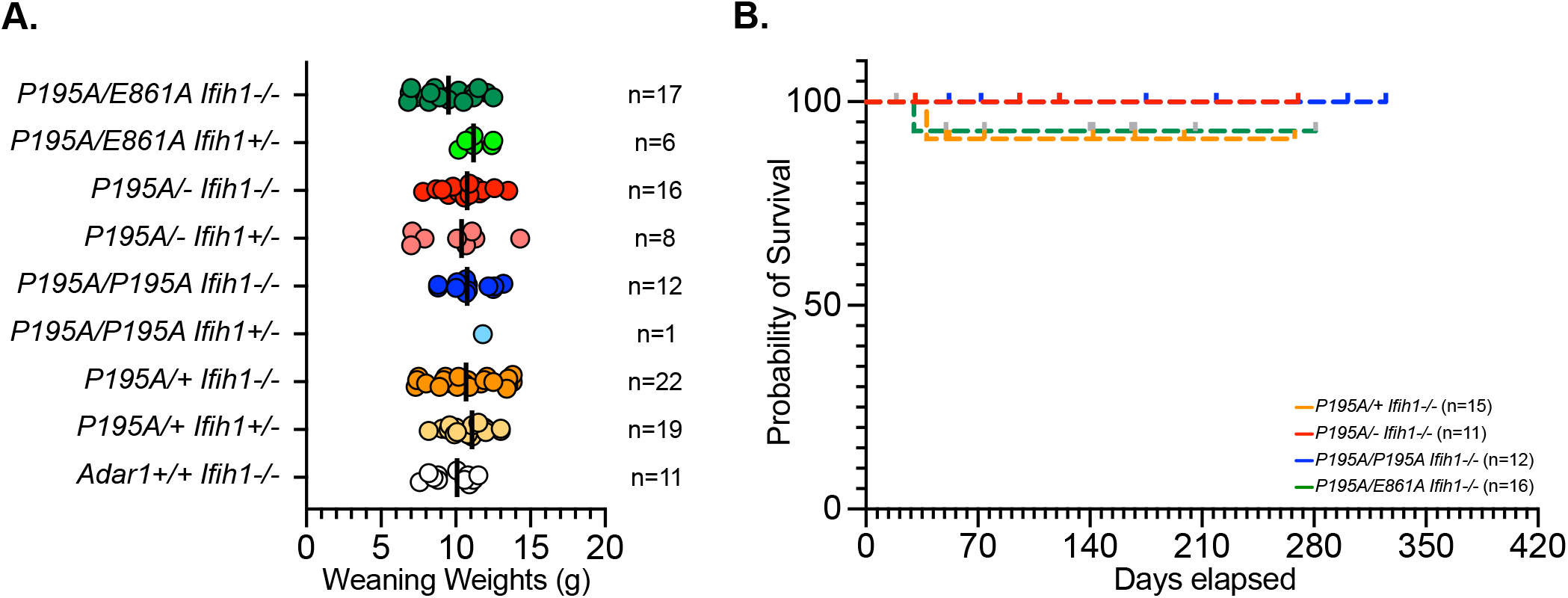
Loss of MDA5 rescues both weight and viability of *Adar1^P195A/−^*. **A)** Weaning weights of mice of the indicated genotypes on an MDA5 heterozygous (*Ifih1^+/−^*) or null (*Ifih1^−/−^*) background. No statistically significant difference across any comparison (Ordinary one-way ANOVA with Tukey’s multiple comparison test). **B)** Survival analysis of mice of the indicated genotypes (all *Ifih1^−/−^*); number as indicated for each genotype. Survival analysis for all *Ifih1^+/−^* and *Ifih1^−/−^*genotypes in Supplemental Fig 2B. No statistical difference between genotypes by Log-rank (Mantel-Cox) test. Data displayed as individual animals with mean indicated. Weaning age was determined by animal facility staff independently of investigators based on animal welfare and facility SOPs. Animals typically weaned at 20-25 days of age; number as indicated for each genotype. Weaning weights not available if animal was found dead prior to weaning; all animals where genotype was confirmed are included in the survival analysis (where possible any found dead prior to weaning were genotyped post-mortem).

### Compound *Adar1^P195A/−^* and *Adar1^P195A/E861A^* have a mild activation of ISGs but not of ISRs

We next sought to determine if the P195A mutation, which is specific to the cytoplasmic p150 isoform of ADAR1, altered the cellular innate immune response. We isolated RNA from brain, liver and kidney tissues from 13-19 week old C57BL/6 mice (n=3; bred and housed in the same facility; same cohort subjected to histopathology), *Adar1^P195A/+^* (n=3), *Adar1^P195A/−^* (n=4), *Adar1^P195A/E861A^* (n=3), *Adar1^P195A/−^Ifih1^+/−^* (n=3) and *Adar1^P195A/−^Ifih1^−/−^* (n=3) and used the same Taqman based qPCR assays as described in the original P195A mouse model description (Maurano *et al*., 2021) to assess the expression of the ISGs *Oas1a*, *Ifi27* and *Irf7* (Fig 4A). We found that the *Adar1^P195A/−^*and the *Adar1^P195A/E861A^* had mildly elevated expression of all three ISGs with variability in the level of increased expression between individuals and tissues (Fig 4), consistent with that reported by Maurano and colleagues (Maurano *et al*., 2021) and in other Zα domain mutants (de Reuver *et al*., 2021; Nakahama *et al*., 2021; Tang *et al*., 2021). The induction of the ISGs was completely prevented by the deletion of MDA5 (Fig 4), consistent with the known function of MDA5 as the primary sensor of under-edited/unedited cellular dsRNA (Liddicoat *et al*., 2015; Mannion *et al*., 2014; Pestal *et al*., 2015). Based on the proposed activation of the integrated stress response (ISR) in the previously reported P195A model (Maurano *et al*., 2021), we assessed the expression of the ISR genes *Asns*, *Cdkn1a*, *Hmox1* using Taqman based qPCR on the same samples assessed for ISGs. In contrast to that reported (Maurano *et al*., 2021), we do not find evidence for an increase or activation of the ISR as determined from the expression of these genes across the genotypes tested (Fig 4B). We further assessed ISG and ISR expression by qPCR in the brain of runted *Adar1^P195A/−^*(n=3; between 3 and 10 weeks of age) compared to littermate non-runted *Adar1^P195A/−^*(n=2) and other control genotypes (Fig 4C-D; Supplemental Fig 3). We saw much higher induction of ISG expression in the runted *Adar1^P195A/−^* compared to the *Adar1^P195A/−^* non-runted. However, consistent with the 13-19 week old animals, we did not see any altered expression of genes in the integrated stress response pathway (Fig 4D). In summary, the compound mutation of P195A with either the null allele (*Adar1^P195A/−^*) or the editing dead allele (*Adar1^P195A/E861A^*) leads to the activation of an MDA5 dependent ISG response. We do not find evidence for activation of the integrated stress response and see complete genetic rescue by the loss of MDA5 *in vivo*.

**Figure 4.**
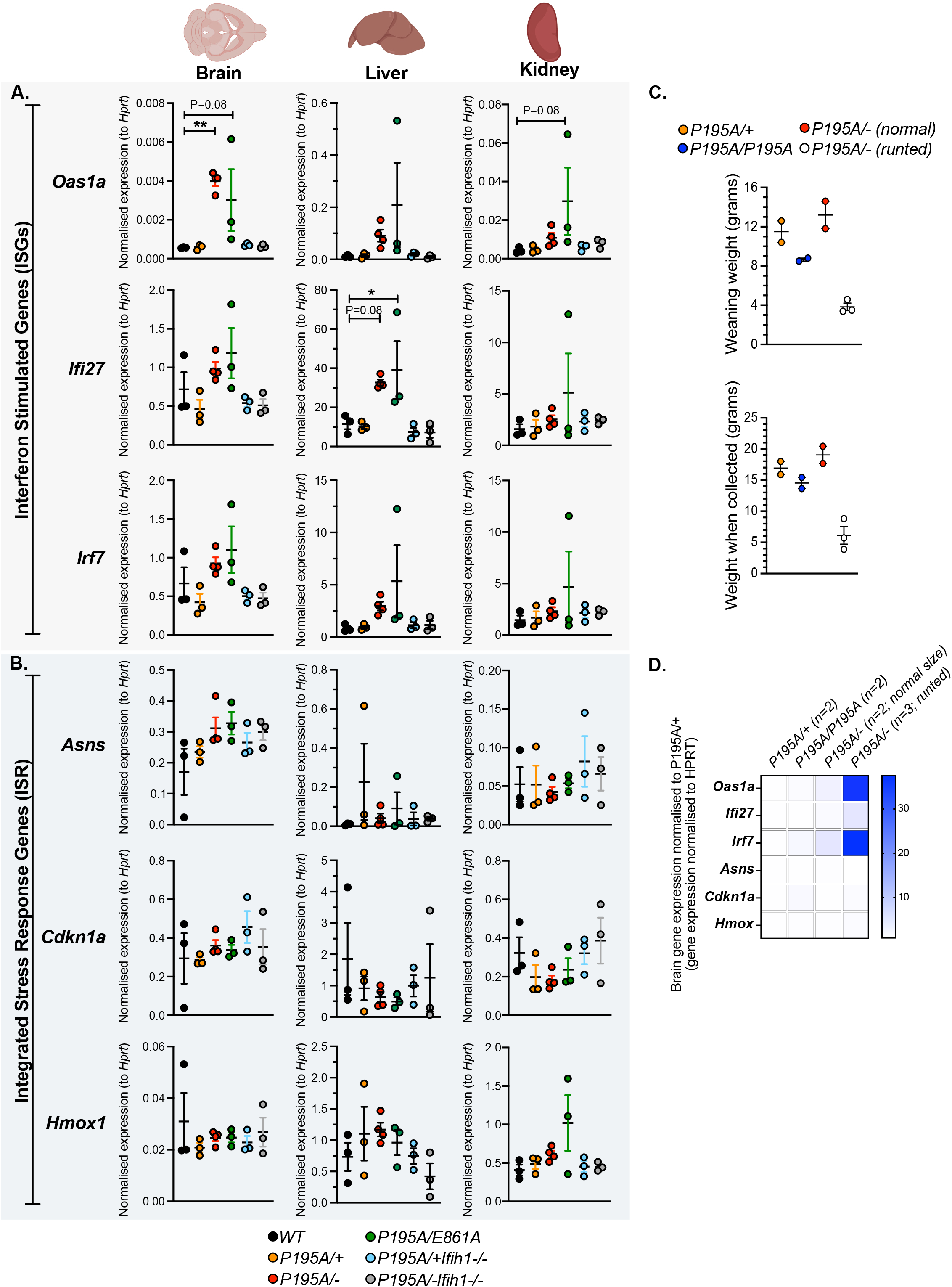
Mild tissue-specific upregulation of interferon stimulated gene expression in *Adar1^P195A/−^* and *Adar1^P195A/E861A^* is MDA5 dependent. **A)** Taqman based qPCR for the interferon stimulated genes *Oas1a*, *Ifi27* and *Irf7* in adult brain, liver and kidney of mice of the indicated genotypes; number of independently derived samples as indicated for each genotype. **B)** Taqman based qPCR for the integrated stress response genes *Asns*, *Cdkn1a* and *Hmox1* in adult brain, liver and kidney of mice of the indicated genotypes; number of independently derived samples as indicated for each genotype. Data expressed as mean +/− SEM gene expression relative to *Hprt* expression. N=3 per genotype; Statistical analysis for all qPCR: Ordinary one-way ANOVA with Dunnett’s multiple comparison test; *P<0.05; **P<0.01 or P as indicated. No P value indicates not statistical significance. **C)** Weights of mice of the indicated genotypes at weaning and when collected. The mice were used for Taqman based qPCR in panel D and Supplemental Figure 3. **D)** Heatmap representation of the expression of interferon stimulated genes (*Oas1a*, *Ifi27* and *Irf7*) and the integrated stress response genes (*Asns*, *Cdkn1a* and *Hmox)* in brain of mice of the indicated genotypes using Taqman based qPCR.

### Acute expression of ADAR1 P195A alone is well tolerated *in vitro*

To assess the consequence of somatic restricted P195A expression, we established HOXA9 immortalised myeloid cell lines (Wang *et al*., 2006) using bone marrow from adult (>8 week old) *R26*-CreER^T2^ *Adar1^fl/+^* and *R26*-CreER^T2^ *Adar1^fl/P195A^*animals (Fig 5A). These cell lines become ADAR1 hetrozygous (*fl/+* becomes *⊗/+*) or ADAR1 P195A only expressing (*fl/P195A* becomes *⊗/P195A*) after treatment with tamoxifen (Fig 5B). Notably, these cells employ the same floxed *Adar1* exon 7-9 allele used by Maurano and colleagues. We used this model to assess the consequences of acute deletion of the floxed *Adar1* allele and expression of only P195A. We cultured the cells with and without tamoxifen over a 14-day time course. We achieved complete deletion of the *Adar1* floxed allele as assessed by genotyping (Fig 5C). The proliferation and viability of the P195A cells was not affected by treatment with tamoxifen and the cells behaved in a manner comparable to the control cells (Fig 5D-5E). We further assessed if there were changes in ISG expression as cells transitioned to being *Adar1^Δ/P195A^*. We did not see any change in expression of the ISGs *Ifit1* or *Irf7* by qPCR, indicating that there was not an activation of the innate immune sensing system in these cells (Fig 5F). We further challenged the *Adar1^Δ/P195A^* cells with transfection with high molecular weight polyinosinic-polycytidylic acid (polyI:C), a synthetic dsRNA, to determine if the P195A mutation altered the response to non-cellular dsRNA. The *Adar1^Δ/P195A^* cells did not have a different response to a dose range of polyI:C compared to control cells when assessing induction of ISGs (Fig 5G). Therefore, at least in an immortalized myeloid cell, the expression of P195A alone is not sufficient to spontaneously activate an innate immune response to cellular dsRNA nor induce a different response to control cells to exogenous dsRNA. As we see activated ISGs *in vivo*, this result suggests that different cell types may respond differently to the P195A mutation as was reported for a biochemical ADAR1 Zα mutant model (Tang *et al*., 2021).

**Figure 5.**
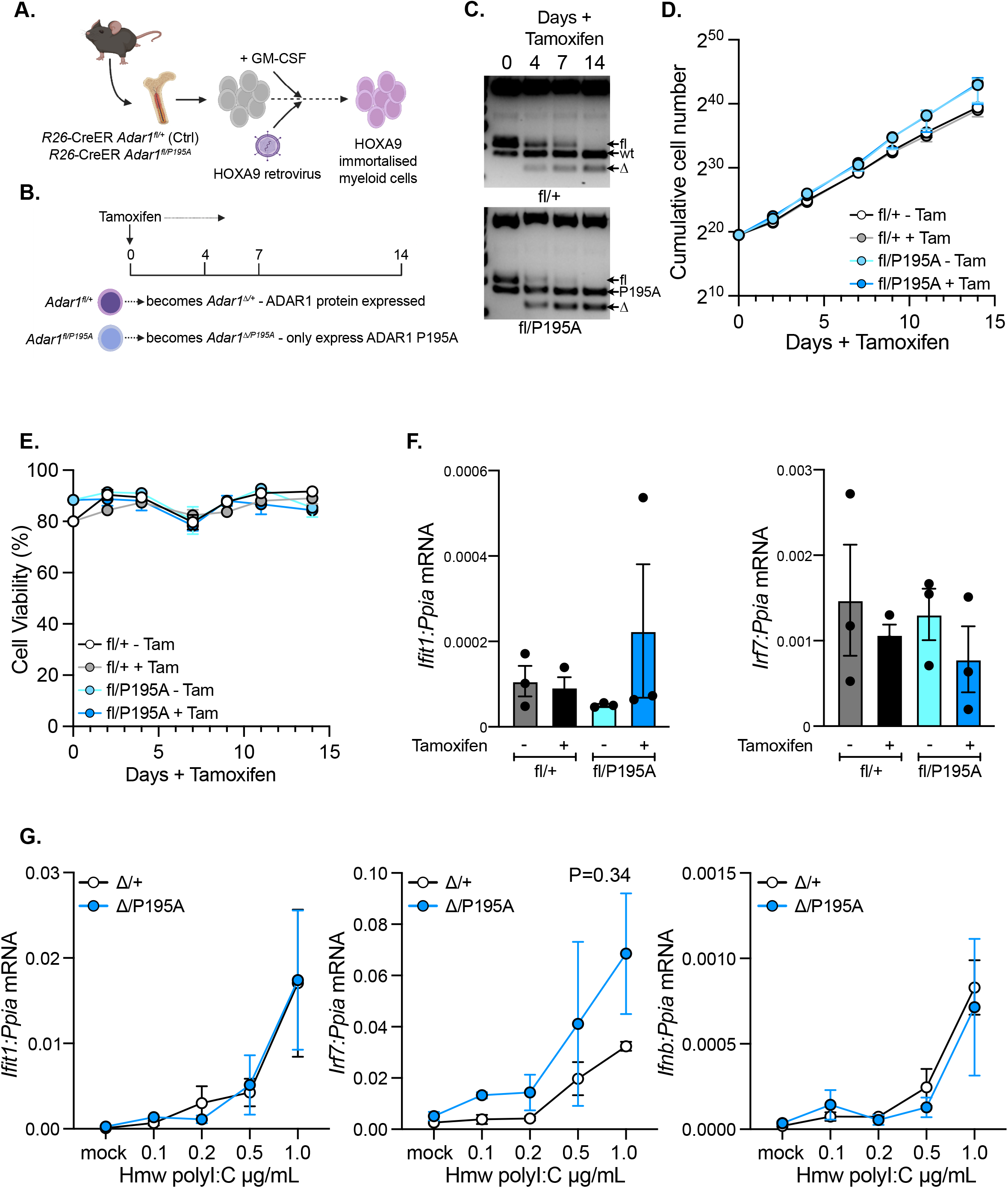
Expression of P195A is well tolerated in myeloid cells. **A)** Schematic outline of how the HOXA9 immortalised myeloid cell lines are derived. **B)** Experimental outline. When tamoxifen is added to the cells this activates deletion of the floxed *Adar1* allele leaving the cells either heterozygous (Controls; *⊗*/+) or P195A only expressing (*⊗*/P195A). **C)** Genomic DNA genotyping demonstrating efficient recombination of the floxed *Adar1* allele following tamoxifen treatment. **D)** Proliferation of cell lines of the indicated genotypes with and without tamoxifen treatment (isogenic pairs). Cells were counted on a Countess II cell counter. **E)** Cell viability of cell lines of the indicated genotypes with and without tamoxifen treatment (isogenic pairs) assessed by trypan blue staining and on a Countess II cell counter. **F)** qPCR (SYBR green) based analysis of *Ifit1* and *Irf7* expression in the cell lines collected at day 14 of analysis. Data expressed as mean +/− SEM gene expression relative to *Ppia* expression. **G)** qPCR (SYBR green) based analysis of *Ifit1*, *Irf7* and *Ifnb* expression in the cell lines transfected with the indicated dose of high molecular weight Polyinosinic-polycytidylic acid (polyI:C). Data expressed as mean +/− SEM gene expression relative to *Ppia* expression. Statistical analysis by t-test using individually calculated AUC per sample. Cell lines had been treated for 14 days with tamoxifen, tamoxifen withdrawn, genotyped, then used for polyI:C dose response. Data from three independently derived cell lines (different donor bone marrow) for each genotype.

### *In vivo* expression of P195A alone activates an ISG signature but is well tolerated

To directly test if there was a difference between the *in vitro* and *in vivo* response to the P195A mutation, we completed an *in vivo* tamoxifen treatment of the *R26*-CreER^T2^ *Adar1^fl/+^* and *R26*-CreER^T2^ *Adar1^fl/P195A^*animals as we previously described (Heraud-Farlow *et al*., 2017). Adult (>8 week old) animals were fed tamoxifen containing diet for 4 weeks (Fig 6A). Analysis of the genomic DNA derived from bone marrow cells at day 28 demonstrated efficient and comparable recombination of the *Adar1* floxed allele (Fig 6B). We did not see evidence for selection against efficient deletion, evidenced by retention of the floxed allele, as we had previously reported with either the *R26*-CreER^T2^ *Adar1^fl/fl^* or *R26*-CreER^T2^ *Adar1^fl/E861A^* in the same experimental model (Heraud-Farlow *et al*., 2017). All animals tolerated the diet well, with no significant difference in weight change between the cohorts compared to the weight at Day 0 (Fig 6C-6D).

**Figure 6.**
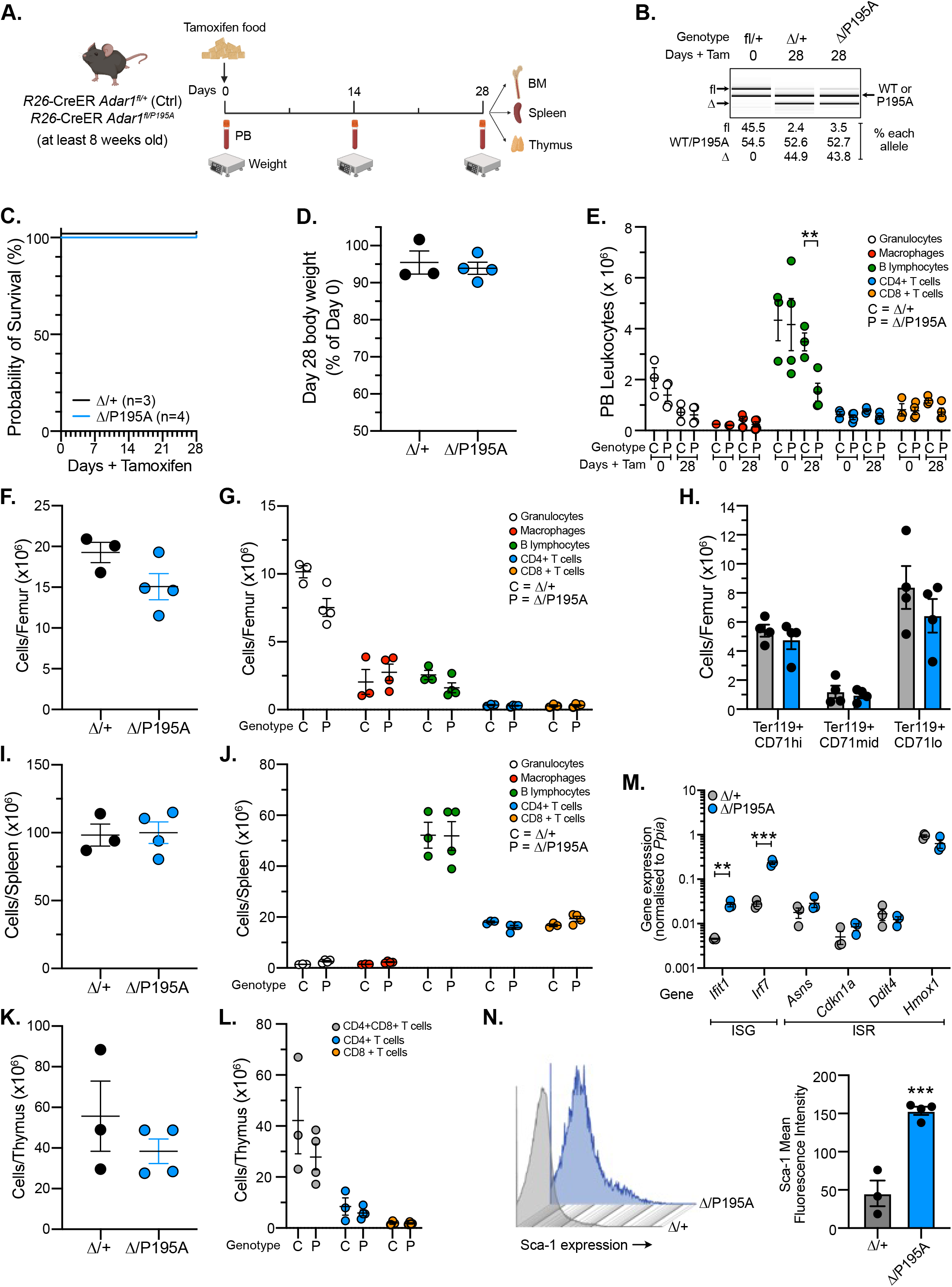
*In vivo* expression of P195A is well tolerated and results in a modest induction of interferon regulated gene expression. **A)** Schematic outline of experiment. **B)** Representative genotyping of recombination of the *Adar1* floxed allele at day 28 using genomic DNA isolated from whole bone marrow cells. Recombination percentage was calculated using LabChip (PerkinElmer) based quantitation of band intensity compared to the WT/P195A allele and known standard/marker. **C)** Survival analysis of mice of the indicated genotypes; number as indicated for each genotype. No statistical difference between genotypes by Log-rank (Mantel-Cox) test using Prism. **D)** Percentage change in body weight of each cohort based on comparison of the weight at day 28 of tamoxifen food compared to day 0 (prior to initiation of tamoxifen containing diet). **E)** Peripheral blood leukocyte populations (by lineage) between genotypes at day 0 and day 28. **F)** Bone marrow cellularity at day 28 between genotypes. **G)** Differential analysis of nucleated cell populations in the bone marrow at day 28. **H)** Erythroid cell populations in the bone marrow at day 28. **I)** Splenic cellularity at day 28 between genotypes. **J)** Differential analysis of nucleated cell populations in the spleen at day 28. **K)** Thymic cellularity at day 28 between genotypes. **L)** Differential analysis of thymocyte populations in the thymus at day 28. **M)** qPCR (SYBR green) based analysis of indicated gene expression in whole bone marrow at day 28. Data expressed as mean +/− SEM gene expression related to *Ppia* expression. **N)** Representative flow cytometry histography of Sca-1 expression between a control (*Δ*/+; grey) and P195A only (*Δ*/P195A; blue) expressing bone marrow sample. Quantitation of Sca-1 mean fluorescence intensity within the lineage negative fraction of whole bone marrow. Unless otherwise stated data expressed as mean +/− SEM; n=3 *R26*-CreER^ki/+^ *Adar1^fl/+^* (control) and n=4 *R26*-CreER^ki/+^ *Adar1^fl/P195A^* (test).

At day 28 of tamoxifen diet, we collected peripheral blood (Fig 6E), bone marrow (Fig 6F-6H), spleen (Fig 6I-6J) and thymus (Fig 6K-6L) and assessed haematopoiesis. We did not see any substantive changes across these organs in terms of cellularity or lineage distribution/differentiation. We also assessed activation of the ISG response by assessing the expression of Sca-1, a known ISG induced in response to a loss of ADAR1, and expression of *Ifit1* and *Irf7* (Essers et al., 2009; Hartner *et al*., 2009; Heraud-Farlow *et al*., 2017) (Fig 6N-6M). We saw increased expression of Sca-1 on the cell surface of the *Adar1^11/P195A^* within the lineage negative fraction of the bone marrow. Prompted by the increased Sca-1 expression, qPCR for the ISGs *Ifit1* and *Irf7* demonstrated a 6-8 fold increased expression of these ISGs in the *Adar1^11/P195A^* animals bone marrow (Fig 6N-6M). We did not find any changes in expression of genes in the integrated stress response pathway in the same samples (Fig 6M). Collectively, these analyses demonstrate that *in vivo Adar1^11/P195A^* animals, but not *in vitro* myeloid cells, have an activation of an innate immune response following somatic restricted expression of P195A. This ISG activation is of a relatively low level, compared to the levels seen when animals and cells are engineered to be ADAR1 null or editing deficient (Heraud-Farlow *et al*., 2017; Liddicoat *et al*., 2016b; Liddicoat *et al*., 2015), and is well tolerated without any effects on the animals overall well-being or survival.

## Discussion

Understanding how disease associated ADAR1 mutations affect the proteins’ function will be critical to the ultimate goal of developing effective treatments for patients with mutations in *ADAR*. To this end, the development of preclinical models that mirror human genetics is an important step. The P193A mutation, which specifically impacts the cytoplasmic ADAR1p150 isoform, is the most commonly reported *ADAR* mutations in humans with AGS and BSN (Livingston *et al*., 2014; Rice *et al*., 2012; Rice *et al*., 2017). In AGS, the P193A mutation is reported as a compound heterozygous mutation with a second mutation, most often one that either compromises expression of the second allele or is predicted to compromise A-to-I editing by the protein product of the second allele. Intriguingly, the P193A mutation is also present in the general human population. As this is the most common human ADAR1 mutation it will be important to understand its effect on both ADAR1’s canonical function in A-to-I RNA editing and in other protein dependent functions of ADAR1.

Herein we describe the independent generation and phenotyping of a murine model of the human P193A mutation. The homologous murine mutation, P195A, is well tolerated and compatible with adult homeostasis when either heterozygous or homozygous. This is consistent with that reported using independently generated P195A knock-in alleles by Maurano et al., (Maurano *et al*., 2021). Whilst homozygosity for P193A is a rare occurrence in the general human population (Karczewski *et al*., 2020) and has not been reported in AGS patients (Rice *et al*., 2012; Rice *et al*., 2017), all groups have established viable and ostensibly normal *Adar1^P195A/P195A^* mice. The present data demonstrate that this mutation is well tolerated and does not significantly compromise ADAR1p150 function *in vivo* under normal laboratory conditions. This proline is not conserved in the Zα family. It is present in the domain wing and affects the kinetics of binding to left-handed nucleic acids that differs between species (Subramani et al., 2016). Interestingly, Zα mutation to highly conserved residues involved in binding Z-DNA (N175A/Y179A) (de Reuver *et al*., 2021; Tang *et al*., 2021) was well tolerated *in vivo* when homozygous, while the W197A mutant (W195 in human), essential to stabilizing the wing, was reported to have a more severe *in vivo* phenotype (Nakahama *et al*., 2021). Collectively, these data indicate that specific mutations within the Zα domain of ADAR1p150, such as the W197A, reveal specific functions for Zα *in vivo* and this warrants further detailed understanding.

The most striking difference between our findings and those reported by Maurano et al., (Maurano *et al*., 2021) are apparent once the P195A allele was crossed to an *Adar1^+/−^* allele. The *Adar1^P195A/−^*genotype more closely approximates that reported in AGS patients, where the P193A allele is reported with a second mutation predicted to be deleterious (Rice *et al*., 2017). Maurano et al., reported that the majority of their *Adar1^P195A/−^* mice were significantly runted and die by 30 days of age, with a single animal surviving to 84 days of age (Maurano *et al*., 2021). In contrast, we see long term (>250 days) survival of the majority of the compound mutant *Adar1^P195A/−^* mice. The basis for this profound difference is not immediately apparent but is important to understand. Whilst differences in vivariums may contribute, it seems unlikely that this is the primary driver of the differences. We believe that two factors may be significant contributors to the differences. Firstly, we observe significant differences in survival and weights of the *Adar1^P195A/−^*mice based on the breeding pair genotype (Fig 3). At present we have results from the breeding of two female and one male *Adar1^P195A/−^*mice crossed to *Adar1^P195A/+^* mice. When an *Adar1^P195A/−^* was used for breeding we see significantly reduced weaning weights and poor survival of the *Adar1^P195A/−^*pups, not dissimilar to that reported by Maurano et al., (Maurano *et al*., 2021). It should be noted that the *Adar1^P195A/−^* genotype is most similar to that of an AGS patient, and to the best of our knowledge there is no available information as to the breeding of these human genotypes. Maurano et al., also tested the effects of the P195A mutation and concurrent deletion of the other ADAR1p150 isoform (*Adar1^P195A/p150-^*) which we have not undertaken (Maurano *et al*., 2021). We did, however, test the effect of the combination of the P195A mutation with the editing dead E861A mutation (Liddicoat *et al*., 2015). The *Adar1^P195A/E861A^* did not have any early lethality and had more consistent weaning weights than the *Adar1^P195A/−^* mice in our cohorts. This demonstrates that A-to-I editing by the non P195A p150 protein is not essential to suppress the observed phenotypes but is required to prevent MDA5 activation, based on ISG expression. Therefore, runting and early lethality of *Adar1^P195A/−^*mice is due to a protein-dependent editing-independent function of ADAR1p150.

A more significant contributor to the dissimilarities is likely to be the different *Adar1* deficient alleles used in the respective studies. Maurano et al, utilised an *Adar1* deficient allele derived by the germ-line deletion of the conditional *Adar1^fl^* allele that deletes exon 7 to 9 (Hartner *et al*., 2004; Hartner *et al*., 2009; Maurano *et al*., 2021; Pestal *et al*., 2015). This allele has recently been demonstrated to yield a truncated, editing deficient and mislocalized protein product impacting both p110 and p150 isoforms (Bajad *et al*., 2020). It is possible that the defective ADAR protein from the exon 7-9 deleted allele could stimulate the integrated stress response reported, as misfolded proteins are known to activate this pathway (Subramani *et al*., 2016). We have used this same genotype in both the immortalised myeloid cells (Fig 4) and somatic deletion models (Fig 5) and in both acute deletion settings we see tolerance of this genotype. Alternatively, the truncated p110 and p150 proteins, whilst non-functional for A-to-I editing, harbour the potential to interfere with the activity of the P195A p150 protein in the cytoplasm and the wild-type p110 expressed from the P195A allele through substrate competition or dimerization (Cho et al., 2003; Valente and Nishikura, 2007). This could potentially result in a less functional, effectively hypomorphic, P195A p150 and WT p110 when heterozygous. In our study we have used an exon 2-13 deletion of *Adar1*, demonstrated to be a null allele and not known to express any protein product (Hartner *et al*., 2004). The data strongly argue that the runting and completely penetrant post-natal lethality of the *Adar1^P195A/−^* mice reported by Maurano et al., is most likely confounded by the effects of the *Adar1^ΔEx7-9^* allele (Maurano *et al*., 2021). When crossed to a true null allele, the majority of *Adar1^P195A/−^*mice survive with no pathological differences in adult animals compared to controls. There may also be other strain differences such as those immune deficiencies previously reported in some lines of C57BL/6 mice (Koehler et al., 2020) or accounting for the high birth weight in the Maurano et al. study. The ultimate cause of death of the runted *Adar1^P195A/−^* mice from our cohorts remains to be determined. Moreover, the P195A mutation can be complemented by the provision of the editing deficient E861A allele. This demonstrates that provision of a full length, editing deficient p150 protein is sufficient to prevent the early lethality and runting we see in a subset of our *Adar1^P195A/−^* cohort that depends on parental genotype. In summary the P195A mutation, homologous to the most common human ADAR1 mutation reported in AGS, is well tolerated *in vivo* and the loss of MDA5 is sufficient to completely rescue the *Adar1^P195A/−^* mice.

The physiologically most important role of A-to-I editing by ADAR1 is to edit cellular/endogenous dsRNA to prevent MDA5 mediated innate immune sensing (Liddicoat et al., 2016a; Liddicoat *et al*., 2015). This has been demonstrated in both mouse models and human cells (Liddicoat *et al*., 2015; Mannion *et al*., 2014; Pestal *et al*., 2015). The P195A mutation is associated with reduced editing of some substrates *in cellulo* (Maurano *et al*., 2021). In both human and mouse, the loss of MDA5-MAVS prevents innate immune activation following the loss of ADAR1 or loss of A-to-I editing by ADAR1. Previous work has not demonstrated rescue from embryonic lethality of *Adar1^−/−^* animals by concurrent loss of PKR (*Eif2ak2^−/−^*) (Wang *et al*., 2004) or by loss of IFNAR (Liddicoat *et al*., 2016b; Mannion *et al*., 2014), unlike the rescue to birth or adulthood by loss of MDA5 (*Ifih1^−/−^*) or MAVS (*Mavs^−/−^*) of the *Adar1^−/−^* or ADAR1 editing deficient (*Adar1^E861A/E861A^*) mice, respectively (Heraud-Farlow *et al*., 2017; Liddicoat *et al*., 2015; Pestal *et al*., 2015). Based on the ISR signature and genetic crosses it was proposed that PKR activation was important in the pathology of the P195A mouse models generated by Maurano et al., and this was genetically tested (Maurano *et al*., 2021). We see a modest activation of the downstream transcriptional markers of MDA5 signalling (interferon stimulated genes) in tissues isolated from ∼normal weight adult *Adar1^P195A/−^* and *Adar1^P195A/E861A^* mice. In the brains of runted *Adar1^P195A/−^* animals there was an even higher level of induction of ISGs, but even in these severely runted animals, we did not see evidence for activation of the PKR-related integrated stress response. We also assessed levels of phospho-eIF2a, a marker of PKR activation, in the kidney of the runted *Adar1^P195A/−^* animals and did not see any difference to normal sized *Adar1^P195A/−^* and control animals (Supp Fig 3D). Most recently it has been proposed by the same group that the pathology of the *Adar1^P195A/−^* animals is due to activation of an alternative protein, ZBP-1 (Hubbard et al., 2022). ZBP-1 is the only other mammalian Zα domain containing protein and has been linked to aspects of ADAR1 biology (Zhang et al., 2022). It is not at present clear how this reconciles with the prior model where activation of PKR and the integrated stress response was proposed as the primary driver of the phenotypes in their *Adar1^P195A/−^* model (Hubbard *et al*., 2022; Maurano *et al*., 2021). We see complete rescue of the *Adar1^P195A/−^* and *Adar1^P195A/E861A^*mice by loss of MDA5 with a full suppression of ISG expression to baseline levels *in vivo*, consistent with a model where MDA5 is the primary, and initiating, sensor of unedited self dsRNA. We do not see evidence in the adult mouse tissues of transcriptional changes consistent with activation of PKR as was reported by Maurano et al., in the *Adar1^P195A/p150-^* animals (Maurano *et al*., 2021). Interestingly there is a dosage dependent effect of MDA5 with a degree of genetic rescue afforded by being *Ifih1^+/−^* (Supplemental Fig 2B). All of the ADAR1 Zα mutants described to date demonstrate complete rescue by loss of either MDA5 (Maurano *et al*., 2021; Nakahama *et al*., 2021) or MAVS (de Reuver *et al*., 2021; Tang *et al*., 2021). Collectively, these results demonstrate that the absence of MDA5, and subsequent innate immune response, is sufficient to rescue the effects of ADAR1 P195A Zα mutants. In summary the P195A mutation, homologous to the most common human ADAR1 mutation reported in AGS, is well tolerated *in vivo* and the loss of MDA5 is sufficient to completely rescue the *Adar1^P195A/−^* mice. These findings are consistent with a role for p150 in down-regulating the ISG response but do not explain the pathology observed in AGS patients.

## Acknowledgements

The authors would like to thank A Herbert for critical discussion and comment; E Tonkin for technical assistance; the Monash Genome Modification Platform (MGMP) at Monash University for the generation of the P195A mice; Monash Antibody Technology Facility (MATF) for purification of ADAR1 antibody from hybridomas; the Phenomics Australia Histopathology and Slide Scanning Service, University of Melbourne for histopathology on samples; and St. Vincent’s Hospital BioResource’s Centre for care of experimental animals. We thank M Kamps (UC San Diego) for HOXA9 plasmids and Addgene for plasmid distribution. Schematic figures were made using BioRender.com.

The P195A mutant mice were produced via CRISPR/Cas9 mediated genome editing by the Monash Genome Modification Platform (MGMP), Monash University as a node of Phenomics Australia. This study utilised the Phenomics Australia Histopathology and Slide Scanning Service, University of Melbourne. Phenomics Australia is supported by the Australian Government Department of Education through the National Collaborative Research Infrastructure Strategy, the Super Science Initiative, and the Collaborative Research Infrastructure Scheme. This work was supported by the National Health and Medical Research Council (NHMRC; APP1183553 to C.R.W and J.H.F; APP1182453 to J.H.F); a Melbourne Research Scholarship (to Z.L. from The University of Melbourne); J.H.F is supported by a fellowship from 5point Foundation; and in part by the Victorian State Government Operational Infrastructure Support Scheme to St Vincent’s Institute.

The funders had no role in study design, data collection, and analysis, decision to publish, or preparation of the manuscript.

## Author Contribution Statement

J.H-F and C.R.W conceptualized the study. Z.L, J.H-F and C.R.W designed the experiments. Z.L, S.T, A.G, J.H-F and C.R.W performed the experiments. Z.L, J.H-F and C.R.W wrote the original manuscript, and all authors reviewed and edited the manuscript; J.H-F and C.R.W were responsible for funding acquisition. J.H.F and C.R.W provided supervision.

## Supplemental Information

**Supplemental Figure 1.**
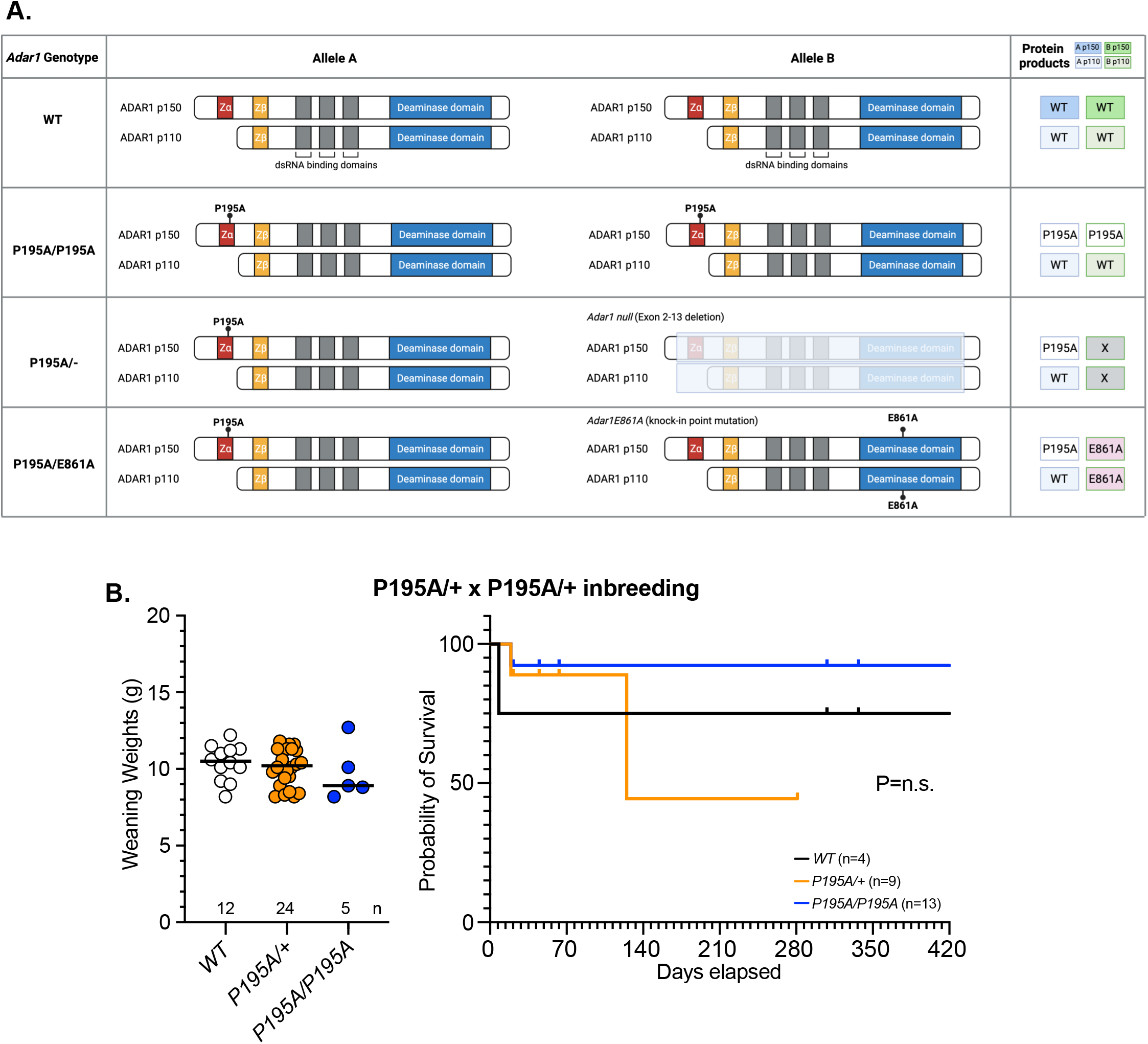
Schematic outline of the different alleles used in this study and weaning weights and survival of pups from *Adar1^P195A/+^* inbreeding. **A)** Outline of the different alleles used and expected consequences on ADAR1 p110 and p150 expression/ function. **B)** Weaning weights and survival analysis of pups derived from inbreeding of *Adar1^P195A/+^*animals; number as indicated for each genotype. For weights: Ordinary one-way ANOVA with Tukey’s multiple comparison test (adjusted P value); survival using Log-rank (Mantel-Cox) test - no significant difference when tested. Data displayed as individual animals with mean indicated. Weaning weights not available if animal was found dead prior to weaning; all animals where genotype was confirmed are included in the survival analysis (where possible any found dead prior to weaning were genotyped post-mortem).

**Supplementary Figure 2.**
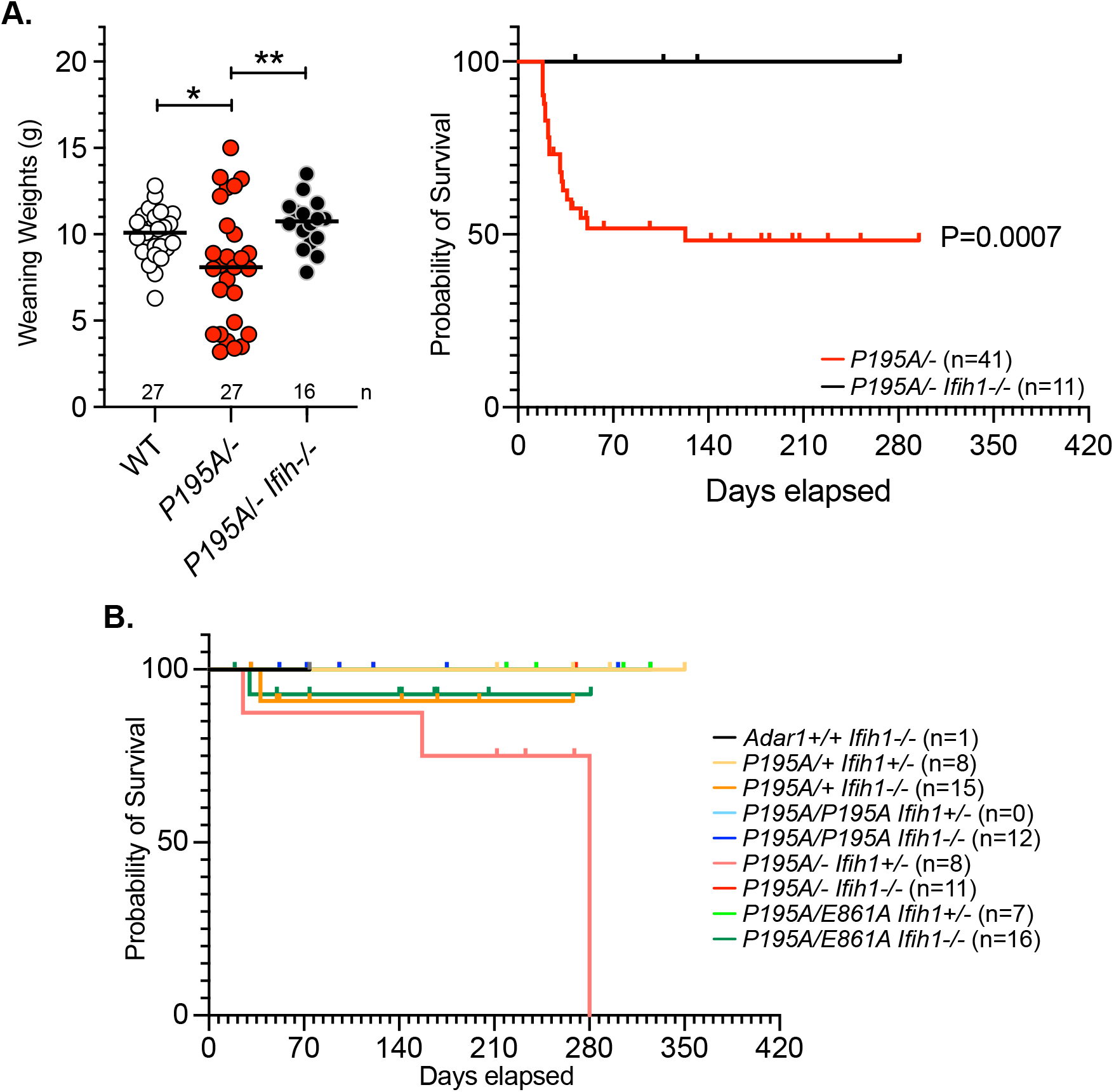
Loss of MDA5 prevents *Adar1^P195A/−^* lethality. **A)** Comparison of survival and weaning weights of *Adar1^P195A/−^* and *Adar1^P195A/−^ Ifih1^−/−^* (datasets replotted from Fig 2C-2F). For survival analysis: *P<0.05 (Log-rank (Mantel-Cox) test); For weaning weights: *P<0.05; **P<0.01 (One-way ANOVA with Tukey’s multiple comparison test; adjusted P Value). **B)** Comparison of survival of all genotypes on an MDA5 heterozygous (*Ifih1^+/−^*) and homozygous null (*Ifih1^−/−^*) background; complete dataset for cohorts presented in Figure 2E. n as indicated in legends.

**Supplemental Figure 3.**
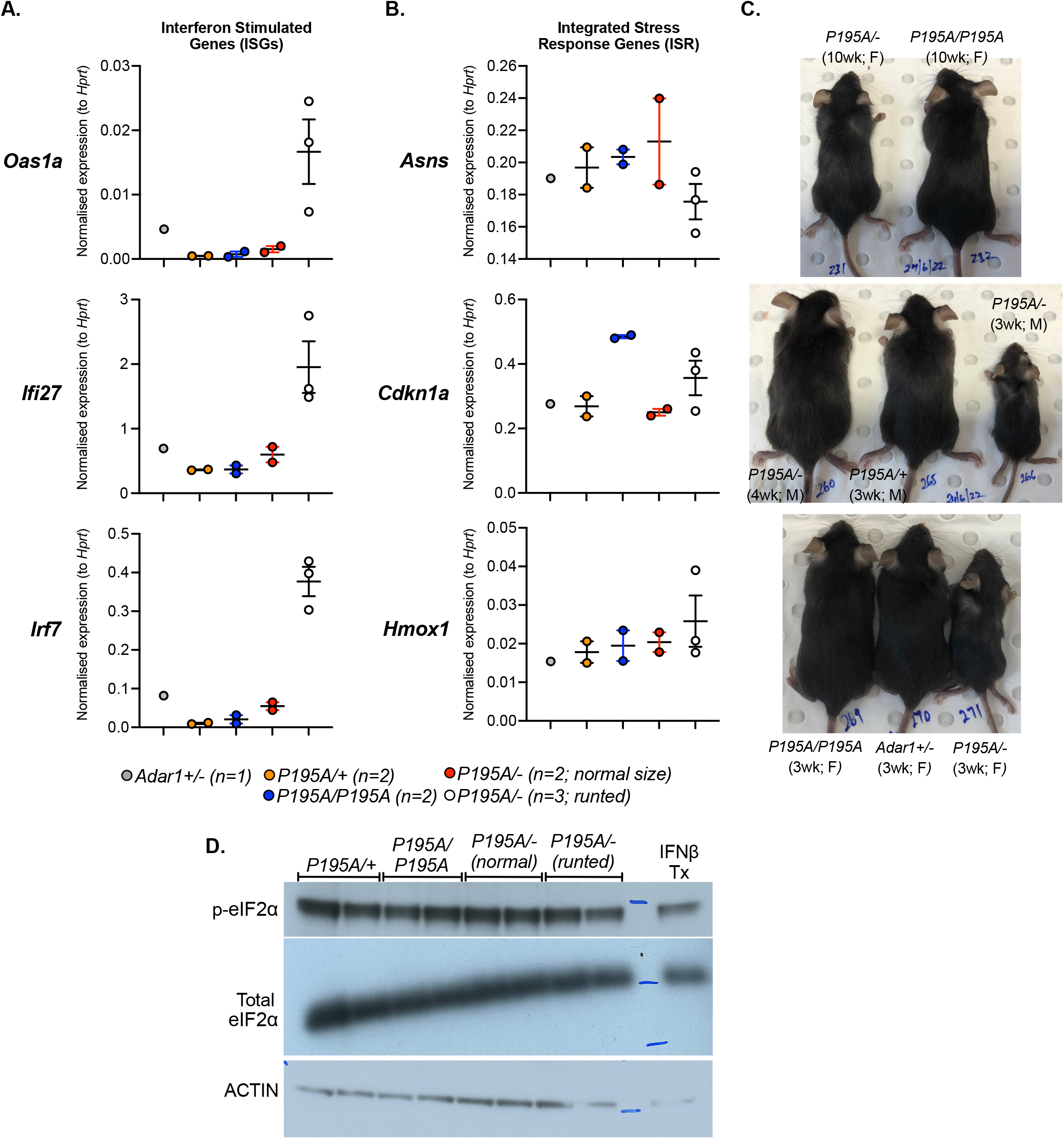
Analysis of runted *Adar1^P195A/−^* compared to normal sized *Adar1^P195A/−^*. **A)** Taqman based qPCR for the interferon stimulated genes *Oas1a*, *Ifi27* and *Irf7* in brain of mice of the indicated genotypes; number of independently derived samples as indicated for each genotype. **B)** Taqman based qPCR for the integrated stress response genes *Asns*, *Cdkn1a* and *Hmox1* in in brain of mice of the indicated genotypes; number of independently derived samples as indicated for each genotype. Data expressed as mean +/− SEM gene expression relative to *Hprt* expression. **C)** Photographs of the animals used for qPCR in panel A and B with indicated age, sex (F= female; M= male) and genotype as indicated. **D)** Western blot of phospho-eIF2α and total eIF2α levels in whole kidney lysates derived from the indicated genotypes; same mice as used for Panels A-C. Positive control is IFNβ treated MEF cell. Data related to Figure 4C and Figure 4D. Data in Figure 4D (heatmap representation of qPCR data from the brain) were derived from data in panel A and B.

## Datasets

**Supplementary Dataset 1:** Full histopathology report of adult *Adar1^P195A/E861A^* and control mice.

**Supplementary Dataset 2:** Full histopathology report of brain, kidney and liver samples from adult *Adar1^P195A/−^* and control mice.

---

**Supplementary Table 1.**
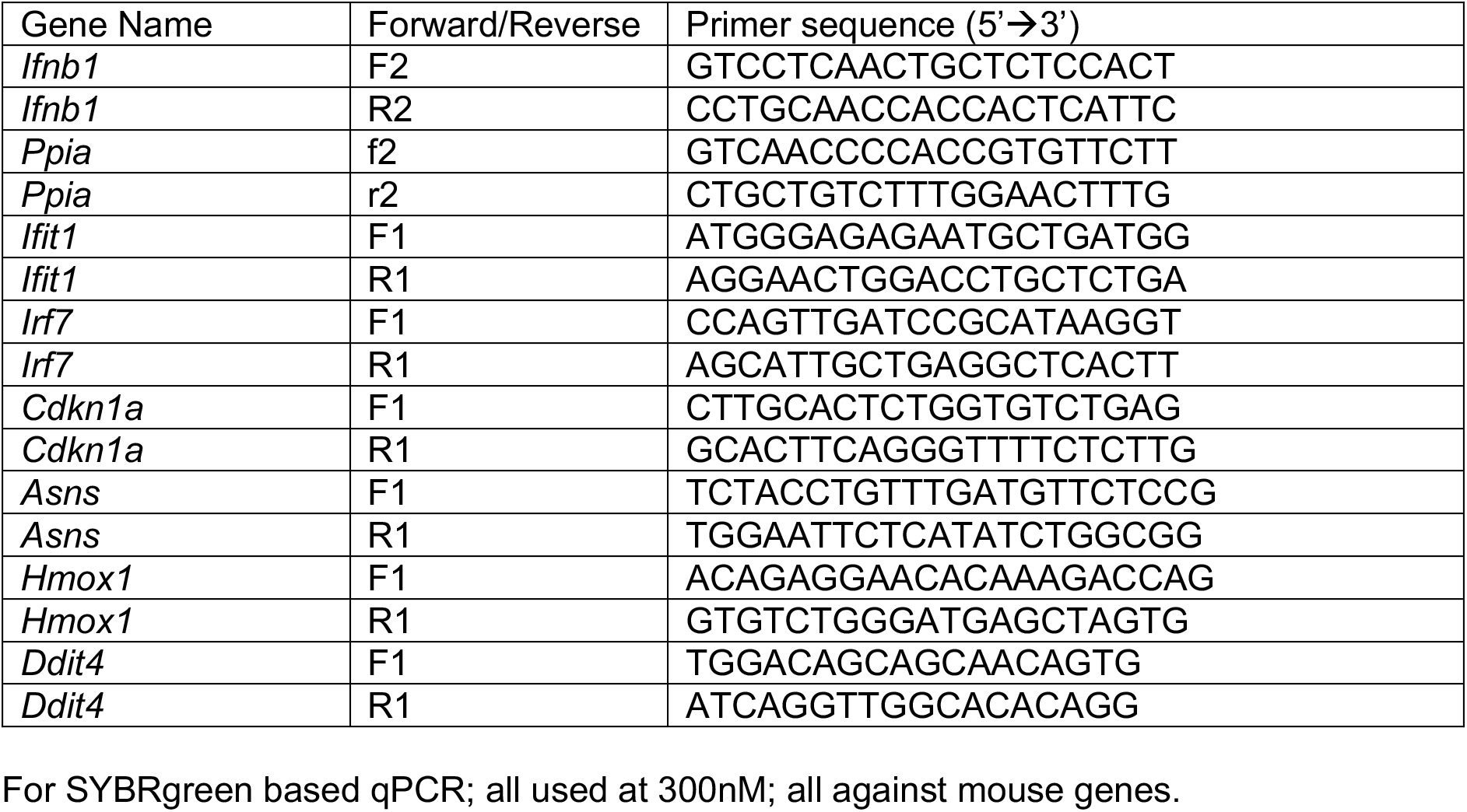
qPCR primer sequences for SYBRgreen based qPCR.

**Supplementary Table 2.**
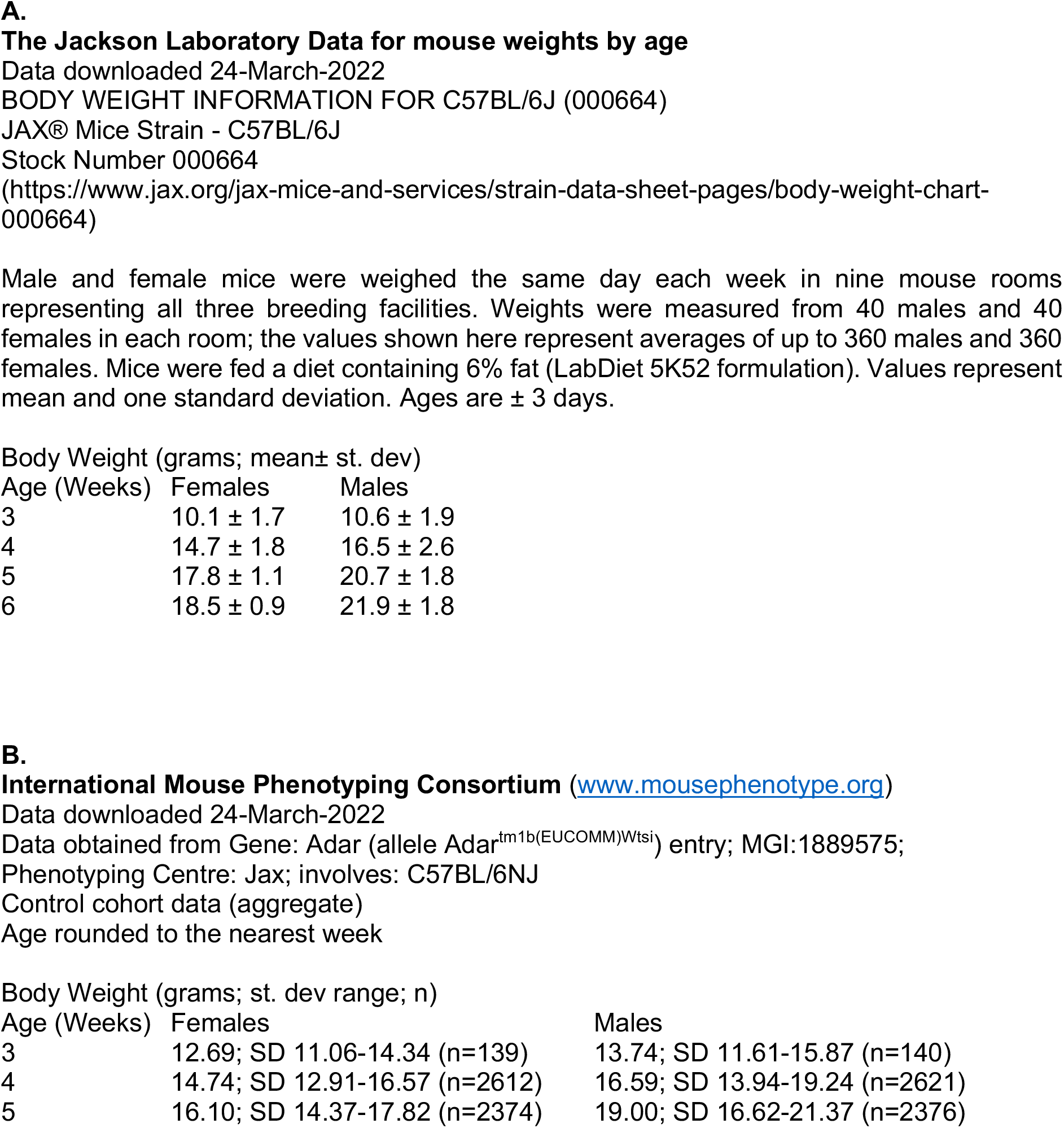
Average mouse weights by age for C57Bl/6 background mice.

## 9.1 Histopathology Report

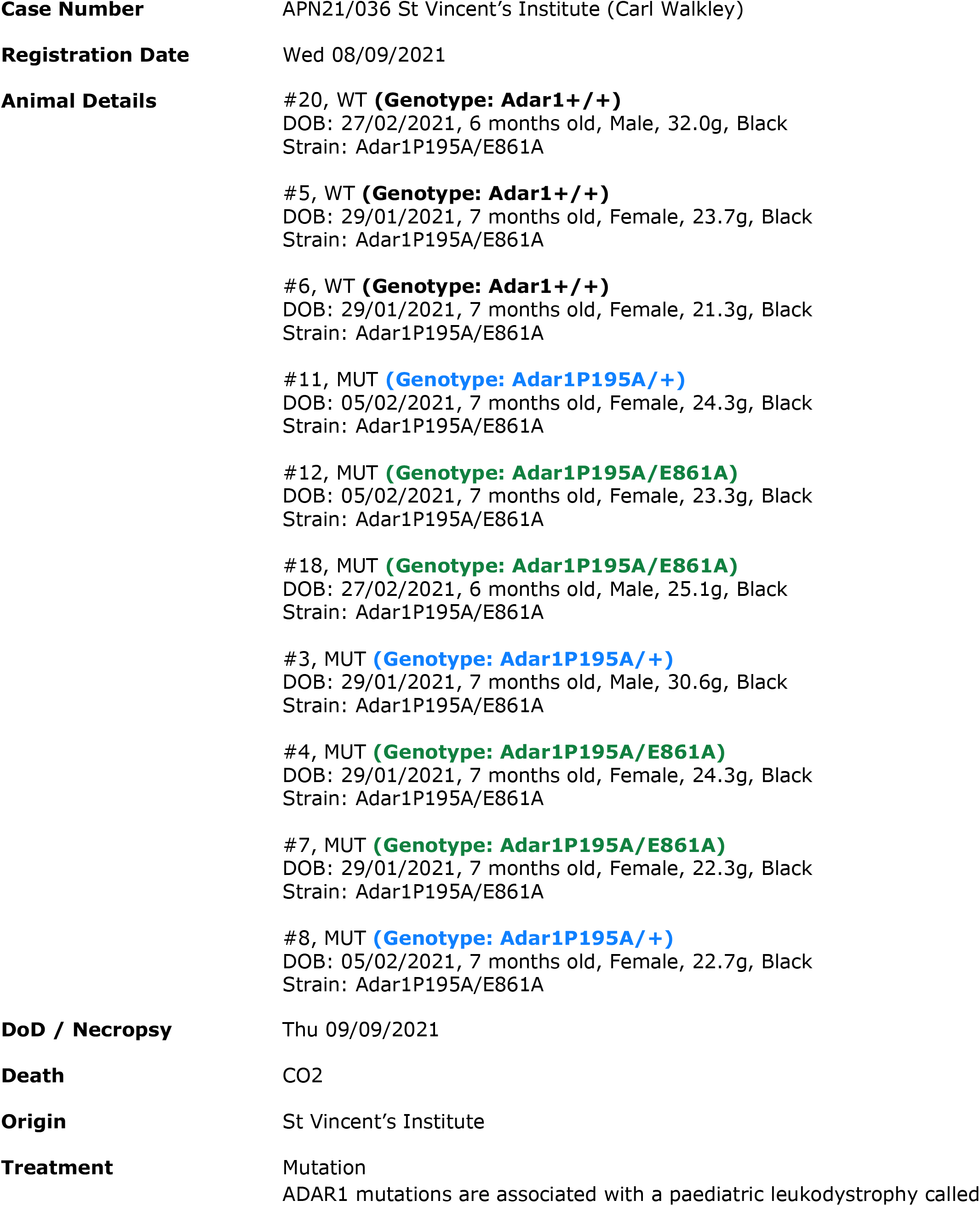
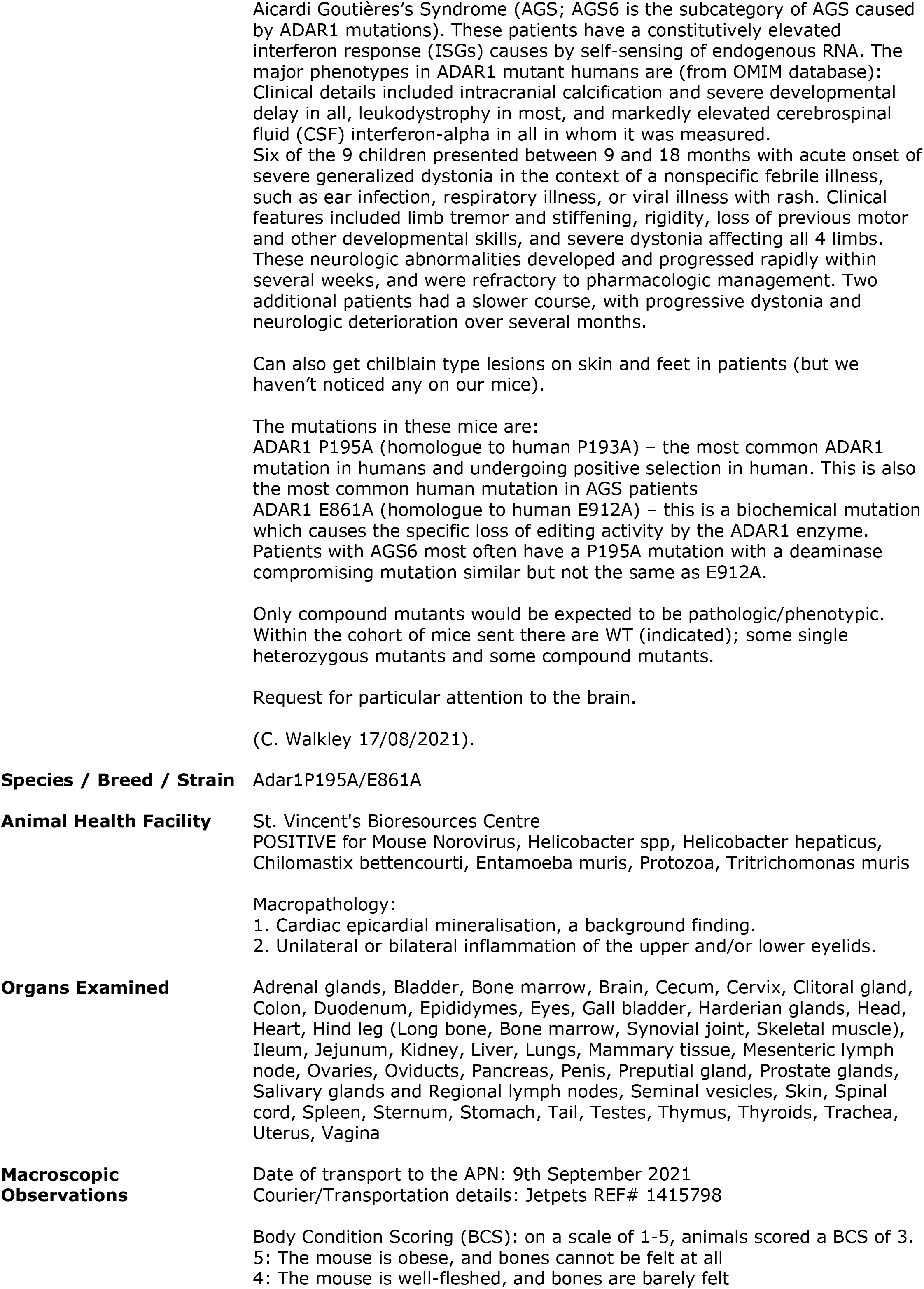
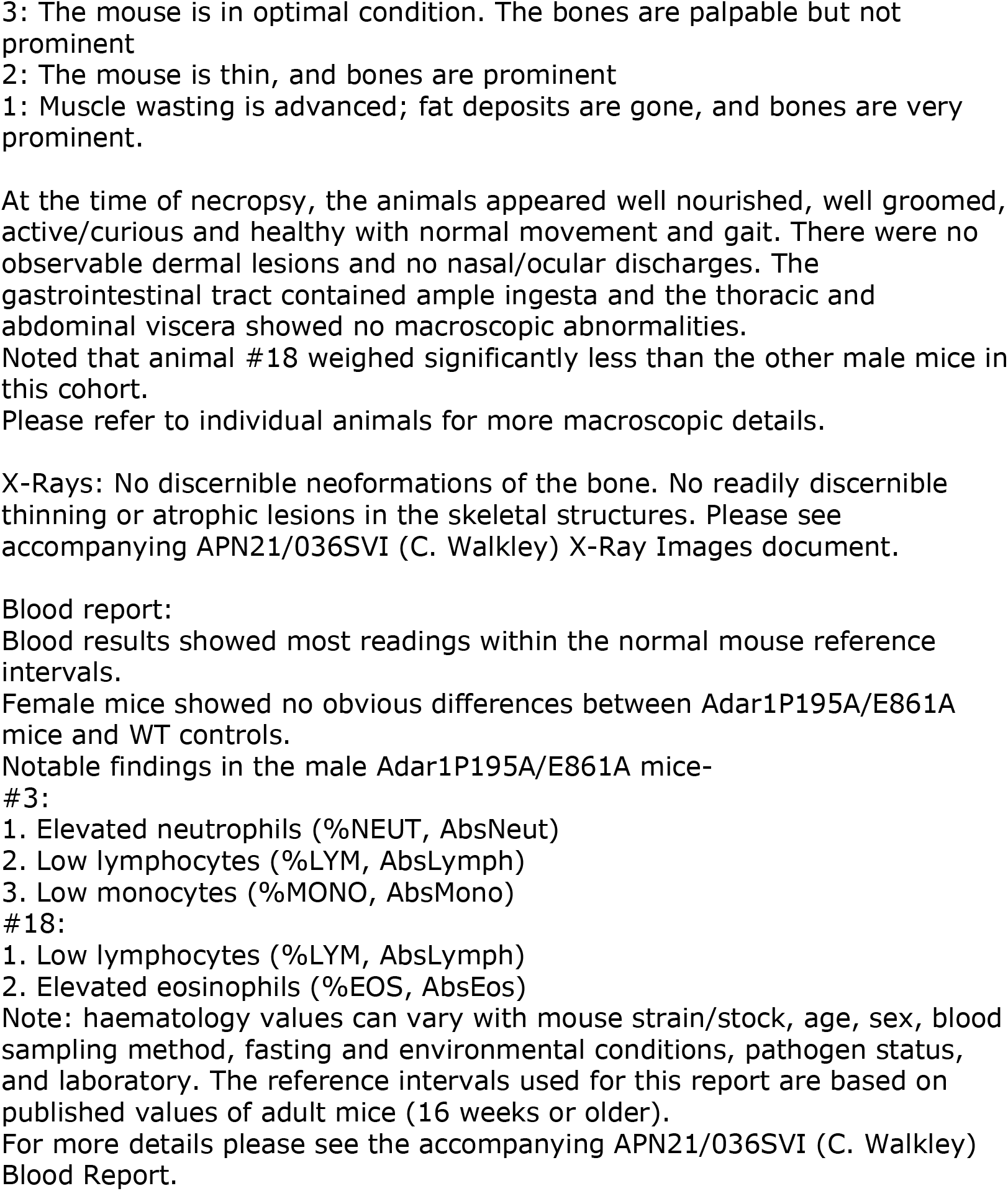

### Microscopic Observations

#### SUMMARY

Both the Adar1P195A/E861A mutant animals and the WT controls of this cohort showed features indicative of increased cerebrospinal fluid in the lateral ventricles of the brain (hydrocephalus). These features were seemingly more prominent in the Adar1P195A/E861A mutants.

Note: hydrocephalus is a background lesion seen in some inbred mouse strains.

Various tissues showed multiple other incidental and/or background lesions - no obvious micromorphological differences were identified between the Adar1P195A/E861A mutant animals and WT controls of this cohort.

Please see individual animals for more micromorphological details.

Notes:

1. Reactive lymph nodes are defined as mild follicular hyperplasia, germinal centre formation and occasional sinus histiocytosis - a common finding in mice.
2. Hyperplastic lymph node follicles are identified by an increase in number and size of follicles and conversion to secondary follicles. Hyperplasia of the paracortex is characterized by an increase in the cell density and, depending on the degree of hyperplasia, an increase in the paracortical area.
3. Accumulation of leukocytes and other cells are common nonneoplastic lesions in many tissues. The term “inflammation” is used when the cell (leukocyte) accumulations are part of an active inflammatory process (typified by concurrent features such as vascular changes, necrosis, fibrosis, and/or tissue disruption). In contrast, cell “infiltration” is used when the cell (e.g., lymphocyte) accumulations are present in tissue without other disruption or pathology.
4. Focal inflammatory cell aggregates consisting of mononuclear, polymorphonuclear, and/or histiocytic cells are frequently observed in ageing mice (Maranport RR. 1999. Pathology of the Mouse.). These can be present as lymphoid aggregates found in various tissues including the renal pelvis, bladder, lungs, liver (Pettan-Brewer C and Treuting PM. 2011. Practical pathology of aging mice.) and salivary glands (Haines DC, Chattopadhyay S and Ward JM. 2001. Pathology of Aging B6;129 Mice).
5. Mild extramedullary haematopoiesis (EMH) in the red pulp of the spleen is a common finding in the mouse. EMH consists of erythroid precursors, myeloid precursors, megakaryocytes or all three. While some degree of extramedullary haematopoiesis is present in normal rodents, especially in mice, increased extramedullary haematopoiesis can result from haematotoxin insult, systemic anaemia, and infections elsewhere in the body. (Suttie AW. 2006. Histopathology of the spleen).
6. The appearance of pigment in the red pulp is a common background lesion in rodents (Suttie AW. 2006. Histopathology of the spleen).
7. Thymic cysts in the rodent represent either a dilatation of thymic tubular structures or remnants of the thymopharyngeal duct. They are common findings in the involuted and/or atrophied thymus glands of rats and mice. Thymic cyst formation becomes more prominent with age and is associated with involution. (Hobbie K, Elmore SA and Kolenda-Roberts HM. 2015. Thymus – Cyst. In: National Toxicology Program Nonneoplastic Lesion Atlas).
8. Cytoplasmic inclusions of homogeneous eosinophilic hyaline-like material may be seen in older mice in intrahepatic biliary epithelial cells, as well as epithelial cells in the gallbladder. In marked cases, there is hyperplasia of the glandular epithelium, and crystalline forms of the eosinophilic inclusion material may be present both intracellularly and extracellularly (Maronpot RR. 1999. Liver and gallbladder. In: Pathology of the Mouse: Reference and Atlas).
9. Focal fatty change of the liver can be a spontaneous lesion and may be more common in some strains than others (Maronpot RR. 2014. Liver-Fatty Change. In: National Toxicology Program Nonneoplastic Lesion Atlas).
10. Fatty change of the liver may occur in mice as a response to a toxicant. It is also seen in old obese controls and is more common in male than in female mice. The degree of fatty metamorphosis may vary and usually starts with a centrilobular distribution (Frith CH, Ward JM. 1988. Digestive System. In: Color Atlas of Neolastic and Non-neoplastic Lesions in Aging Mice).
11. Cystic endometrial hyperplasia, occurs more commonly in mice than in rats and is particularly associated with increasing age. (Willson G, Cimon KY. 2015. Uterus, Endometrium – Hyperplasia, Cystic. In: National Toxicology Program Nonneoplastic Lesion Atlas)
12. Focal pancreatic infiltrates of lymphocytes, plasma cells, and macrophages are uncommon in aged mice. The infiltrates are generally minimal and may be associated with atrophy (Maronpot RR. 1999. Exocrine and Endocrine Pancreas. In: Pathology of the Mouse: Reference and Atlas).
13. Yellow to brown pigment is often seen as an aging change in mouse and rat ovaries. Ceroid is the pigment most frequently seen, and this accumulates mainly in the cytoplasm of interstitial cells.Golden brown to darker brown pigment is sometimes seen in the uterus. Hemosiderin may be seen secondary to haemorrhage. Other types of pigment may also be seen, such as lipofuscin or ceroid. (Willson G, Cimon KY. 2015. Uterus, Endometrium – Hyperplasia, Cystic. In: National Toxicology Program Nonneoplastic Lesion Atlas).
14. The early histopathologic changes of nephropathy are characterised by the presence of focal to multifocal cortical tubules which have slight cytoplasmic basophilia and nuclear crowding. The basement membrane of these tubules are often variably thickened….As the disease progresses more affected tubules become evident and occasional eosinophilic proteinaceous tubular casts are noted (Maronpot RR. 1999. Kidney – Nephropathy. In: Pathology of the Mouse).
15. Lymphocyte apoptosis is characterized by cell shrinkage, nuclear pyknosis and fragmentation with apoptotic bodies. This type of cell death normally occurs within the germinal centres of secondary follicles where it is an important homeostatic mechanism (Hobbie K, Elmore SA and Kolenda-Roberts HM. 2015. Lymph Node – Apoptosis, Lymphocyte. In: National Toxicology Program Nonneoplastic Lesion Atlas).
16. Valvular myxomatous changes or degeneration can be an age-related spontaneous or chemical-induced change. The lesion ischaracterized by focal or segmental thickening of the subendocardium in the valve leaflets andexpansion of the spongiosa of the valve leaflet with extracellular fibromyxoid material composedpredominantly of glycosaminoglycans.Occasionally, fibrin deposits or thrombi and collections of neutrophils or mononuclear cells are seen (Johnson CL, Nyska A. 2017. Heart, Valve, – Degeneration. In: National Toxicology Program Nonneoplastic Lesion Atlas).
17. A number of intestinal parasites may be seen within the lumen of the large intestine, the ceacum, and less commonly the small intestine. The significance and presence of protozoan organisms is questionable since infected animals are normally asymptomatic. The presence of Protozoa such as flagellates, ciliatesm etc. have not been associated with any microscopic lesions of clinical significance in mice (Maronpot RR. 1999. Intestines and Mesentry: Small and Large Intestine. In: Pathology of the Mouse: Reference and Atlas).
18. Necrosis of the exocrine pancreas, a naturally occurring lesion in mice usually only involving a few acini. The spontaneous lesion is both uncommon and very limited within the pancreas (Maronpot RR. 1999. Exocrine and Endocrine Pancreas. In: Pathology of the Mouse: Reference and Atlas).
19. Epidermal ulcersare among the most common spontaneous findings in mice and rats but may also be associated with dermal application of test agents.Ulcers are characterized by segmental or more extensive loss of the epidermis, including the basement membrane, with exposure of the underlying dermis. Erosion is characterized by the partial loss of the epithelium, with the basement membrane left intact. (Boyle MH, Hill GD. 2014. Skin - Ulcer and Erosion. NTP Nonneoplastic Lesion Atlas).
20. Focal hepatic necrosis is a non-specific entity quite often encountered as an incidental finding in the liver of mice. It can be the result of viruses (mouse hepatitis), bacteria (Clostridium piliforme), toxicants, and ischemia while the etiology is often unknown. It may involve single cells, single or multiple lobules, and it may vary in distribution. Coagulation necrosis with distinct eosinophilic cytoplasm and pyknotic or absent nuclei is the typical morphologic feature (Frith CH, Ward JM. 1988. Color Atlas of Neolastic and Non-neoplastic Lesions in Aging Mice).
21. Ovarian cysts are a common finding in rats and mice. Ovarian cysts may be unilateral or bilateral, single or multiple, and may become quite large… size and number of cysts increase with age. (Willson G, Cimon KY. 2015. Ovary - Cyst. In: National Toxicology Program Nonneoplastic Lesion Atlas).
22. Hydrocephalus may be communicating or noncommunicating; that is, the former has no apparent obstructive process, whereas the latter has an obstructive cause somewhere in the ventricular connections. Most commonly, communicating hydrocephalus is considered to result from an idiopathic increase in cerebrospinal fluid production or deceased resorption. (Little P, Rao DB. 2014. Brain – Hydrocephalus. In: National Toxicology Program Nonneoplastic Lesion Atlas).
23. Germ cell degeneration of the testes is a nonspecific term that generally includes a number of degenerative features, such as tubular vacuolation, partial depletion of germ cells, degenerating (multinucleated or apoptotic) germ cells, and disordered arrangement of the germ cell layers. Chemically induced germ cell degeneration can be multifocal in distribution, but it is most often a bilateral lesion that affects most of the seminiferous tubules to varying degrees. It can also be an incidental background finding in rats and mice of any age, but the incidence increases with age. (Wilson G and Cimon K Y. 2015. Testis, Germ cell – Degeneration. In: National Toxicology Program Nonneoplastic Lesion Atlas).
24. Small focal clusters of inflammatory cells in the interstitium are common incidental findings in the Harderian gland of rats and mice. These infiltrates are most commonly mononuclear cells (lymphocytes), but other inflammatory cells may also be present (Gruebbel MM. 2014. Harderian Gland – Infiltration Cellular, Mononuclear Cell. In: National Toxicology Program Nonneoplastic Lesion Atlas).
25. Exocrine acinar atrophy is the most common spontaneous degenerative change in the pancreas of both rats and mice. Acinar atrophy may range from focal atrophic exocrine acini with no inflammation or fibrosis to diffuse atrophy of exocrine acini with replacement by adipose tissue and residual ducts, vasculature and islets of Langerhans. Acinar atrophy frequently represents the sequel of chronic inflammation, and as such, is often accompanied by infiltrates of mononuclear cells and fibrosis. (Nolte T, Brander-Weber P and Dangler C. 2016. Pancreas (Exocrine) – Atrophy, acinar cell. In: Nonproliferative and Proliferative Lesions of the Gastrointestinal Tract, Pancreas and Salivary Glands of the Rat and Mouse. J Toxicol Pathol 2016; 29 (1 Suppl): 1S–124S).

#### #20 (control)

Macro Observations

Tail suspension test for neurological defects - negative.
Dentition, tongue and oral cavity was unremarkable.
BCS: 3
Testes: 8×5×4mm, symmetrical
Spleen: 15×5×3mm
Kidneys: 13×7×6mm, symmetrical
Thymus: 7×5×2mm
Lungs inflated.
Heart: 10×8×7mm
Brain: 15×11×5mm, symmetrical
Pituitary gland identified, macroscopically normal
Tail length: 83mm (straight)
Head harvested for evaluation of auditory and vestibular structures.
Bone marrow smear taken from left hind leg.
No macroscopic lesions identified.
Excessive adipose tissue: abdominal viscera (+).

Micro Observations

Animal #20 was used as a male histological control for this cohort.
Marrow smear: Examination of the smear showed representative cells from the myeloid and erythroid series. Occasional cells from the lymphoid series. Occasional and unremarkable megakaryoblasts. (85195)
Peripheral blood smear: Examination of the smear showed red blood cells (majority of cells shown), occasional white blood cells including lymphocytes, segmented neutrophils, monocytes and platelets (clumps). No discernible morphological changes or detectable parasites. (85196)
Organs examined: Testes/Epididymes (85198), Seminal vesicles (85199, 85200), Prostate glands (85199, 85200), Penis/Preputial gland (85201), Urinary Bladder (85199, 85200), Liver/Gall Bladder (85202), Stomach (85203), Duodenum/jejunum/ileum/GALT (85203, 85204, 85205), Cecum/Colon/GALT (85205), Mesenteric Lymph Node (85197), Spleen (85199, 85200), Pancreas (85199, 85200), Kidneys/Adrenal Glands (85206), Salivary Glands/regional lymph nodes (85197), Thyroids (85208), Trachea/Lungs (85207, 85208), Thymus (85207, 85208), Heart (85207, 85208), Skin (85209), Tail (85279, 85280, 85281), Eyes/Harderian Glands (85210), Brain (85211), Spinal cord (85283, 85284, 85285), Hind leg (85279, 85280, 85281), Head (85312, 85313, 85314), Sternum (85282).
Micromorphological changes- Testes: few tubules showing tubular degeneration/atrophy (85198). Liver: multiple small foci of hepatocyte cell loss/necrosis and associated leucocyte infiltrates (85202). Liver: minimal multifocal perivascular mononuclear cell infiltrates (85202). Gall bladder: epithelial hyperplasia, cytoplasmic hyaline droplet accumulation, and neutrophilic infiltrates in the lamina propria (85202). Large intestine: increased mononuclear cell infiltrates in the lamina propria of the colon (85205). Pancreas: focal necrosis and associated leukocyte infiltrates (85203). Pancreas: minimal multifocal perivascular mononuclear cell infiltrates (85199, 85200). Kidneys: minimal multifocal perivascular/peripelvic mononuclear cell infiltrates (85206). Kidneys: single granular cast-like structure in cortex (85206). Salivary glands: mild multifocal perivascular mononuclear cell infiltrates, also seen in the associated lacrimal gland (85197). Lung: minimal multifocal perivascular mononuclear cell infiltrates (85207, 85208). Thymus: few small sized cysts (85208). Heart: few foci of cardiomyocyte vacuolation, indicative of degeneration (85207, 85208). Eye: query thinning/loss of the corneal epithelial cell layer, unilateral (85210). No other indicators of inflammation or damage. Brain: some flattening/stretching of the ependymal cells in the lateral ventricles (85211), indicative of increased cerebrospinal fluid (hydrocephalus). Hind leg: two foci of myodegeneration and necrosis (85280, 85281). Some regenerating myofibres identified. Spinal cord: few foci of myodegeneration (85284). Head: few foci of cytoplasmic hyaline droplet accumulation in the nasal epithelium with scattered mucosal neutrophils (85313). Head: inflammation of a single hair follicle in the upper lip (85314)

#### #5 (control)

Macro Observations

Tail suspension test for neurological defects - negative.
Dentition, tongue and oral cavity was unremarkable.
BCS: 3
Spleen: 18×5×2mm
Kidneys: 12×8×6mm, symmetrical
Thymus: 6×6×2mm
Lungs inflated.
Heart: 12×6×5mm
Brain: 16×11×5mm, symmetrical
Pituitary gland identified, macroscopically normal
Tail length: 85mm (straight)
Head harvested for evaluation of auditory and vestibular structures.
Bone marrow smear taken from left hind leg.
No macroscopic lesions identified.

Micro Observations

Animal #5 was used as a female histological control for this cohort.
Marrow smear: Examination of the smear showed representative cells from the myeloid and erythroid series. Occasional cells from the lymphoid series. Occasional and unremarkable megakaryoblasts. (85084)
Peripheral blood smear: Examination of the smear showed red blood cells (majority of cells shown), occasional white blood cells including lymphocytes, segmented neutrophils, monocytes and platelets (clumps). No discernible morphological changes or detectable parasites. (85085)
Organs examined: Mammary glands (85092, 85096), Ovaries/oviducts (85086, 85087), Uterus/cervix/vagina (85086, 85087), Urinary Bladder (85086, 85087), Liver/Gall Bladder (85088), Stomach (85089), Duodenum/jejunum/ileum/GALT (85089, 85090, 85091), Cecum/Colon/GALT (85091), Mesenteric Lymph Node (85092), Spleen (85086, 85087), Pancreas (85086, 85087), Kidneys/Adrenal Glands (85093), Salivary Glands/regional lymph nodes (85092), Thyroids (85094, 85095), Trachea/Lungs (85094, 85095), Thymus (85094, 85095), Heart (85094, 85095), Skin (85096), Tail (85247, 85248, 85249), Eyes/Harderian Glands (85097), Brain (85098), Spinal cord (85245, 85246), Hind leg (85247, 85248, 85249), Head (85292, 85293), Sternum (85244).
Micromorphological changes- Ovaries: occasional pigment laden interstitial cells (likely ceroid-lipofucsin) (85086, 85087). Oviducts: epithelial cytoplasmic vacuolation, bilateral (85086, 85087). Uterus: cystic endometrial hyperplasia (85086, 85087). Liver: mild multifocal perivascular mononuclear cell infiltrates (85088). Liver: small foci of hepatocyte cell loss/necrosis and associated leucocyte infiltrates (mixed) (85088). Gall bladder: scattered neutrophils in the lamina propria (85088). Large intestine: of the cecum, abundant luminal protozoa, and increased mononuclear cell infiltrates in the lamina propria (85091). Pancreas: minimal multifocal perivascular mononuclear cell infiltrates (85086, 85087). Pancreas: focal loss/necrosis of the exocrine pancreas (acini) with mainly mononuclear cell infiltrates (85089). Kidneys: several protein casts within the cortex and medulla regions. Few in the medulla are mildly dilated (85093). Kidney: minimal multifocal perivascular and parenchymal mononuclear cell infiltrates (85093). Salivary glands: mild multifocal perivascular mononuclear cell infiltrates within the submandibular gland (85092). Thymus: single small sized cyst (85094). Lungs: few small foci of intra-alveolar haemorrhage, judged to be artefactual (85094, 85095). Heart: few foci of mononuclear cell infiltrates and cardiomyocyte degeneration (85094). Heart: myxomatous valvular changes (thickened leaflets) (85094). Brain: some flattening/stretching of the ependymal cells in the lateral ventricles (85098), indicative of increased cerebrospinal fluid (hydrocephalus). Uterus: cystic endometrial hyperplasia (85086, 85087). Liver: mild multifocal perivascular mononuclear cell infiltrates (85088). Liver: small foci of hepatocyte cell loss/necrosis and associated leucocyte infiltrates (mixed) (85088). Gall bladder: scattered neutrophils in the lamina propria (85088). Large intestine: of the cecum, abundant luminal protozoa, and increased mononuclear cell infiltrates in the lamina propria (85091). Pancreas: minimal multifocal perivascular mononuclear cell infiltrates (85086, 85087). Pancreas: focal loss/necrosis of the exocrine pancreas (acini) with mainly mononuclear cell infiltrates (85089). Kidneys: several protein casts within the cortex and medulla regions. Few in the medulla are mildly dilated (85093). Kidney: minimal multifocal perivascular and parenchymal mononuclear cell infiltrates (85093). Salivary glands: mild multifocal perivascular mononuclear cell infiltrates within the submandibular gland (85092). Thymus: single small sized cyst (85094). Lungs: few small foci of intra-alveolar haemorrhage, judged to be artefactual (85094, 85095). Heart: few foci of mononuclear cell infiltrates and cardiomyocyte degeneration (85094). Heart: myxomatous valvular changes (thickened leaflets) (85094). Brain: some flattening/stretching of the ependymal cells in the lateral ventricles (85098), indicative of increased cerebrospinal fluid (hydrocephalus). Spinal: minimal focal eosinophilic infiltrate in the surrounding connective tissue (85245). Head: minimal multifocal peri-glandular mononuclear cell infiltrates within the cervical mammary tissue (85292, 85293). Head: few foci of mixed leucocyte infiltrates within the neck muscle (85292). Multiple small to parge sized perivascular mononuclear cell aggregates within the surrounding peri-adipose tissues of the lungs and reproductive organs.

#### #6 (control)

Macro Observations

Tail suspension test for neurological defects - negative.
Dentition, tongue and oral cavity was unremarkable.
BCS: 3
Spleen: 13×4×2mm
Kidneys: 11×6×4mm, symmetrical
Thymus: 7×7×2mm
Lungs inflated.
Heart: 9×8×5mm
Brain: 15×10×5mm, symmetrical
Pituitary gland identified, macroscopically normal
Tail length: 84mm (straight)
Head harvested for evaluation of auditory and vestibular structures.
Bone marrow smear taken from left hind leg.
No macroscopic lesions identified
Notable head shape-arch of snout (tip of nose) to crown (top of head between the ears) was prominently rounded/convex.

Micro Observations

Animal #6 was used as a female histological control for this cohort.
Marrow smear: Examination of the smear showed representative cells from the myeloid and erythroid series. Occasional cells from the lymphoid series. Occasional and unremarkable megakaryoblasts. (85099)
Peripheral blood smear: Examination of the smear showed red blood cells (majority of cells shown), occasional white blood cells including lymphocytes, segmented neutrophils, monocytes and platelets (clumps). No discernible morphological changes or detectable parasites. (85100)
Organs examined: Mammary glands (85107, 85111), Ovaries/oviducts (85101, 85102), Uterus/cervix/vagina (85101, 85102), Urinary Bladder (85101, 85102), Liver/Gall Bladder (85103), Stomach (85104), Duodenum/jejunum/ ileum/GALT (85104, 85105, 85106), Cecum/Colon/GALT (85106), Mesenteric Lymph Node (85107), Spleen (85101, 85102), Pancreas (85101, 85102), Kidneys/Adrenal Gland (85108), Salivary Glands/regional lymph nodes (85107), Thyroids (85109), Trachea/Lungs (85109, 85110), Thymus (85109, 85110), Heart (85109, 85110), Skin (85111), Tail (85234, 85235, 85236), Eyes/Harderian Glands (85112), Brain (85113), Spinal cord (85231, 85232), Hind leg (85234, 85235, 85236), Head (85294, 85295, 85296), Sternum (85233).
Micromorphological changes- Ovaries: occasional pigment laden interstitial cells (likely ceroid-lipofucsin) (85101, 85102). Oviducts: focal epithelial cytoplasmic vacuolation, bilateral (85101, 85102). Uterus: cystic endometrial hyperplasia (85101, 85102). Liver: multiple small parenchymal leukocyte aggregates with some hepatocyte cell loss/necrosis (85103). Liver: minimal multifocal perivascular mononuclear cell infiltrates (85103). Gall bladder: focal epithelial hyperplasia, and underlying neutrophils in the lamina propria (85103). Small intestine: of the distal half, increased number of eosinophils in the lamina propria (85105, 85106). Large intestine: in mainly the cecum, abundant luminal protozoa (85106). Pancreas: minimal focal perivascular mononuclear cell infiltrate (85101). Salivary glands: of the parotid gland, small cluster of enlarged basophilic acini cells nearby a small focus of glandular atrophy (ducts and acini) (85107). Salivary glands: another larger focus of glandular atrophy (ducts and acini), likely of the parotid gland (85107). Kidney: multiple mildly dilated protein casts within the medulla (85108). Thymus: few small sized cysts (85109, 85110). Heart: myxomatous valvular changes (thickened leaflets) (85109). Heart: focal myocardiocyte degeneration of the ventricular wall (85109). Lungs: single small sized parenchymal mononuclear cell aggregate (85110). Lungs: few small mononuclear cell aggregates within the surrounding peri-adipose tissue (85109, 85110). Brain: some flattening/stretching of the ependymal cells in the lateral ventricles (85113), indicative of increased cerebrospinal fluid (hydrocephalus). Hind leg: multifocal mixed leucocyte inflammation within the digits of the foot (85235, 85236). Hing leg: oedamatous changes within the subcutis above the foot (85234, 85235, 85236). Head: some hyaline droplet accumulation within the nasal epithelium (85294, 85295, 85296). Head: multiple small inflammatory foci in the skin of the outer ear (85295, 85296), query focal inflammation/necrosis of the outer surface of the tympanic membrane (85295).

#### #11

Macro Observations

Tail suspension test for neurological defects - negative.
Dentition, tongue and oral cavity was unremarkable.
BCS: 3
Spleen: 13×4×3mm
Kidneys: 11×6×5mm, symmetrical
Thymus: 8×7×2mm
Lungs inflated.
Heart: 8×8×6mm
Brain: 14×10×5mm, symmetrical
Pituitary gland identified, macroscopically normal
Tail length: 81mm (straight)
Head harvested for evaluation of auditory and vestibular structures.
Bone marrow smear taken from left hind leg.
No macroscopic lesions identified.

Micro Observations

Marrow smear: Examination of the smear showed representative cells from the myeloid and erythroid series. Occasional cells from the lymphoid series. Occasional and unremarkable megakaryoblasts. (85148)
Peripheral blood smear: Examination of the smear showed red blood cells (majority of cells shown), occasional white blood cells including lymphocytes, segmented neutrophils, monocytes and platelets (clumps). No discernible morphological changes or detectable parasites. (85149)

Mammary glands

Typical mammary fat pad with developing lactiferous ducts, blood vessels, and nerve bundles.
(85156) No lesions of significance

Ovaries/Oviducts

Unremarkable ovaries containing follicles at various stages of development (primary through to antral) and several corpora lutea.
Clusters of pigment laden interstitial cells (likely ceroid-lipofucsin), an age-related change - feature seen in the controls.
Mostly unremarkable oviduct micromorphology with typical columnar epithelium and mucosal folds.
Focal epithelial cytoplasmic vacuolation, an age-related change - feature seen in the controls. (85150, 85151)
No lesions of significance

Uterus/Cervix/Vagina/Clitoral gland

Cystic endometrial hyperplasia, an age-related change - feature seen in the controls.
Unremarkable architecture myometrium and adventitia.
The micromorphology of the uterus and vagina places the animal at proestrus.
(85150, 85151)
No lesions of significance

Urinary Bladder

Unremarkable bladder with typical urothelium and detrusor muscle.
(85150, 85151)
No lesions of significance

Liver/Gall bladder

Liver parenchyma including hepatocytes, Kupffer cells, portal triads and central veins.
Areas of hydropic degeneration of hepatocytes, an age-related finding in mice.
Multiple small foci of parenchymal leukocyte aggregates and some hepatocyte cell loss/necrosis - common incidental/background finding - feature seen in the controls.
Gall bladder with epithelial hyperplasia, cytoplasmic hyaline droplet accumulation, and underlying neutrophils in the lamina propria, common incidental/background finding - feature seen in the controls.
(85152)
*Comments:*
*Pathology to comment*

Stomach

Unremarkable fore and glandular portions of the stomach with limiting ridge.
Includes pyloric sphincter and duodenal bulb.
(85153)
No lesions of significance

Small Intestine (Duodenum, Jejunum & Ileum)/GALT

Typical mucosal villi and submucosal layers.
Peyer’s patches display typical reactive nodal histology. Occasional, typical lymphoid cluster (cryptopatches).
(85153, 85154, 85155)
No lesions of significance

Cecum/Colon/GALT

Typical mucosal folds and submucosal layers. Unremarkable muscularis and discernible ganglion cells of the plexuses.
Occasional, typical lymphoid cluster (Peyer’s patch).
(85155)
No lesions of significance

Mesenteric lymph node

Typical reactive nodal histology including enlarged size, mild follicular hyperplasia, an expansive paracortical area with some lymphocyte apoptosis, and sinus histiocytosis.
(85156)
No lesions of significance

Spleen

Mild follicular hyperplasia with germinal centre formation and some lymphocyte apoptosis. Mild coalescence of the white pulp.
Expansive red pulp with marked extramedullary haematopoiesis and conspicuous haemosiderin laden macrophages.
(85151)
No lesions of significance

Pancreas

Representative exocrine tissue (serous acini) and endocrine tissue (islets of Langerhans).
(85150, 85151)
No lesions of significance

Kidney

Sections show a cortex, medulla, and papilla. There is a uniform distribution of glomeruli and accompanying nephron components and the micromorphology of the tubules is unremarkable. Minimal perivascular/peripelvic mononuclear cell infiltrates, incidental/age-related change - feature seen in the controls.
Several protein casts within the medulla and papilla, moderately dilated within the medulla, incidental/age-related change - feature seen in the controls.
Renal lymph nodes with typical reactive nodal histology and mild sinus histiocytosis.
(85157)
No lesions of significance

Adrenal glands

Adrenal glands with typical cortex/medulla micromorphology.
(85157)
No lesions of significance

Salivary glands and Regional lymph nodes

Unremarkable submandibular, sublingual and parotid glands.
Regional lymph nodes display typical reactive nodal histology.
(85156)
No lesions of significance

Thyroids

Normal lateral lobes of the thyroid gland with typical colloid secreting follicles lined by cuboidal epithelium.
Small sheet-like mass of polygonal cells, characteristic of the parathyroid gland.
(85158, 85159)
No lesions of significance

Trachea/Lungs

Typical lung parenchyma/alveoli, bronchioles, blood vessels and parabronchial lymph node. Minimal multifocal perivascular mononuclear cell infiltrates, an incidental/age-related change - feature seen in the controls.
Trachea with unremarkable mucosal epithelial lining and hyaline cartilage.
Oesophagus with typical features including stratified squamous epithelium.
(85158, 85159)
No lesions of significance

Thymus

Typical medulla/cortex distribution and micromorphology.
Single small sized cyst (85158), common incidental/background finding - feature seen in the controls.
(85158, 85159)
No lesions of significance

Heart/chambers/vessels/valves

Mostly typical micromorphology observed in cardiac muscle, chambers, valves and great vessels of the heart.
Myxomatous valvular changes (thickened leaflets), an age-related change - feature seen in the controls.
The cardiac muscle fibres demonstrated typical features including central nuclei, branching fibres and striations.
(85158, 85159)
No lesions of significance

Skin

Typical dermal appendages and distribution. Unremarkable thin layer of striated muscle (panniculus carnosus).
Minimal focal mononuclear cell infiltrate surrounding one hair follicle, common incidental finding in mice.
(85160)
No lesions of significance

Tail

Typical tail components including keratinized squamous epithelium, dense regular connective tissue, tendons, caudal vertebra, bone marrow, intervertebral disc, skeletal muscle, nerves and blood vessels.
(85267, 85268, 85269)
No lesions of significance

Eyes/Harderian glands

Unremarkable retina, cornea, iris, ciliary body, lens, sclera and choroid.
Typical branched tubuloalveolar formation of the Harderian gland.
Includes portion of unremarkable optic nerve and extraocular muscles.
(85161)
No lesions of significance

Brain

Sections were prepared from the standard levels of the brain:
Level I forebrain: including cortex, corpus callosum, caudate putamen and lateral ventricles. Level II midbrain: including the hippocampus, thalamus, hypothalamus and lateral and third ventricles. Level III hindbrain: includes the cerebellum, pons and fourth ventricle.
Sections of brain appear symmetrical with unremarkable meninges and typical lamination. The cerebellum appears symmetrical with typical architecture and Purkinje cells. There was no obvious neuronal loss and the myelination appears normal.
Some flattening/stretching of the ependymal cells and increased size of the lateral ventricular space, features indicative of increased cerebrospinal fluid (hydrocephalus), a known background lesion in mice - similar feature seen in the controls.
(85162)
*Comments:*
*Neuropathology to comment*

Spinal cord

Representative thoracic and lumbar region of spinal cord, vertebral bone, intervertebral disc, striated muscle, peripheral nerves, brown adipose tissue, and bone marrow. (85271, 85272) No lesions of significance
*Comments:*
*Neuropathology to comment*

(Hind leg) Long bone/Bone marrow/Synovial joint/Skeletal muscle

Mostly unremarkable long bone, striated muscle, examples of nerve fascicles, fibrocartilage of the meniscus, synovial joint and bone marrow. The skeletal muscle shows consistent fibre size with peripheral nuclei.
Focal myodegeneration adjacent the hip joint (85269) - feature seen in the controls. (85267, 85268, 85269)
*Comments:*
*Pathology to comment*

Head

Multiple levels through the head demonstrate dermal appendages, nasal cavity, oral cavity, teeth and tongue including muscle bundles. Sections also show unremarkable pituitary gland including pars intermedia, pars distalis and pars nervosa as well and the trigeminal nerve/ganglia (85303). The outer and middle regions of the ear are discernible. The tympanic membrane is intact and the ossicles are unremarkable and include the stapedial annular ligaments (85304-85306). Typical components of the inner ear including bony labyrinth, organ of corti, stria vascularis and scala cavities are discernible. Based on multiple levels, the organ of corti is unremarkable with no discernible loss of inner/outer hair cells and typical tectorial membrane (85304-85306). The cochlear nerve and spiral ganglion is also demonstrated and based on several levels, there is no reduction in the density of the spiral ganglion cells. Examples of otolith organs can be seen with typical features such as the hair cells and mineral otoliths. The ampulla including the crista ridge with hair cells is discernible (85306).
Focal and bilateral inflammation of the nasal epithelium, seemingly due to inhaled foreign material.
(85303-85306) No lesions of significance

Sternum

Representative sternebrae, costal cartilage, intersternebral joint, intercostal skeletal muscle and brown fat. Haematopoietic tissue islands surrounded by vascular sinuses interspersed within a meshwork of trabecular bone. The bone marrow morphology demonstrated typical myeloid features including conspicuous megakaryoblasts and lymphoid features. (85270) No lesions of significance

#### #12

Macro Observations

Tail suspension test for neurological defects - negative. Dentition, tongue and oral cavity was unremarkable. BCS: 3 Testes: 8×5×4mm, symmetrical Spleen: 15×5×2mm Kidneys: 14×8×5mm, symmetrical Thymus: 6×6×2mm Lungs inflated. Heart: 10×8×6mm Brain: 16×10×5mm, symmetrical Pituitary gland identified, macroscopically normal Tail length: 85mm (straight) Head harvested for evaluation of auditory and vestibular structures. Bone marrow smear taken from left hind leg.
No macroscopic lesions identified.

Micro Observations

Marrow smear: Examination of the smear showed representative cells from the myeloid and erythroid series. Occasional cells from the lymphoid series. Occasional and unremarkable megakaryoblasts. (85163)
Peripheral blood smear: Examination of the smear showed red blood cells (majority of cells shown), occasional white blood cells including lymphocytes, segmented neutrophils, monocytes and platelets (clumps). No discernible morphological changes or detectable parasites. (85164)

Mammary glands

Typical mammary fat pad with developing lactiferous ducts, blood vessels, and nerve bundles. (85175) No lesions of significance

Ovaries/Oviducts

Unremarkable ovaries containing follicles at various stages of development (primary through to antral) and several corpora lutea. Scattered pigment laden interstitial cells (likely ceroid-lipofucsin), an age-related change - feature seen in the controls. Single ovarian cyst, an incidental/background finding in mice. Mononuclear cell infiltration of the bursa membrane, unilateral, an age-related finding in mice. Mostly unremarkable oviduct micromorphology with typical columnar epithelium and mucosal folds. Focal epithelial cytoplasmic vacuolation, an age-related change - feature seen in the controls. (85165, 85166) No lesions of significance

Uterus/Cervix/Vagina/Clitoral gland

Endometrial hyperplasia, an age-related change - feature seen in the controls. Unremarkable architecture myometrium and adventitia. (85165, 85166) No lesions of significance

Urinary Bladder

Unremarkable bladder with typical urothelium and detrusor muscle. (85165, 85166) No lesions of significance

Liver/Gall bladder

Liver parenchyma including hepatocytes, Kupffer cells, portal triads and central veins.
Areas of hydropic degeneration of hepatocytes, an age-related finding in mice.
Several small parenchymal leukocyte aggregates - common incidental/background finding - feature seen in the controls.
Section does not include gall bladder.
(85167)
*Comments:*
*Pathology to comment*

Stomach

Unremarkable fore and glandular portions of the stomach with limiting ridge.
Small focal basophilic deposit within the gland lumen (?mineralisation) - likely an incidental finding.
Includes pyloric sphincter and duodenal bulb.
(85168)
No lesions of significance

Small Intestine (Duodenum, Jejunum & Ileum)/GALT

Typical mucosal villi and submucosal layers.
Peyer’s patches display typical reactive nodal histology. Occasional, typical lymphoid cluster (cryptopatches).
(85168, 85169, 85170)
No lesions of significance

Cecum/Colon/GALT

Typical mucosal folds and submucosal layers. Unremarkable muscularis and discernible ganglion cells of the plexuses.
Occasional, typical lymphoid cluster (Peyer’s patch).
In mainly the cecum, abundant luminal protozoa - feature seen in the controls.
(85170)
No lesions of significance

Mesenteric lymph node

Typical reactive nodal histology with enlarged size, representative cortex including the occasional follicle, an expansive paracortical area with lymphocyte apoptosis, and sinus histiocytosis.
(85171)
No lesions of significance

Spleen

Mild follicular hyperplasia with germinal centre formation and some lymphocyte apoptosis. Mild coalescence of the white pulp.
Expansive red pulp with marked extramedullary haematopoiesis.
(85166)
No lesions of significance

Pancreas

Representative exocrine tissue (serous acini) and endocrine tissue (islets of Langerhans).
(85166)
No lesions of significance

Kidney

Sections show a cortex, medulla, and papilla. There is a uniform distribution of glomeruli and accompanying nephron components and the micromorphology of the tubules is unremarkable. Minimal perivascular/peripelvic mononuclear cell infiltrates, incidental/age-related change - feature seen in the controls.
Few protein casts within the medulla, incidental/age-related change - feature seen in the controls.
Renal lymph nodes with typical reactive nodal histology.
(85172)
No lesions of significance

Adrenal glands

Adrenal glands with typical cortex/medulla micromorphology.
(85172)
No lesions of significance

Salivary glands and Regional lymph nodes

Unremarkable submandibular, sublingual and parotid glands.
Mild multifocal perivascular mononuclear cell infiltrates within the submandibular gland, an age-related change - feature seen in the controls.
Regional lymph nodes display typical reactive nodal histology. (85171)
No lesions of significance

Thyroids

Normal lateral lobes of the thyroid gland with typical colloid secreting follicles lined by cuboidal epithelium.
Small sheet-like mass of polygonal cells, characteristic of the parathyroid gland. (85173, 85174)
No lesions of significance

Trachea/Lungs

Typical lung parenchyma/alveoli, bronchioles, blood vessels and parabronchial lymph node. Mild multifocal perivascular/peribronchial mononuclear cell infiltrates, an incidental/age-related change - feature seen in the controls. Trachea with unremarkable mucosal epithelial lining and hyaline cartilage.
Oesophagus with typical features including stratified squamous epithelium.
Several small to medium sized mononuclear cell aggegrates in the surrounding peri-adipose tissue, common finding - feature seen in the controls.
(85173, 85174)
No lesions of significance

Thymus

Typical medulla/cortex distribution and micromorphology.
(85173, 85174)
No lesions of significance

Heart/chambers/vessels/valves

Mostly typical micromorphology observed in cardiac muscle, chambers, valves and great vessels of the heart.
Myxomatous valvular changes (thickened leaflets), an age-related change - feature seen in the controls.
The cardiac muscle fibres demonstrated typical features including central nuclei, branching fibres and striations.
(85173, 85174)
No lesions of significance

Skin

Typical dermal appendages and distribution. Unremarkable thin layer of striated muscle (panniculus carnosus).
(85175)
No lesions of significance

Tail

Typical tail components including keratinized squamous epithelium, dense regular connective tissue, tendons, caudal vertebra, bone marrow, intervertebral disc, skeletal muscle, nerves and blood vessels.
(85261, 85262, 85263)
No lesions of significance

Eyes/Harderian glands

Unremarkable retina, cornea, iris, ciliary body, lens, sclera and choroid.
Typical branched tubuloalveolar formation of the Harderian gland.
Includes portion of unremarkable optic nerve and extraocular muscles.
(85176)
No lesions of significance

Brain

Sections were prepared from the standard levels of the brain:
Level I forebrain: including cortex, corpus callosum, caudate putamen and lateral ventricles. Level II midbrain: including the hippocampus, thalamus, hypothalamus and lateral and third ventricles. Level III hindbrain: includes the cerebellum, pons and fourth ventricle.
Sections of brain appear symmetrical with unremarkable meninges and typical lamination. The cerebellum appears symmetrical with typical architecture and Purkinje cells. There was no obvious neuronal loss and the myelination appears normal.
Prominent flattening/stretching of the ependymal cells and increased size of the lateral ventricular space, features indicative of increased cerebrospinal fluid (hydrocephalus), a known background lesion in mice - similar feature seen in the controls.
Query vacuolation of the adjacent white matter, possible artefact.
(85177)
*Comments:*
Neuropathology *to comment*

Spinal cord

Representative thoracic and lumbar region of spinal cord, vertebral bone, intervertebral disc, striated muscle, peripheral nerves, brown adipose tissue, and bone marrow.
(85265, 85266)
No lesions of significance
*Comments:*
*Neuropathology to comment*

(Hind leg) Long bone/Bone marrow/Synovial joint/Skeletal muscle

Unremarkable long bone, striated muscle, examples of nerve fascicles, fibrocartilage of the meniscus, synovial joint and bone marrow. The skeletal muscle shows consistent fibre size with peripheral nuclei.
Small focus of accumulated mononuclear cell infiltrates in the dermis (85263), common incidental finding.
(85261, 85262, 85263)
No lesions of significance

Head

Multiple levels through the head demonstrate dermal appendages, nasal cavity, oral cavity, teeth and tongue including muscle bundles. Sections also show unremarkable pituitary gland including pars intermedia, pars distalis and pars nervosa as well and the trigeminal nerve/ganglia (85307). The outer and middle regions of the ear are discernible. The tympanic membrane is intact and the ossicles are unremarkable and include the stapedial annular ligaments (85308, 85309). Typical components of the inner ear including bony labyrinth, organ of corti, stria vascularis and scala cavities are discernible. Based on multiple levels, the organ of corti is unremarkable with no discernible loss of inner/outer hair cells and typical tectorial membrane (85308, 85309). The cochlear nerve and spiral ganglion is also demonstrated and based on several levels, there is no reduction in the density of the spiral ganglion cells. Examples of otolith organs can be seen with typical features such as the hair cells and mineral otoliths. The ampulla including the crista ridge with hair cells is discernible (85308, 85309).
(85307-85309)
No lesions of significance

Sternum

Representative sternebrae, costal cartilage, intersternebral joint, intercostal skeletal muscle and brown fat.
Haematopoietic tissue islands surrounded by vascular sinuses interspersed within a meshwork of trabecular bone. The bone marrow morphology demonstrated typical myeloid features including conspicuous megakaryoblasts and lymphoid features.
(85264)
No lesions of significance

#### #18

Macro Observations

Tail suspension test for neurological defects - negative.
Dentition, tongue and oral cavity was unremarkable.
BCS: 3
Testes: 8×6×4mm, symmetrical
Spleen: 18×6×2mm
Kidneys: 15×8×6mm, symmetrical
Thymus: 7×7×2mm
Lungs inflated.
Heart: 12×8×6mm
Brain: 16×11×6mm, symmetrical
Pituitary gland identified, macroscopically normal
Tail length: 80mm (straight)
Head harvested for evaluation of auditory and vestibular structures.
Bone marrow smear taken from left hind leg.
No macroscopic lesions identified.

Micro Observations

Marrow smear: Examination of the smear showed representative cells from the myeloid and erythroid series. Occasional cells from the lymphoid series. Occasional and unremarkable megakaryoblasts.
(85178)
Peripheral blood smear: Examination of the smear showed red blood cells (majority of cells shown), occasional white blood cells including lymphocytes, segmented neutrophils, monocytes and platelets (clumps). No discernible morphological changes or detectable parasites.
(85179)

Testes/Epididymes

Mostly typical convoluted seminiferous tubules at various stages of cycle surrounded by the tunica albuginea. Within the tubules, unremarkable spermatogenic cells including, Sertoli cells, spermatogonia, developing spermatocytes and spermatids. Typical interstitial Leydig cells. Multifocal tubular degeneration/atrophy, an age-related change - feature seen in the control. Unremarkable vas deferens with typical intraluminal sperm. Architecture of the epididymis is mostly typical, with numerous intraluminal elongated spermatozoa. Focal interstitial oedema-like changes in an epididymis, possible plane of section artefact. (85180)

Seminal vesicles

Unremarkable tall columnar epithelium and folded mucosa.
Presence of typical intraluminal eosinophilic secretions.
(85181, 85182)
No lesions of significance

Prostate glands

Unremarkable dorsal lateral/ventral/coagulating glands with typical intraluminal secretions.
Portion of unremarkable vas deferens.
(85181, 85182)
No lesions of significance

Penis/Preputial gland

Typical penile structures including prepuce, glans, corpus cavernosum and urethra.
Typical preputial glands including basal and secretory cells.
Skin shows focal accummulation of dermal melanocytes, a common incidental/background finding in mice.
(85183)
No lesions of significance

Urinary Bladder

Unremarkable bladder with typical urothelium and detrusor muscle.\
(85181, 85182)
No lesions of significance

Liver/Gall bladder

Liver parenchyma including hepatocytes, Kupffer cells, portal triads and central veins.
Several small parenchymal leucocyte aggregates, incidental/background finding - feature seen in the controls.
Unremarkable gall bladder/section does not show gall bladder.
(85184)
No lesions of significance

Stomach

Mostly unremarkable fore and glandular portions of the stomach with limiting ridge.
Segmental eosinophil infiltrates in the submucsa and lamina propria, common incidental finding in mice.
Includes pyloric sphincter and duodenal bulb.
(85185)
No lesions of significance

Small Intestine (Duodenum, Jejunum & Ileum)/GALT

Typical mucosal villi and submucosal layers.
Peyer’s patches display typical nodal histology. Occasional, typical lymphoid cluster (cryptopatches).
(85185, 85186, 85187)
No lesions of significance

Cecum/Colon/GALT

Typical mucosal folds and submucosal layers. Unremarkable muscularis and discernible ganglion cells of the plexuses.
Occasional, typical lymphoid cluster (Peyer’s patch). (85187)
No lesions of significance

Mesenteric lymph node

Typical reactive nodal histology with enlarged size, representative cortex including the occasional follicle, an expansive paracortical area with some lymphocyte apoptosis, and sinus histiocytosis.
(85188)
No lesions of significance

Spleen

Mild follicular hyperplasia with germinal centre formation and some lymphocyte apoptosis. Mild coalescence of the white pulp.
Extramedullary haematopoiesis in the red pulp.
(85181, 85182)
No lesions of significance

Pancreas

Representative exocrine tissue (serous acini) and endocrine tissue (islets of Langerhans).
(85181, 85182)
No lesions of significance

Kidney

Sections show a cortex, medulla, and papilla. There is a uniform distribution of glomeruli and accompanying nephron components and the micromorphology of the tubules is unremarkable. Minimal perivascular/peripelvic mononuclear cell infiltrates, incidental/age-related change - feature seen in the controls.
Few protein casts within the medulla, incidental/age-related change - feature seen in the controls.
Focal cluster of irregular tubules showing tubule basophilia and shrunken cytoplasm/nuclear crowding, an incidental/background finding in mice.
Renal lymph nodes with typical reactive nodal histology and mild sinus histiocytosis. (85189)
No lesions of significance

Adrenal glands

Adrenal glands with typical cortex/medulla micromorphology.
(85189)
No lesions of significance

Salivary glands and Regional lymph nodes

Unremarkable submandibular, sublingual and parotid glands.
Minimal multifocal perivascular mononuclear cell infiltrates within the submandibular gland, an age-related change - feature seen in the controls.
Regional lymph nodes display typical reactive nodal histology. (85188)
No lesions of significance

Thyroids

Typical colloid secreting follicles lined by cuboidal epithelium.
(85190, 85191)
No lesions of significance

Trachea/Lungs

Typical lung parenchyma/alveoli, bronchioles, and blood vessels.
Minimal perivascular mononuclear cell infiltrates, an incidental/age-related change - feature seen in the controls.
Trachea with unremarkable mucosal epithelial lining and hyaline cartilage. Oesophagus with typical features including stratified squamous epithelium.
(85190, 85191)
No lesions of significance

Thymus

Typical medulla/cortex distribution and micromorphology.
(85190, 85191)
No lesions of significance

Heart/chambers/vessels/valves

Mostly typical micromorphology observed in cardiac muscle, chambers, valves and great vessels of the heart.
Myxomatous valvular changes (thickened leaflets) (85191), incidental/age-related finding - feature seen in the controls.
Several foci of cardiomycoyte vacuolation and degeneration, likely incidental/age-related - feature seen in the controls.
Remaining cardiac muscle fibres demonstrated typical features including central nuclei, branching fibres and striations.
(85190, 85191)

Skin

Typical dermal appendages and distribution. Unremarkable thin layer of striated muscle (panniculus carnosus).
(85192)
No lesions of significance

Tail

Typical tail components including keratinized squamous epithelium, dense regular connective tissue, tendons, caudal vertebra, bone marrow, intervertebral disc, skeletal muscle, nerves and blood vessels.
(85273, 85274, 85275)
No lesions of significance

Eyes/Harderian glands

Unremarkable retina, cornea, iris, ciliary body, lens, sclera and choroid.
Typical branched tubuloalveolar formation of the Harderian gland.
Includes portion of unremarkable optic nerve and extraocular muscles. (85193)
No lesions of significance

Brain

Sections were prepared from the standard levels of the brain:
Level I forebrain: including cortex, corpus callosum, caudate putamen and lateral ventricles. Level II midbrain: including the hippocampus, thalamus, hypothalamus and lateral and third ventricles. Level III hindbrain: includes the cerebellum, pons and fourth ventricle.
Sections of brain appear symmetrical with unremarkable meninges and typical lamination.
The cerebellum appears symmetrical with typical architecture and Purkinje cells.
There was no obvious neuronal loss and the myelination appears normal.
Some flattening/stretching of the ependymal cells, indicative of increased cerebrospinal fluid (hydrocephalus), a known background lesion in mice - similar feature seen in the controls.
(85194)
*Comments:*
*Neuropathology to comment*

Spinal cord

Representative thoracic and lumbar region of spinal cord, vertebral bone, intervertebral disc, striated muscle, peripheral nerves, brown adipose tissue, and bone marrow.
(85277, 85278)
No lesions of significance
*Comments:*
*Neuropathology to comment*

(Hind leg) Long bone/Bone marrow/Synovial joint/Skeletal muscle

Representative long bone, striated muscle, examples of nerve fascicles, fibrocartilage of the meniscus, synovial joint and bone marrow. The skeletal muscle shows consistent fibre size with peripheral nuclei.
Query focal necrosis in the foot (?bone) (85273).
(85273, 85274, 85275)
*Comments:*
*Pathology to comment*

Head

Multiple levels through the head demonstrate dermal appendages, nasal cavity, oral cavity, teeth and tongue including muscle bundles. Sections also show unremarkable pituitary gland including pars intermedia, pars distalis and pars nervosa as well and the trigeminal nerve/ganglia (85310). The outer and middle regions of the ear are discernible. The tympanic membrane is intact and the ossicles are unremarkable and include the stapedial annular ligaments.
Typical components of the inner ear including bony labyrinth, organ of corti, stria vascularis and scala cavities are discernible. Based on multiple levels, the organ of corti is unremarkable with no discernible loss of inner/outer hair cells and typical tectorial membrane.
The cochlear nerve and spiral ganglion is also demonstrated and based on several levels, there is no reduction in the density of the spiral ganglion cells. Examples of otolith organs can be seen with typical features such as the hair cells and mineral otoliths. The ampulla including the crista ridge with hair cells is discernible (85311).
Inflammation of the middle ear characterised by oedema, luminal cellular debris, multinucleated giant cells, and basophilic deposit/mineralisation, unilateral - an incidental/background finding in mice.
(85310, 85311)

Sternum

Representative sternebrae, costal cartilage, intersternebral joint, intercostal skeletal muscle and brown fat.
Haematopoietic tissue islands surrounded by vascular sinuses interspersed within a meshwork of trabecular bone. The bone marrow morphology demonstrated typical myeloid features including conspicuous megakaryoblasts and lymphoid features.
(85276)
No lesions of significance

#### #3

Macro Observations

Tail suspension test for neurological defects - negative.
Dentition, tongue and oral cavity was unremarkable.
BCS: 3
Testes: 8×5×4mm, symmetrical
Spleen: 16×5×3mm
Kidneys: 13×7×4mm, symmetrical
Thymus: 7×6×2mm
Lungs inflated.
Heart: 11×8×7mm
Brain: 15×10×5mm, symmetrical
Pituitary gland identified, macroscopically normal
Tail length: 84mm (straight)
Head harvested for evaluation of auditory and vestibular structures.
Bone marrow smear taken from left hind leg.
Fur was slightly ruffled.
No macroscopic lesions identified.

Micro Observations

Marrow smear: Examination of the smear showed representative cells from the myeloid and erythroid series. Occasional cells from the lymphoid series. Occasional and unremarkable megakaryoblasts.
(85050)
Peripheral blood smear: Examination of the smear showed red blood cells (majority of cells shown), occasional white blood cells including lymphocytes, segmented neutrophils, monocytes and platelets (clumps). No discernible morphological changes or detectable parasites.
(85051)

Testes/Epididymes

Mostly typical convoluted seminiferous tubules at various stages of cycle surrounded by the tunica albuginea. Within the tubules, unremarkable spermatogenic cells including, Sertoli cells, spermatogonia, developing spermatocytes and spermatids. Typical interstitial Leydig cells.
Focal cluster of tubules showing degeneration/atrophy, an age-related change - feature seen in the control.
Unremarkable vas deferens with typical intraluminal sperm.
Architecture of the epididymis is typical, with numerous intraluminal elongated spermatozoa. (85052)
No lesions of significance

Seminal vesicles

Unremarkable tall columnar epithelium and folded mucosa. Presence of typical intraluminal eosinophilic secretions.
(85053, 85054)
No lesions of significance

Prostate glands

Unremarkable dorsal lateral/ventral/coagulating glands with typical intraluminal secretions. Portion of unremarkable vas deferens.
(85053, 85054)
No lesions of significance

Penis/Preputial gland

Typical penile structures including prepuce, glans, corpus cavernosum and urethra.
Typical preputial glands including basal and secretory cells.
(85055, 85056)
No lesions of significance

Urinary Bladder

Unremarkable bladder with typical urothelium and detrusor muscle.
(85053, 85054)
No lesions of significance

Liver/Gall bladder

Typical liver parenchyma including hepatocytes, Kupffer cells, portal triads and central veins.
Unremarkable gall bladder.
(85057)
No lesions of significance

Stomach

Unremarkable fore and glandular portions of the stomach with limiting ridge.
Includes pyloric sphincter and duodenal bulb.
(85058)
No lesions of significance

Small Intestine (Duodenum, Jejunum & Ileum)/GALT

Typical mucosal villi and submucosal layers.
Peyer’s patches display typical reactive nodal histology. Occasional, typical lymphoid cluster (cryptopatches).
(85058, 85059, 85060)
No lesions of significance

Cecum/Colon/GALT

Typical mucosal folds and submucosal layers. Unremarkable muscularis and discernible ganglion cells of the plexuses.
Occasional, typical lymphoid cluster (Peyer’s patch).
(85060)
No lesions of significance

Mesenteric lymph node

Typical reactive mesenteric lymph node with representative cortex including the occasional follicle, an expansive paracortical area with lymphocyte apoptosis, and sinus histiocytosis.
(85061)
No lesions of significance

Spleen

Mild follicular hyperplasia with germinal centre formation and some lymphocyte apoptosis. Mild coalescence of the white pulp.
Extramedullary haematopoiesis in the red pulp.
Haemosiderin laden macrophages identified in both the red and white pulp.
(85053, 85054)
No lesions of significance

Pancreas

Representative exocrine tissue (serous acini) and endocrine tissue (islets of Langerhans).
(85053, 85054)
No lesions of significance

Kidney

Sections show a cortex, medulla, and papilla. There is a uniform distribution of glomeruli and accompanying nephron components and the micromorphology of the tubules is unremarkable. Minimal multifocal peripelvic mononuclear cell infiltrates, incidental/age-related finding - feature seen in the controls.
Few vacuolated glomeruli, change considered too mild to be significant.
Few protein casts within the cortex and medulla regions, incidental/age related finding - feature seen in the controls.
Sections do not include renal lymph nodes. (85062)
No lesions of significance

Adrenal glands

Adrenal glands with typical cortex/medulla micromorphology.
(85062)
No lesions of significance

Salivary glands and Regional lymph nodes

Unremarkable submandibular, sublingual and parotid glands.
Minimal multifocal mononuclear cell infiltrates within the submandibular and parotid glands, an age-related change - feature seen in the controls.
Focal necrosis of the parotid gland, likely an incidental/spontaneous change (85061). Regional lymph nodes display typical reactive nodal histology.
(85061)
*Comments:*
*Pathology to comment*

Thyroids

Normal lateral lobes of the thyroid gland with typical colloid secreting follicles lined by cuboidal epithelium.
Small sheet-like mass of polygonal cells, characteristic of the parathyroid gland. (85063, 85064)
No lesions of significance

Trachea/Lungs

Typical lung parenchyma/alveoli, bronchioles, blood vessels and parabronchial lymph node.
Focal parenchymal congestion and collapse judged to be artefactual.
Trachea with unremarkable mucosal epithelial lining and hyaline cartilage. Oesophagus with typical features including stratified squamous epithelium.
Few small sized mononuclear cell aggegrates within the surrounding peri-adipose tissue, common finding - feature seen in the controls.
(85063, 85064)
No lesions of significance

Thymus

Typical medulla/cortex distribution and micromorphology.
Few small sized cysts (85064), common incidental/background finding - feature seen in the controls.
(85063, 85064)
No lesions of significance

Heart/chambers/vessels/valves

Mostly typical micromorphology observed in cardiac muscle, chambers, valves and great vessels of the heart.
Myxomatous valvular changes (thickened leaflets), incidental/age-related finding - feature seen in the controls.
Few foci of cardiomyocyte vacuolation and degeneration, likely incidental/age-related - feature seen in the controls.
Remaining cardiac muscle fibres demonstrated typical features including central nuclei, branching fibres and striations.
(85063, 85064)
*Comments:*
*Pathology to comment*

Skin

Typical dermal appendages and distribution. Unremarkable thin layer of striated muscle (panniculus carnosus).
Multifocal epidermal hyperplasia with underlying dermal infiltrates that are mainly neutrophilic, common incidental/background finding in mice.
(85065)
No lesions of significance

Tail

Typical tail components including keratinized squamous epithelium, dense regular connective tissue, tendons, caudal vertebra, bone marrow, intervertebral disc, skeletal muscle, nerves and blood vessels.
(85237, 85238, 85239)
No lesions of significance

Eyes/Harderian glands

Unremarkable retina, cornea, iris, ciliary body, lens, sclera and choroid.
Typical branched tubuloalveolar formation of the Harderian gland.
Unremarkable optic nerve and extraocular muscles. (85066)
No lesions of significance

Brain

Sections were prepared from the standard levels of the brain:
Level I forebrain: including cortex, corpus callosum, caudate putamen and lateral ventricles. Level II midbrain: including the hippocampus, thalamus, hypothalamus and lateral and third ventricles. Level III hindbrain: includes the cerebellum, pons and fourth ventricle.
Sections of brain appear symmetrical with unremarkable meninges and typical lamination. The cerebellum appears symmetrical with typical architecture and Purkinje cells. There was no obvious neuronal loss and the myelination appears normal.
Prominent flattening/stretching of the ependymal cells and increased size of the lateral ventricular space, features indicative of increased cerebrospinal fluid (hydrocephalus), a known background lesion in mice - similar feature seen in the controls. Query vacuolation of the adjacent white matter, possible artefact. (85067)
*Comments:*
*Neuropathology to comment*

Spinal cord

Representative thoracic and lumbar region of spinal cord, vertebral bone, intervertebral disc, striated muscle, peripheral nerves, brown adipose tissue, and bone marrow. Focal loss of the epidermis (ulcer) and multifocal inflammattion affecting mainly the subcutis. Segmental degeneration of the adajcent panniculus muscle (85243). (85241, 85242, 85243)
*Comments:*
*Neuropathology to comment*

(Hind leg) Long bone/Bone marrow/Synovial joint/Skeletal muscle

Representative long bone, striated muscle, examples of nerve fascicles, fibrocartilage of the meniscus, synovial joint and bone marrow. Focal mixed leucocyte inflammation within one digit of the foot (85237). Focal myodegeneration adjacent the hip joint (85237). Displays some features of regeneration. Considered to be incidental changes - features seen in the controls. Minimal focal perivascular eosinophlic infiltrate (85238, 85239). Query scattered necrotic myofibres (85238, 85239). Remaining skeletal muscle shows consistent fibre size with peripheral nuclei. (85237, 85238, 85239)
*Comments:*
*Pathology to comment*

Head

Multiple levels through the head demonstrate dermal appendages, nasal cavity, oral cavity, teeth and tongue including muscle bundles. Sections also show unremarkable pituitary gland including pars intermedia, pars distalis and pars nervosa as well and the trigeminal nerve/ganglia (85286). The outer and middle regions of the ear are discernible. The tympanic membrane is intact and the ossicles are unremarkable and include the stapedial annular ligaments (85287, 85288). Typical components of the inner ear including bony labyrinth, organ of corti, stria vascularis and scala cavities are discernible. Based on multiple levels, the organ of corti is unremarkable with no discernible loss of inner/outer hair cells and typical tectorial membrane (85287, 85288). The cochlear nerve and spiral ganglion is also demonstrated and based on several levels, there is no reduction in the density of the spiral ganglion cells. Examples of otolith organs can be seen with typical features such as the hair cells and mineral otoliths. The ampulla including the crista ridge with hair cells is discernible (85288).
(85286-85288)
No lesions of significance

Sternum

Representative sternebrae, costal cartilage, intersternebral joint, intercostal skeletal muscle and brown fat.
Haematopoietic tissue islands surrounded by vascular sinuses interspersed within a meshwork of trabecular bone. The bone marrow morphology demonstrated typical myeloid features including conspicuous megakaryoblasts and lymphoid features.
(85240)
No lesions of significance

#### #4

Macro Observations

Tail suspension test for neurological defects - negative.
Dentition, tongue and oral cavity was unremarkable.
BCS: 3
Spleen: 20×8×2mm
Kidneys: 14×8×5mm, symmetrical
Thymus: 7×7×2mm
Lungs inflated.
Heart: 12×10×6mm
Brain: 16×11×5mm, symmetrical
Pituitary gland identified, macroscopically normal
Tail length: 81mm (straight)
Head harvested for evaluation of auditory and vestibular structures.
Bone marrow smear taken from left hind leg.
No macroscopic lesions identified.

Micro Observations

Marrow smear: Examination of the smear showed representative cells from the myeloid and erythroid series. Occasional cells from the lymphoid series. Occasional and unremarkable megakaryoblasts. (85068)
Peripheral blood smear: Examination of the smear showed red blood cells (majority of cells shown), occasional white blood cells including lymphocytes, segmented neutrophils, monocytes and platelets (clumps). No discernible morphological changes or detectable parasites. (85069)

Mammary glands

Typical mammary fat pad with developing lactiferous ducts, blood vessels, and nerve bundles. No lesions of significance

Ovaries/Oviducts

Unremarkable ovaries containing follicles at various stages of development (primary through to antral) and several corpora lutea.
Few small to medium sized cysts in one ovary, common incidental/background finding in the mouse.
Mostly unremarkable oviduct micromorphology with typical columnar epithelium and mucosal folds.
Focal epithelial cytoplasmic vacuolation (85072), an age-related change - feature seen in the controls.
(85070, 85071, 85072)
No lesions of significance

Uterus/Cervix/Vagina/Clitoral gland

Cystic endometrial hyperplasia, an age-related change - feature seen in the controls.
Unremarkable myometrium and adventitia.
Minimal multifocal perivascular mononuclear cell infiltrates within the surrounding peri-adipose tissue, common background finding - feature seen in the controls.
The micromorphology of the uterus and vagina places the animal at estrus.
(85070, 85071, 85072)
No lesions of significance

Urinary Bladder

Portion of unremarkable detrusor muscle.
(85072)
No lesions of significance

Liver/Gall bladder

Liver parenchyma including hepatocytes, Kupffer cells, portal triads and central veins.
Multiple small foci of parenchymal inflammation and necrosis, common incidental/background finding - similar feature seen in the controls.
Unremarkable gall bladder.
(85073)
*Comments:*
*Pathology to comment*

Stomach

Unremarkable fore and glandular portions of the stomach with limiting ridge.
Includes pyloric sphincter and duodenal bulb.
(85074)
No lesions of significance

Small Intestine (Duodenum, Jejunum & Ileum)/GALT

Typical mucosal villi and submucosal layers.
Peyer’s patches display typical reactive nodal histology. Occasional, typical lymphoid cluster (cryptopatches).
(85074, 85075, 85076)
No lesions of significance

Cecum/Colon/GALT

Typical mucosal folds and submucosal layers. Unremarkable muscularis and discernible ganglion cells of the plexuses.
Occasional, typical lymphoid cluster (Peyer’s patch).
In mainly the cecum, abundant luminal protozoa - feature seen in the controls.
(85076)
No lesions of significance

Mesenteric lymph node

Highly reactive micromorphology including enlarged size, follicular hyperplasia with germinal centre formation, an expansive paracortical area containing lymphocyte apoptosis, and sinus histiocytosis.
(85077)
No lesions of significance

Spleen

Mild follicular hyperplasia with germinal centre formation and some lymphocyte apoptosis. Mild coalescence of the white pulp.
Expansive red pulp with marked extramedullary haematopoiesis. Haemosiderin laden macrophages identified in both the red and white pulp.
(85070, 85071, 85072)
No lesions of significance

Pancreas

Representative exocrine tissue (serous acini) and endocrine tissue (islets of Langerhans).
(85070, 85071, 85072)
No lesions of significance

Kidney

Sections show a cortex, medulla, and papilla. There is a uniform distribution of glomeruli and accompanying nephron components and the micromorphology of the tubules is unremarkable. Minimal focal perivascular/peripelvic mononuclear cell infiltrate, incidental/age-related change - feature seen in the controls.
Few mildly dilated protein casts within the medulla, incidental/age-related change - feature seen in the controls.
Renal lymph node with typical reactive nodal histology.
(85078)
No lesions of significance

Adrenal glands

Adrenal glands with typical cortex/medulla micromorphology.
(85078)
No lesions of significance

Salivary glands and Regional lymph nodes

Unremarkable submandibular, sublingual and parotid glands.
Mild multifocal perivascular mononuclear cell infiltrates within the submandibular gland, an age-related change - feature seen in the controls.
Regional lymph nodes display typical reactive nodal histology. (85077)
No lesions of significance

Thyroids

Normal lateral lobes of the thyroid gland with typical colloid secreting follicles lined by cuboidal epithelium.
Small sheet-like mass of polygonal cells, characteristic of the parathyroid gland.
(85079, 85080)
No lesions of significance

Trachea/Lungs

Typical lung parenchyma/alveoli, bronchioles, blood vessels and parabronchial lymph node.
Trachea with unremarkable mucosal epithelial lining and hyaline cartilage.
Oesophagus with typical features including stratified squamous epithelium.
Mild multifocal perivascular mononuclear cell infiltrates within the surrounding peri-adipose tissue, common finding - feature seen in the controls.
(85079, 85080)
No lesions of significance

Thymus

Typical medulla/cortex distribution and micromorphology.
(85079, 85080)
No lesions of significance

Heart/chambers/vessels/valves

Mostly typical micromorphology observed in cardiac muscle, chambers, valves and great vessels of the heart.
Myxomatous valvular changes (thickened leaflets) (85080), incidental/age-related finding - feature seen in the controls.
Occasioanal foci of cardiomycoyte vacuolation and degeneration, likely incidental/age-related - feature seen in the controls.
Remaining cardiac muscle fibres demonstrated typical features including central nuclei, branching fibres and striations.
(85079, 85080)
*Comments:*
*Pathology to comment*

Skin

Typical dermal appendages and distribution. Unremarkable thin layer of striated muscle (panniculus carnosus).
(85081)
No lesions of significance

Tail

Typical tail components including keratinized squamous epithelium, dense regular connective tissue, tendons, caudal vertebra, bone marrow, intervertebral disc, skeletal muscle, nerves and blood vessels.
(85255, 85256, 85257)
No lesions of significance

Eyes/Harderian glands

Unremarkable retina, cornea, iris, ciliary body, lens, sclera and choroid.
Typical branched tubuloalveolar formation of the Harderian gland.
Includes portion of unremarkable optic nerve and extraocular muscles.
(85082)
No lesions of significance

Brain

Sections were prepared from the standard levels of the brain:
Level I forebrain: including cortex, corpus callosum, caudate putamen and lateral ventricles. Level II midbrain: including the hippocampus, thalamus, hypothalamus and lateral and third ventricles. Level III hindbrain: includes the cerebellum, pons and fourth ventricle.
Sections of brain appear symmetrical with unremarkable meninges and typical lamination.
The cerebellum appears symmetrical with typical architecture and Purkinje cells.
There was no obvious neuronal loss and the myelination appears normal.
Some flattening/stretching of the ependymal cells, indicative of increased cerebrospinal fluid (hydrocephalus), a known background lesion in mice - similar feature seen in the controls.
Query vacuolation of the adjacent white matter, possible artefact.
(85083)
*Comments:*
*Neuropathology to comment*

Spinal cord

Representative thoracic and lumbar region of spinal cord, vertebral bone, intervertebral disc, striated muscle, peripheral nerves, brown adipose tissue, and bone marrow.
(85259, 85260)
No lesions of significance
*Comments:*
*Neuropathology to comment*

(Hind leg) Long bone/Bone marrow/Synovial joint/Skeletal muscle

Unremarkable long bone, striated muscle, examples of nerve fascicles, fibrocartilage of the meniscus, synovial joint and bone marrow. The skeletal muscle mostly shows consistent fibre size with peripheral nuclei.
Focal myodegeneration adjacent the hip joint, likely incidental - feature seen in the controls.
Marked focal mixed leukocyte inflammation of the foot (85257),, likely incidental - feature seen in the controls.
(85255, 85256, 85257)
*Comments:*
*Pathology to comment*

Head

Multiple levels through the head demonstrate dermal appendages, nasal cavity, oral cavity, teeth and tongue including muscle bundles. Sections also show unremarkable pituitary gland including pars intermedia, pars distalis and pars nervosa as well and the trigeminal nerve/ganglia (85289). The outer and middle regions of the ear are discernible. The tympanic membrane is intact and the ossicles are unremarkable and include the stapedial annular ligaments (85290, 85291). Typical components of the inner ear including bony labyrinth, organ of corti, stria vascularis and scala cavities are discernible. Based on multiple levels, the organ of corti is unremarkable with no discernible loss of inner/outer hair cells and typical tectorial membrane (85290, 85291). The cochlear nerve and spiral ganglion is also demonstrated and based on several levels, there is no reduction in the density of the spiral ganglion cells. Examples of otolith organs can be seen with typical features such as the hair cells and mineral otoliths. The ampulla including the crista ridge with hair cells is discernible (85291).
Inflammation of the middle ear characterised by oedema, luminal cellular debris, and some formation of cholesterol clefts (85291), unilateral - a known incidental/background finding in mice.
Few foci of cytoplasmic hyaline droplet accumulation within the nasal epithelium (85290, 85291), incidental - feature seen in the controls.
(85289-85291)
*Comments:*
*Pathology to comment*

Sternum

Representative sternebrae, costal cartilage, intersternebral joint, intercostal skeletal muscle and brown fat.
Haematopoietic tissue islands surrounded by vascular sinuses interspersed within a meshwork of trabecular bone. The bone marrow morphology demonstrated typical myeloid features including conspicuous megakaryoblasts and lymphoid features.
(85258)
No lesions of significance

#### #7

Macro Observations

Tail suspension test for neurological defects - negative.
Dentition, tongue and oral cavity was unremarkable.
BCS: 3
Spleen: 20×8×2mm
Kidneys: 14×8×5mm, symmetrical (left kidney bisected coronally)
Thymus: 7×7×2mm
Lungs inflated.
Heart: 12×10×6mm
Brain: 16×10×6mm, symmetrical
Pituitary gland identified, macroscopically normal Tail length: 82mm (straight)
Head harvested for evaluation of auditory and vestibular structures. Bone marrow smear taken from left hind leg.
No macroscopic lesions identified.

Micro Observations

Marrow smear: Examination of the smear showed representative cells from the myeloid and erythroid series. Occasional cells from the lymphoid series. Occasional and unremarkable megakaryoblasts.
(85114)
Peripheral blood smear: Examination of the smear showed red blood cells (majority of cells shown), occasional white blood cells including lymphocytes, segmented neutrophils, monocytes and platelets (clumps). No discernible morphological changes or detectable parasites.
(85115)

Mammary glands

Typical mammary fat pad with developing lactiferous ducts, blood vessels, and nerve bundles. Minimal multifocal perivascular mononuclear cell infilltrates (85128), incidental/age-related - feature seen in the controls.
(85128)
No lesions of significance

Ovaries/Oviducts

Unremarkable ovaries containing follicles at various stages of development (primary through to antral) and several corpora lutea.
Moderate mononuclear cell infiltration of the bursa membrane, bilateral, an age-related change in mice.
Mostly unremarkable oviduct micromorphology with typical columnar epithelium and mucosal folds.
Focal epithelial cytoplasmic vacuolation, bilateral, an age-related change - feature seen in the controls.
Mild multifocal perivascular mononuclear cell infiltrates in the surrounding peri-adipose tissue, an age-related change - feature seen in the controls.
(85116-85119)
No lesions of significance

Uterus/Cervix/Vagina/Clitoral gland

Endometrial hyperplasia, an age-related change - feature seen in the controls.\Unremarkable architecture of the myometrium and adventitia.\The micromorphology of the uterus and vagina places the animal at metestrus.
(85116-85119)
No lesions of significance
Urinary Bladder
Portion of unremarkable bladder with typical urothelium and detrusor muscle.
(85119)
No lesions of significance

Liver/Gall bladder

Liver parenchyma including hepatocytes, Kupffer cells, portal triads and central veins.
Minimal multifocal perivascular mononuclear cell infiltrates, an incidental/age-related change - feature seen in the controls.
Multiple small parenchymal leucocyte aggregates, and focal parenchymal necrosis, common incidental/background findings in mice - similar feature seen in the controls.
Several small foci of hepatocyte vacuolation resembling fatty change (steatosis), an incidental/age-related finding in mice.
Unremarkable gall bladder.
(85120)
*Comments:*
*Pathology to comment*

Stomach

Unremarkable fore portion of the stomach.
Of the glandular portion, marked and mainly submucosal eosinophilic inflammatoin (transmural), common incidental finding in mice.
Sections includes pyloric sphincter and duodenal bulb.
(85121)

Small Intestine (Duodenum, Jejunum & Ileum)/GALT\

Mostly typical mucosal villi and submucosal layers.
Marked and mainly eosinophilc infiltration of the distal ileum (transmural) (85122), likely an incidental finding.
Peyer’s patches display typical reactive nodal histology. Occasional, typical lymphoid cluster (cryptopatches).
(85121, 85131, 85122)

Cecum/Colon/GALT

Typical mucosal folds and submucosal layers. Unremarkable muscularis and discernible ganglion cells of the plexuses.
Occasional, typical lymphoid cluster (Peyer’s patch).
Abundant luminal protozoa in the cecum and proximal colon - feature seen in the controls. (85123)
No lesions of significance

Mesenteric lymph node

Highly reactive micromorphology including enlarged size, follicular hyperplasia with germinal centre formation, an expansive paracortical area, scattered lymphocyte apoptosis, and sinus histiocytosis.
(85124)
No lesions of significance

Spleen

Mild follicular hyperplasia with germinal centre formation and lymphocyte apoptosis. Mild coalescence of the white pulp.
Expansive red pulp with marked extramedullary haematopoiesis.
(85116-85119)
No lesions of significance

Pancreas

Representative exocrine tissue (serous acini) and endocrine tissue (islets of Langerhans).
(85116-85119)
No lesions of significance

Kidney

Sections show a cortex, medulla, and papilla. There is a uniform distribution of glomeruli and accompanying nephron components and the micromorphology of the tubules is unremarkable. Renal lymph nodes with typical reactive nodal histology.
(85120, 85125)
No lesions of significance

Adrenal glands

Adrenal glands with typical cortex/medulla micromorphology.
(85120, 85125)
No lesions of significance

Salivary glands and Regional lymph nodes

Mild multifocal perivascular mononuclear cell infiltrates within the submandibular gland, an age-related change - feature seen in the controls.
Unremarkable sublingual and parotid glands.
Regional lymph nodes display typical reactive nodal histology. (85124)
No lesions of significance

Thyroids

Normal lateral lobes of the thyroid gland with typical colloid secreting follicles lined by cuboidal epithelium.
(85126, 85127)
No lesions of significance

Trachea/Lungs

Typical lung parenchyma/alveoli, bronchioles, and blood vessels.
Trachea with unremarkable mucosal epithelial lining and hyaline cartilage. Oesophagus with typical features including stratified squamous epithelium.
Mild multifocal perivascular mononuclear cell infiltrates in the surrounding periadipose tissue, common background finding - feature seen in the controls.
(85126, 85127)
No lesions of significance

Thymus

Typical medulla/cortex distribution and micromorphology.
Single small sized cyst (85216), common incidental/background finding - feature seen in the controls.
(85126, 85127)
No lesions of significance

Heart/chambers/vessels/valves

Representative cardiac muscle, chambers, valves and great vessels of the heart.
Some myxomatous valvular changes (thickened leaflets) (85126), an age-related change - feature seen in the controls.
The cardiac muscle fibres demonstrated typical features including central nuclei, branching fibres and striations.
(85126, 85127)
No lesions of significance

Skin

Typical dermal appendages and distribution. Unremarkable thin layer of striated muscle
(panniculus carnosus).
(85128)
No lesions of significance

Tail

Typical tail components including keratinized squamous epithelium, dense regular connective tissue, tendons, caudal vertebra, bone marrow, intervertebral disc, skeletal muscle, nerves and blood vessels.
(85250, 85251)
No lesions of significance

Eyes/Harderian glands

Unremarkable retina, cornea, iris, ciliary body, lens, sclera and choroid.
Typical branched tubuloalveolar formation of the Harderian gland.
Includes portion of unremarkable optic nerve and extraocular muscles. (85129)
No lesions of significance

Brain

Sections were prepared from the standard levels of the brain:
Level I forebrain: including cortex, corpus callosum, caudate putamen and lateral ventricles. Level II midbrain: including the hippocampus, thalamus, hypothalamus and lateral and third ventricles. Level III hindbrain: includes the cerebellum, pons and fourth ventricle.
Sections of brain appear symmetrical with unremarkable meninges and typical lamination. The cerebellum appears symmetrical with typical architecture and Purkinje cells. There was no obvious neuronal loss and the myelination appears normal.
Prominent flattening/stretching of the ependymal cells and increased size of the lateral ventricular space, features indicative of increased cerebrospinal fluid (hydrocephalus), a known background lesion in mice - similar feature seen in the controls.
(85130)
*Comments:*
*Neuropathology to comment*

Spinal cord

Representative thoracic and lumbar region of spinal cord, vertebral bone, intervertebral disc, striated muscle, peripheral nerves, brown adipose tissue, and bone marrow.
(85253, 85254)
No lesions of significance
*Comments:*
*Neuropathology to comment*

(Hind leg) Long bone/Bone marrow/Synovial joint/Skeletal muscle

Mostly unremarkable long bone, striated muscle, examples of nerve fascicles, fibrocartilage of the meniscus, synovial joint and bone marrow. The skeletal muscle shows consistent fibre size with peripheral nuclei.
Focal mixed leucocyte inflammation surrounding a tendon of the distal leg, likely incidental - similar feature seen in the controls.
(85250, 85251)
*Comments:*
*Pathology to comment*

Head

Multiple levels through the head demonstrate dermal appendages, nasal cavity, oral cavity, teeth and tongue including muscle bundles. Sections also show unremarkable pituitary gland including pars intermedia, pars distalis and pars nervosa as well and the trigeminal nerve/ganglia (85297). The outer and middle regions of the ear are discernible. The tympanic membrane is intact and the ossicles are unremarkable and include the stapedial annular ligaments (85298, 85299). Typical components of the inner ear including bony labyrinth, organ of corti, stria vascularis and scala cavities are discernible. Based on multiple levels, the organ of corti is unremarkable with no discernible loss of inner/outer hair cells and typical tectorial membrane (85298, 85299). The cochlear nerve and spiral ganglion is also demonstrated and based on several levels, there is no reduction in the density of the spiral ganglion cells. Examples of otolith organs can be seen with typical features such as the hair cells and mineral otoliths. The ampulla including the crista ridge with hair cells is discernible (85299).
(85297-85299)
No lesions of significance

Sternum

Representative sternebrae, costal cartilage, intersternebral joint, intercostal skeletal muscle and brown fat.
Haematopoietic tissue islands surrounded by vascular sinuses interspersed within a meshwork of trabecular bone. The bone marrow morphology demonstrated typical myeloid features including conspicuous megakaryoblasts and lymphoid features.
(85252)
No lesions of significance

#### #8

Macro Observations

Tail suspension test for neurological defects - negative.
Dentition, tongue and oral cavity was unremarkable.
BCS: 3
Spleen: 14×5×3mm
Kidneys: not measured, symmetrical
Thymus: 8×8×2mm
Lungs inflated.
Heart: 9×7×5mm
Brain: 15×10×5mm, symmetrical
Pituitary gland identified, macroscopically normal
Tail length: 80mm (straight)
Head harvested for evaluation of auditory and vestibular structures. Bone marrow smear taken from left hind leg.
No macroscopic lesions identified.

Micro Observations

Marrow smear: Examination of the smear showed representative cells from the myeloid and erythroid series. Occasional cells from the lymphoid series. Occasional and unremarkable megakaryoblasts.
(85132)
Peripheral blood smear: Examination of the smear showed red blood cells (majority of cells shown), occasional white blood cells including lymphocytes, segmented neutrophils, monocytes and platelets (clumps). No discernible morphological changes or detectable parasites.
(85133)

Mammary glands

Typical mammary fat pad with developing lactiferous ducts, blood vessels, and nerve bundles. (85141, 85145)
No lesions of significance

Ovaries/Oviducts

Unremarkable ovaries containing follicles at various stages of development (primary through to antral) and several corpora lutea.
Few pigment laden interstitial cells (likely ceroid-lipofucsin), an age-related change - feature seen in the controls.
Mononuclear cell infiltration of the bursa membrane, bilateral, an age-related finding in mice.
Mostly unremarkable oviduct micromorphology with typical columnar epithelium and mucosal folds.
Focal epithelial cytoplasmic vacuolation, an age-related change - feature seen in the controls. (85134, 85135, 85136)
No lesions of significance

Uterus/Cervix/Vagina/Clitoral gland

Cystic endometrial hyperplasia, an age-related change - feature seen in the controls. Unremarkable architecture myometrium and adventitia.
The micromorphology of the uterus and vagina places the animal at estrus.
(85134, 85135, 85136)
No lesions of significance

Urinary Bladder

Unremarkable bladder with typical urothelium and detrusor muscle. (85134, 85135, 85136)
No lesions of significance

Liver/Gall bladder

Liver parenchyma including hepatocytes, Kupffer cells, portal triads and central veins.
Multiple small parenchymal leukocyte aggregates with some hepatocyte cell loss/necrosis - common incidental/background finding - feature seen in the controls.
Mild multifocal perivascular mononuclear cell infiltrates, an incidental/age-related change - feature seen in the controls.
Gall bladder with epithelial hyperplasia, cytoplasmic hyaline droplet accumulation, and underlying neutrophils in the lamina propria, common incidental/background finding - feature seen in the controls.
(85137)
No lesions of significance

Stomach

Unremarkable fore and glandular portions of the stomach with limiting ridge.
Includes pyloric sphincter and duodenal bulb.
(85138)
No lesions of significance

Small Intestine (Duodenum, Jejunum & Ileum)/GALT

Typical mucosal villi and submucosal layers.
Peyer’s patches display typical reactive nodal histology. Occasional, typical lymphoid cluster (cryptopatches).
(85138, 85139, 85140)
No lesions of significance

Cecum/Colon/GALT

Typical mucosal folds and submucosal layers. Unremarkable muscularis and discernible ganglion cells of the plexuses.
Occasional, typical lymphoid cluster (Peyer’s patch).
Clusters of luminal protozoa in the cecum - feature seen in the controls.
Increased mononuclear cell infiltrates in the lamina of the proximal colon, incidental - feature seen in the controls.
(85140)
No lesions of significance

Mesenteric lymph node

Typical reactive nodal histology with representative cortex including the occasional follicle, an expansive paracortical area with lymphocyte apoptosis, and sinus histiocytosis.
(85141)
No lesions of significance

Spleen

Mild follicular hyperplasia with germinal centre formation and some lymphocyte apoptosis. Mild coalescence of the white pulp.
Expansive red pulp with marked extramedullary haematopoiesis.
(85136)
No lesions of significance

Pancreas

Representative exocrine tissue (serous acini) and endocrine tissue (islets of Langerhans).
(85134, 85135, 85136)
No lesions of significance

Kidney

Sections show a cortex, medulla, and papilla. There is a uniform distribution of glomeruli and accompanying nephron components and the micromorphology of the tubules is unremarkable. Minimal perivascular/peripelvic mononuclear cell infiltrates, incidental/age-related change - feature seen in the controls.
Several mildly dilated protein casts within the medulla, incidental/age-related change - feature seen in the controls.
Renal lymph nodes with typical reactive nodal histology and mild sinus histiocytosis. (85142)
No lesions of significance

Adrenal glands

Adrenal glands with typical cortex/medulla micromorphology.
(85142
No lesions of significance

Salivary glands and Regional lymph nodes

Mild multifocal perivascular mononuclear cell infiltrates within the submandibular gland, an age-related change - feature seen in the controls.
Unremarkable sublingual and parotid glands.
Regional lymph nodes display typical reactive nodal histology. (85141)
No lesions of significance

Thyroids

Normal lateral lobes of the thyroid gland with typical colloid secreting follicles lined by cuboidal epithelium.
(85143, 85144)
No lesions of significance

Trachea/Lungs

Typical lung parenchyma/alveoli, bronchioles, and blood vessels.
Focal parenchymal congestion and collapse judged to be artefactual.
Mild multifocal perivascular mononuclear cell infiltrates, an incidental/age-related change - feature seen in the controls.
Trachea with unremarkable mucosal epithelial lining and hyaline cartilage.
Oesophagus with typical features including stratified squamous epithelium. (85143, 85144)
No lesions of significance

Thymus

Typical medulla/cortex distribution and micromorphology.
(85143, 85144)
No lesions of significance

Heart/chambers/vessels/valves

Mostly typical micromorphology observed in cardiac muscle, chambers, valves and great vessels of the heart.
Myxomatous valvular changes (thickened leaflets), an age-related change - feature seen in the controls.
The cardiac muscle fibres demonstrated typical features including central nuclei, branching fibres and striations.
Focal basophilic deposit in the ventricular muscle resembling mineralisation, likely incidental.
(85143, 85144)
*Comments:*
*Pathology to comment*

Skin

Typical dermal appendages and distribution. Unremarkable thin layer of striated muscle
(panniculus carnosus).
(85145)
No lesions of significance

Tail

Typical tail components including keratinized squamous epithelium, dense regular connective tissue, tendons, caudal vertebra, bone marrow, intervertebral disc, skeletal muscle, nerves and blood vessels.
(85224, 85225, 85226)
No lesions of significance

Eyes/Harderian glands

Unremarkable retina, cornea, iris, ciliary body, lens, sclera and choroid.
Typical branched tubuloalveolar formation of the Harderian gland.
Minimal multifocal glandular mononuclear cell infiltrates - an incidental/age-related finding in mice.
Includes portion of unremarkable optic nerve and extraocular muscles.
(85146)
No lesions of significance

Brain

Sections were prepared from the standard levels of the brain:
Level I forebrain: including cortex, corpus callosum, caudate putamen and lateral ventricles. Level II midbrain: including the hippocampus, thalamus, hypothalamus and lateral and third ventricles. Level III hindbrain: includes the cerebellum, pons and fourth ventricle.
Sections of brain appear symmetrical with unremarkable meninges and typical lamination. The cerebellum appears symmetrical with typical architecture and Purkinje cells. There was no obvious neuronal loss and the myelination appears normal.
Prominent flattening/stretching of the ependymal cells and increased size of the lateral ventricular space, features indicative of increased cerebrospinal fluid (hydrocephalus), a known background lesion in mice - similar feature seen in the controls. Query vacuolation of the adjacent white matter, possible artefact. (85147)
*Comments:*
*Neuropathology to comment*

Spinal cord

Representative thoracic and lumbar region of spinal cord, vertebral bone, intervertebral disc, striated muscle, peripheral nerves, brown adipose tissue, and bone marrow.
(85227, 85228, 85229)
No lesions of significance
*Comments:*
*Neuropathology to comment*

(Hind leg) Long bone/Bone marrow/Synovial joint/Skeletal muscle

Unremarkable long bone, striated muscle, examples of nerve fascicles, fibrocartilage of the meniscus, synovial joint and bone marrow. The skeletal muscle shows consistent fibre size with peripheral nuclei.
(85224, 85225, 85226)
No lesions of significance

Head

Multiple levels through the head demonstrate dermal appendages, nasal cavity, oral cavity, teeth and tongue including muscle bundles. Sections also show unremarkable pituitary gland including pars intermedia, pars distalis and pars nervosa as well and the trigeminal nerve/ganglia (85300). The outer and middle regions of the ear are discernible. The tympanic membrane is intact and the ossicles are unremarkable and include the stapedial annular ligaments (85301, 85302). Typical components of the inner ear including bony labyrinth, organ of corti, stria vascularis and scala cavities are discernible. Based on multiple levels, the organ of corti is unremarkable with no discernible loss of inner/outer hair cells and typical tectorial membrane (85301, 85302). The cochlear nerve and spiral ganglion is also demonstrated and based on several levels, there is no reduction in the density of the spiral ganglion cells. Examples of otolith organs can be seen with typical features such as the hair cells and mineral otoliths. The ampulla including the crista ridge with hair cells is discernible (85302).
Segmental hyaline droplet accumulation within the nasal epithelium, incidental - feature seen in the controls.
(85300-85302) No lesions of significance

Sternum

Representative sternebrae, costal cartilage, intersternebral joint, intercostal skeletal muscle and brown fat.
Haematopoietic tissue islands surrounded by vascular sinuses interspersed within a meshwork of trabecular bone. The bone marrow morphology demonstrated typical myeloid features including conspicuous megakaryoblasts and lymphoid features. (85230)
No lesions of significance

#### Comment / Plan

Case APN21/036SVI (C. Walkley) will be referred to Dr. John Finnie, University of Adelaide for comment.
Aira Nuguid
18th October, 2021

### Supplementary Pathology Report

#### #20 (control)

85203 – Pancreas – focal necrosis of exocrine pancreas with lymphocytic infiltration.
85207 – Heart – cardiomyocyte degeneration with cytoplasmic vacuolation.
85210 – Eye – apparent focal loss of corneal epithelium (? traumatic aetiology).
85281 – Hind leg – focal skeletal myocyte degeneration and necrosis.

#### #5 (control)

85088 – Liver – multifocal hepatocellular necrosis with mononuclear cell reaction, likely bacterial, and periportal lymphocytic infiltration, both common incidental findings.
85089 – Pancreas – focal necrosis of exocrine pancreas with lymphocytic infiltration.
85094 – Heart – multifocal cardiomyocyte degeneration and necrosis and lymphocytic infiltration.

#### #6 (control)

85107 – Salivary gland – acinar atrophy.
85295 – Head/external ear – focal coagulation necrosis of lining epithelium.
85234 – Hind leg – marked subcutaneous oedema (anasarca).
85235 – Hind leg – subcutaneous degeneration with mast cell hyperplasia and eosinophil infiltration (? allergic aetiology).

#### #11

85152 – Liver – mild diffuse cloudy swelling and hydropic degeneration of hepatocytes.
85269 – Hind leg – focal skeletal myocyte degeneration and necrosis.

#### #12

85167 – Liver – mild diffuse cloudy swelling and hydropic degeneration of hepatocytes.

#### #18

85273 – Hind leg – necrotic osteolysis (? traumatic aetiology).

#### #3

85061 – Salivary gland – large focus of acute acinar necrosis
85063 – Heart – multifocal cardiomyocyte degeneration and necrosis.
85237 – Hind leg – focal skeletal myocyte degeneration and regeneration.=
85239 – Hind leg – focal coagulation necrosis of skeletal myocytes.
85243 – Skin – acute focal epidermal ulceration and degeneration and necrosis of panniculus muscle.

#### #4

85073 – Liver – multifocal hepatocellular necrosis with mononuclear cell reaction, likely bacterial, and periportal lymphocytic infiltration, both common incidental findings.
85079 – Heart – multifocal cardiomyocyte degeneration and necrosis.
85291 – Head/middle ear – luminal macrophages and multinucleated giant cells with cholesterol clefts in wall.
85257 – Hind leg – skeletal myocyte degeneration and necrosis.
85120 – Liver – multifocal hepatocellular necrosis with mononuclear cell reaction, likely bacterial, and periportal lymphocytic infiltration, both common incidental findings.
85120 – Liver –focal micro and macro vesicular steatosis of hepatocytes. 85251 – Hind leg – apparent cellulitis with mixed inflammatory cell infiltration.

#### #8

85144 – Heart – focal mineralization.

#### Summary

Please see individual animals in the above report for comments.
Note: NAD = No abnormalities detected.
Pathologist name and details removed CW230222 27th October, 2021

### Supplementary Neuropathology Report

#### #20 (control)

85211 – Brain – NAD.
85285 – Spinal cord – NAD.

#### #5 (control)

85098 – Brain – variable dilation of lateral ventricles (internal hydrocephalus).
85246 – Spinal cord – NAD.

#### #6 (control)

85113 – Brain – variable dilation of lateral ventricles (internal hydrocephalus).
85232 – Spinal cord – NAD.

#### #11

85162 – Brain – variable dilation of lateral ventricles (internal hydrocephalus).
85272 – Spinal cord – NAD.

#### #12

85177 – Brain – variable dilation of lateral ventricles (internal hydrocephalus).
85266 – Spinal cord – NAD.
85194 – Brain – variable dilation of lateral ventricles (internal hydrocephalus).
85278 – Spinal cord – NAD.

#### #3

85067 – Brain – variable dilation of lateral ventricles (internal hydrocephalus) and oedema of surrounding white matter tracts.
85343 – Spinal cord – NAD.

#### #4

85083 – Brain – variable dilation of lateral ventricles (internal hydrocephalus).
85260 – Spinal cord – NAD.

#### #7

85130 – Brain – variable dilation of lateral ventricles (internal hydrocephalus).
85254 – Spinal cord – NAD.

#### #8

85147 – Brain – variable dilation of lateral ventricles (internal hydrocephalus) and oedema of surrounding white matter tracts.
85229 – Spinal cord – NAD.

#### Summary

Brains – #4, #5, #6, #7, #11, #12 and #18 – variable dilation of lateral ventricles (internal hydrocephalus).
Brains – #3, #8 – variable dilation of lateral ventricles (internal hydrocephalus) and oedema of surrounding white matter tracts.
Sections of the spinal cord show no significant findings.
Note: NAD = No abnormalities detected.
(Pathologist name and contact details removed_CW23022) 27th October, 2021
Phenomics Australia advises all research groups that images or results obtained are to be acknowledged in resultant publications. Example acknowledgement: “This study utilised Phenomics Australia Histopathology and Slide Scanning Service, University of Melbourne”.
Authorised by (name removed_CW230222), Phenomics Australia and Slide Scanning Service manager.

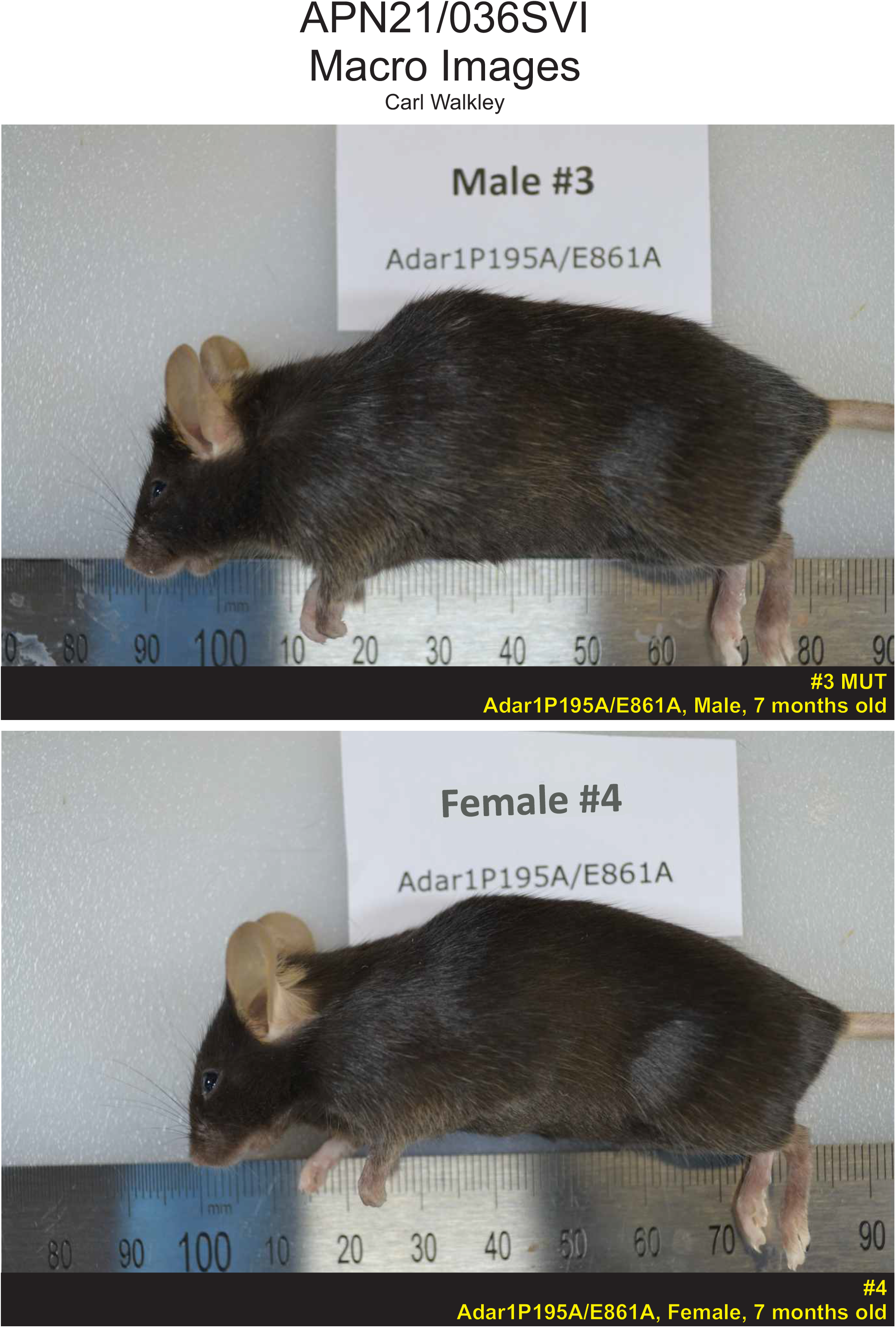
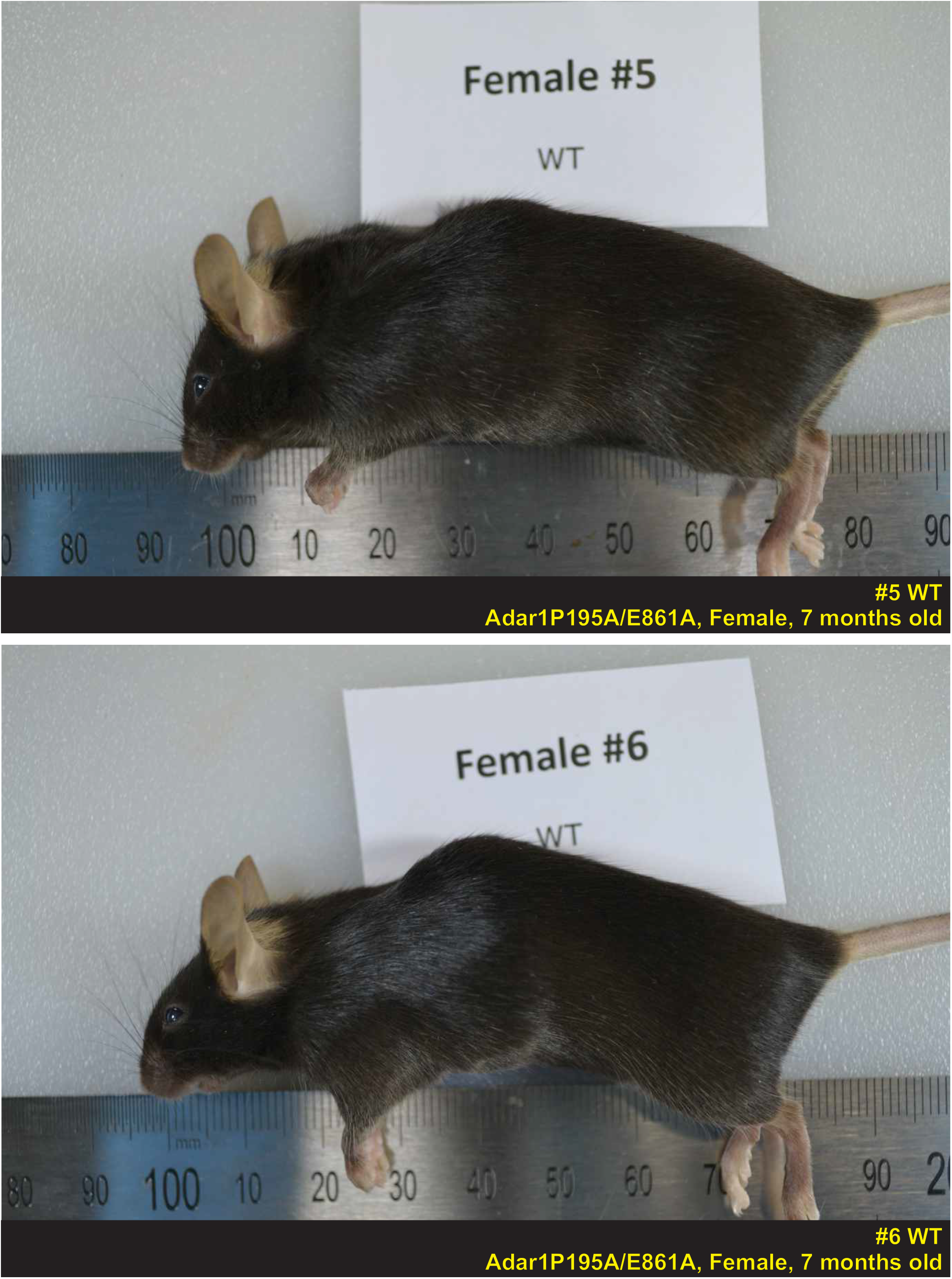
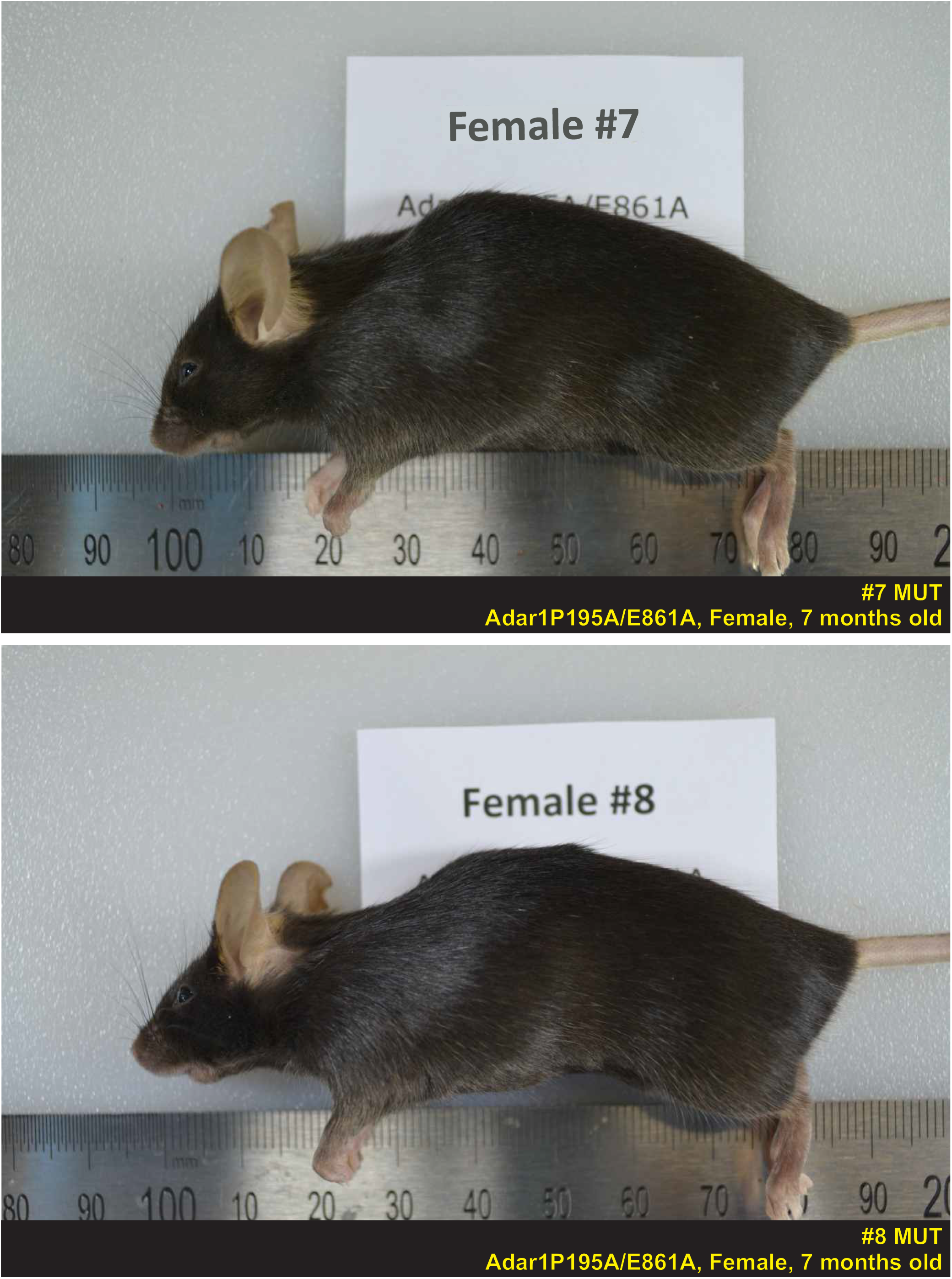
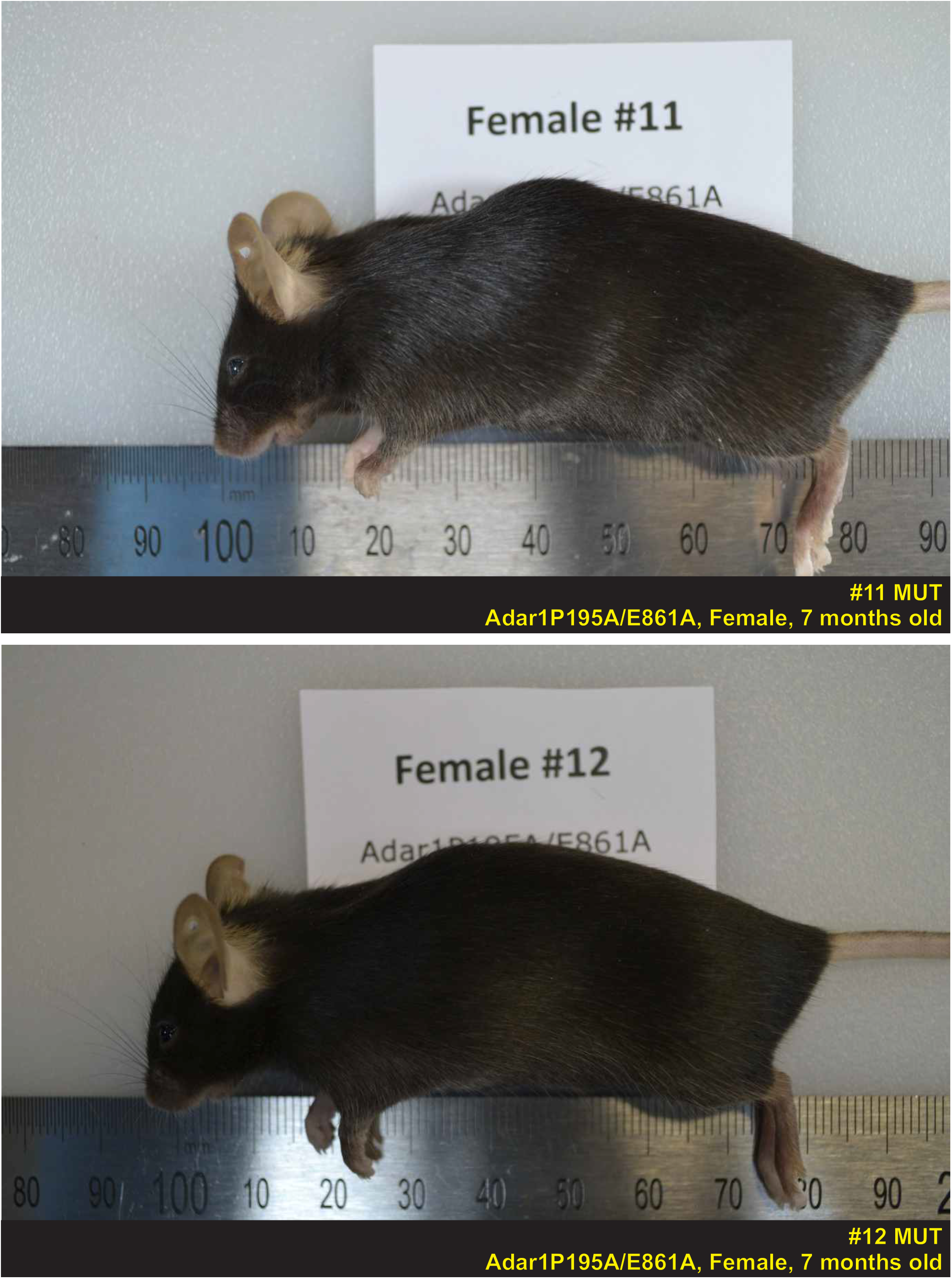
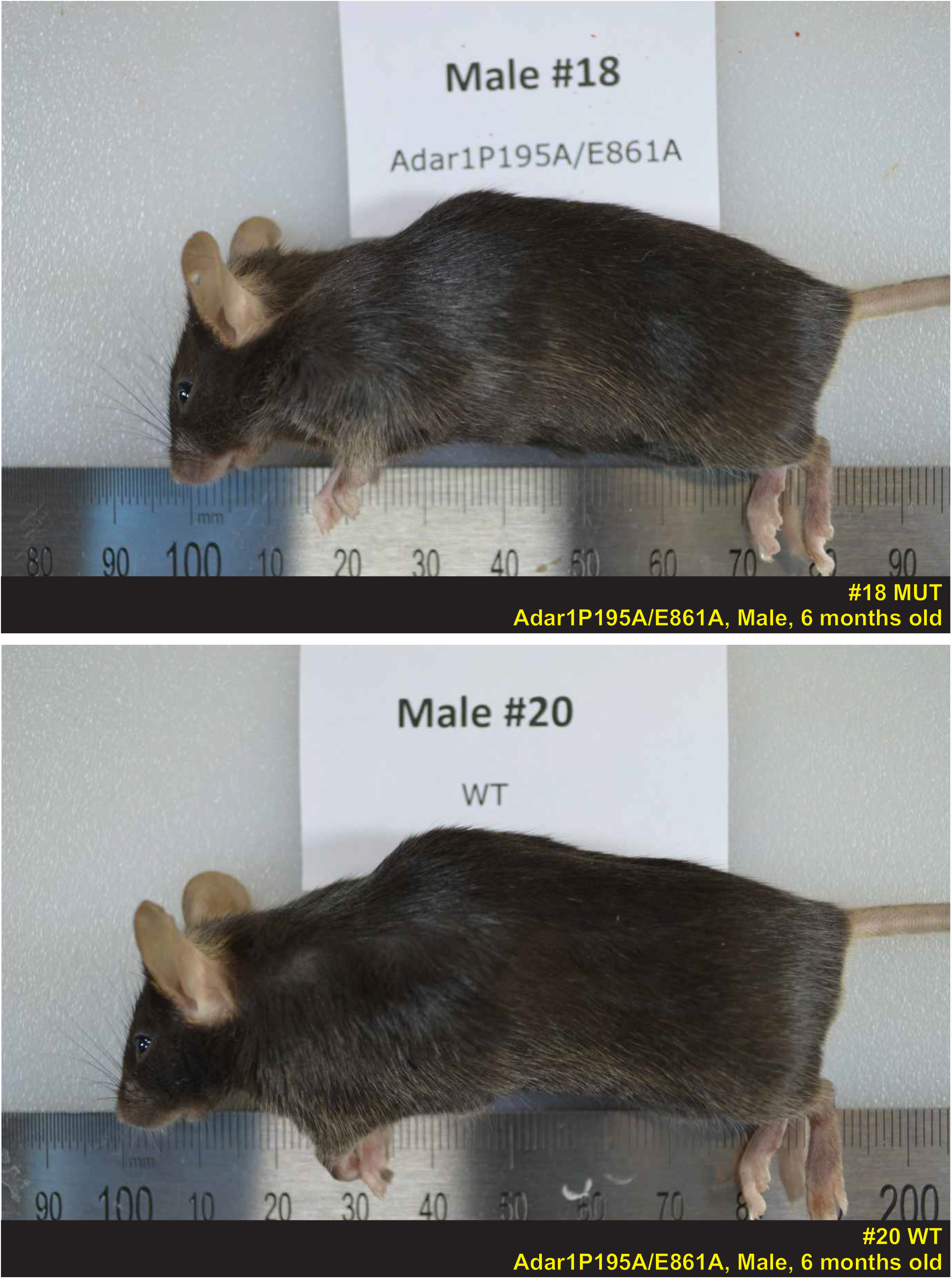

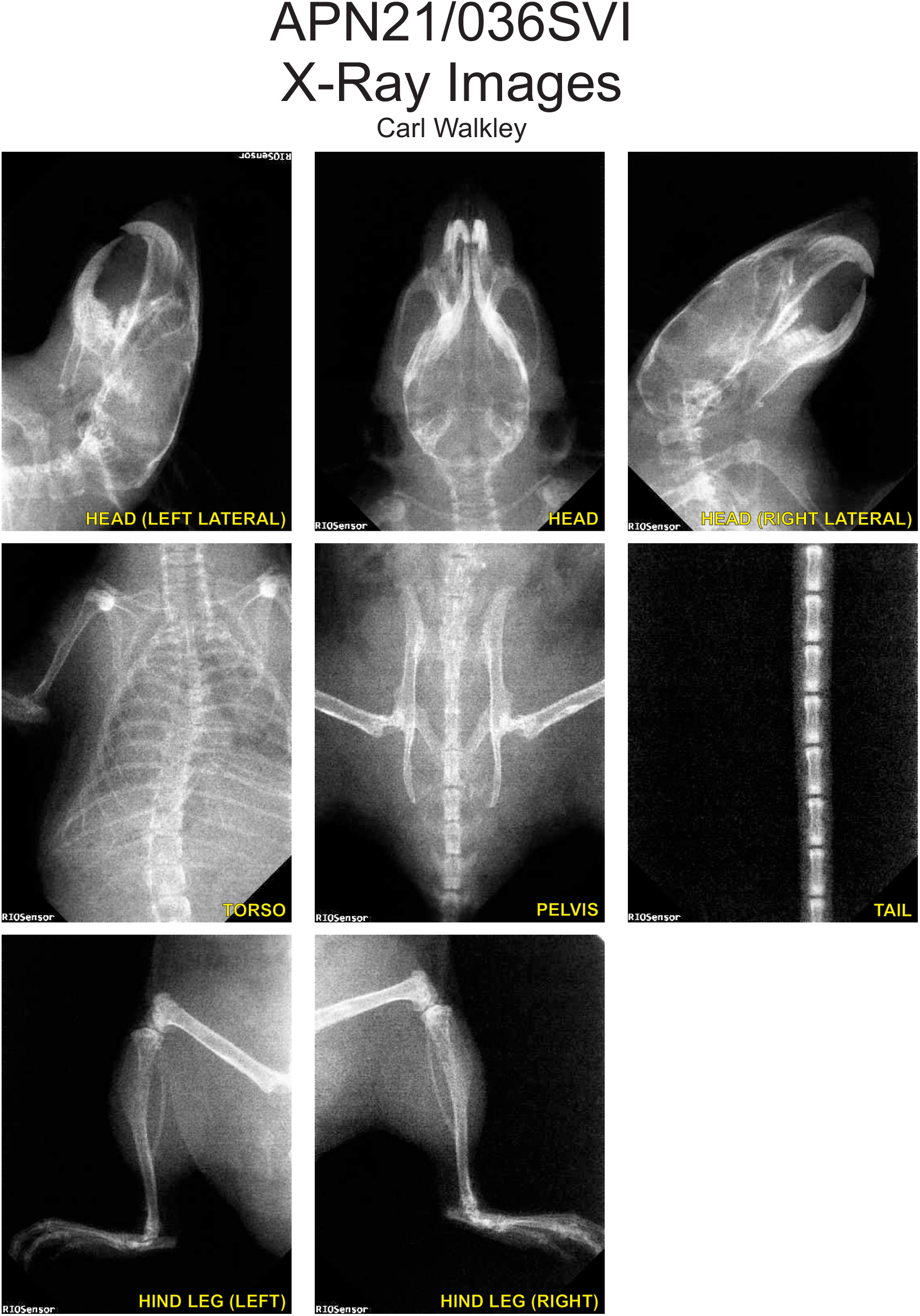
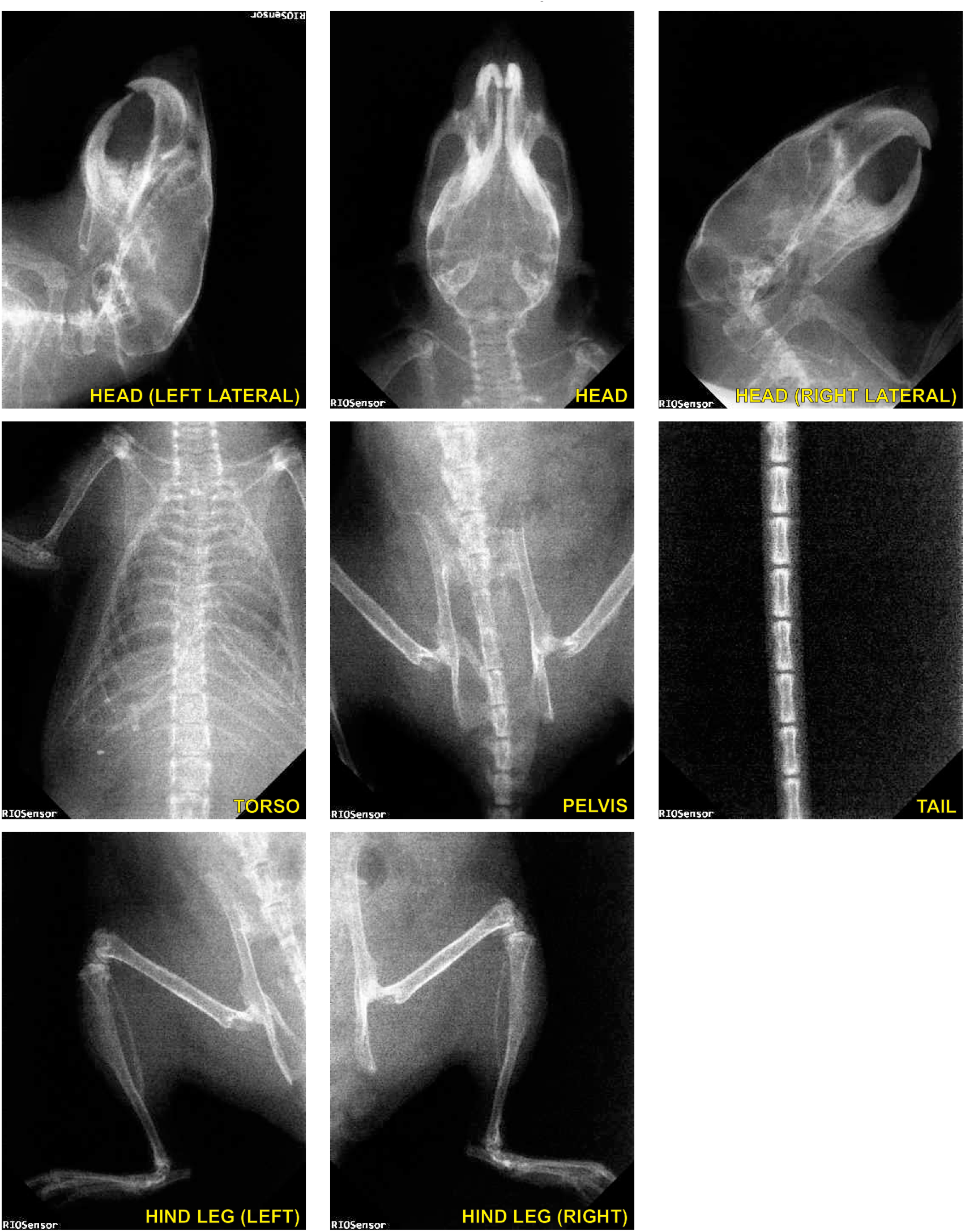
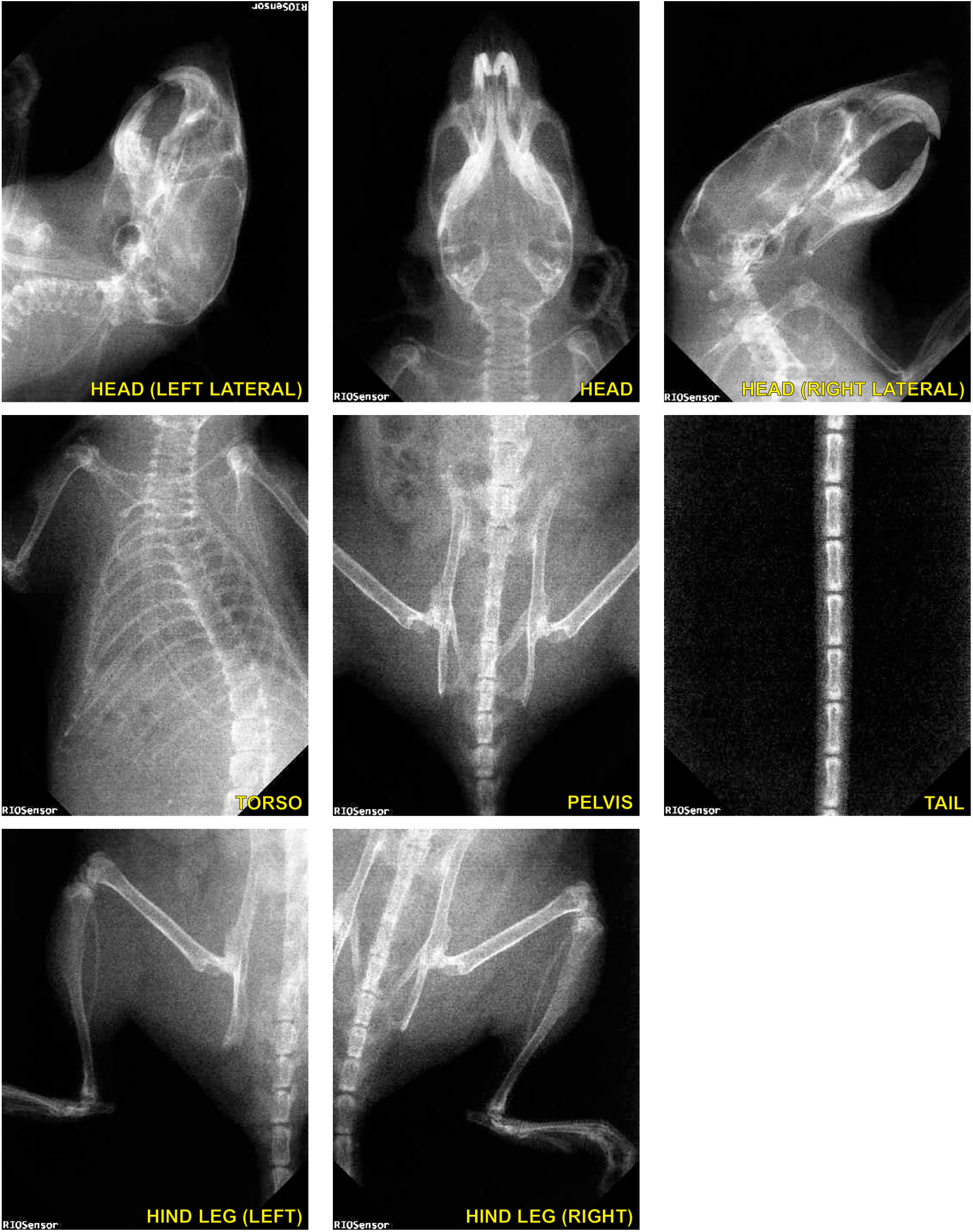
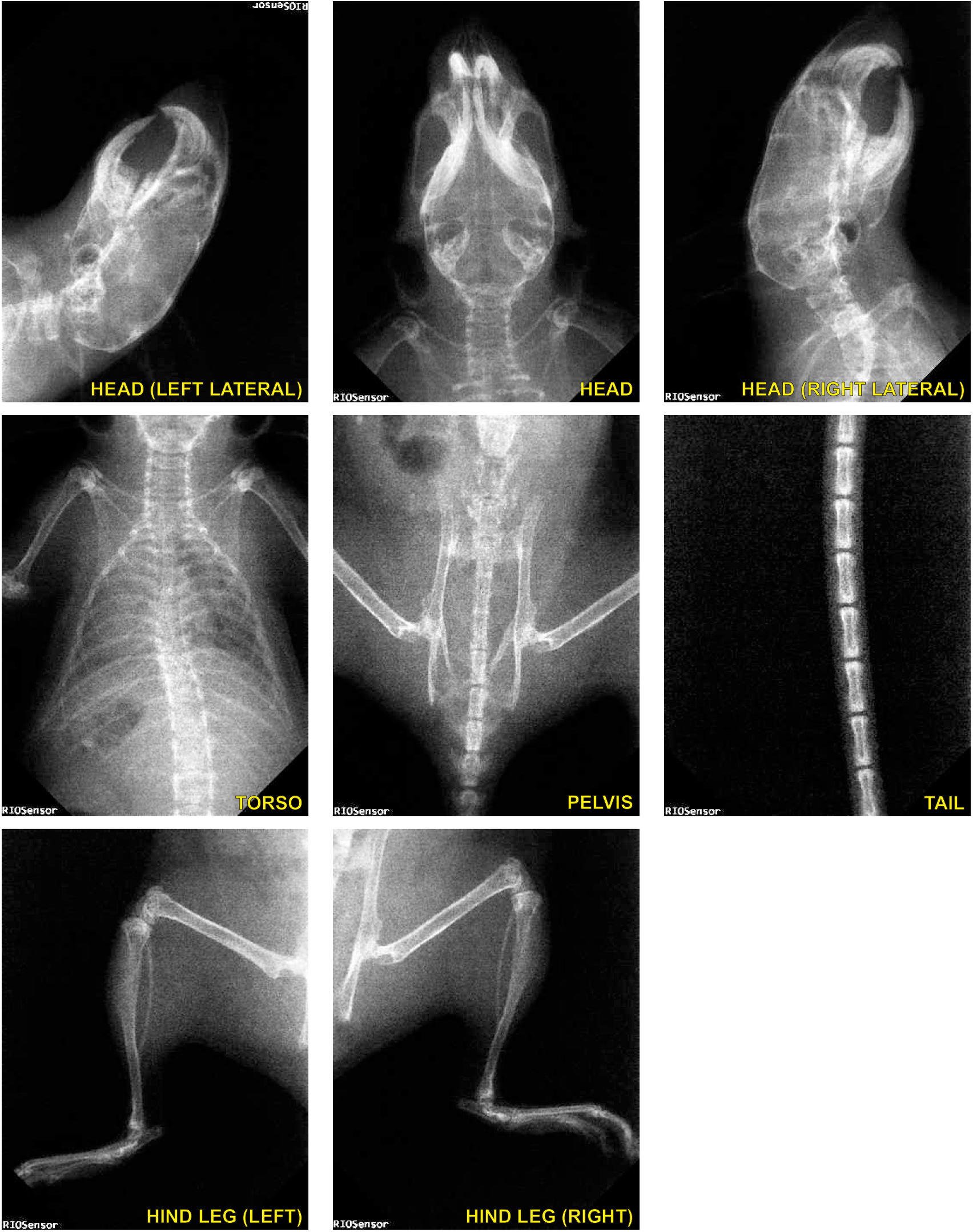
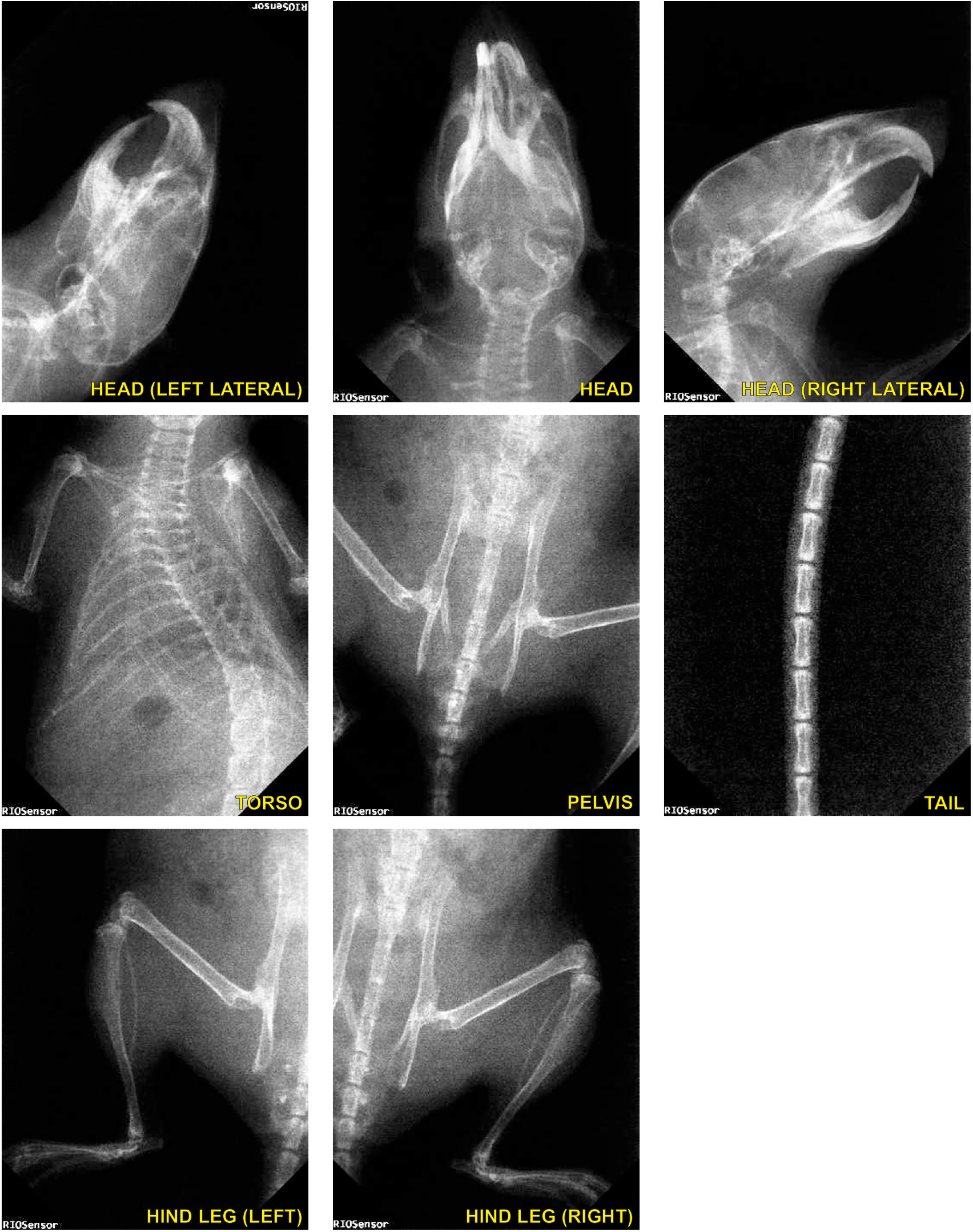
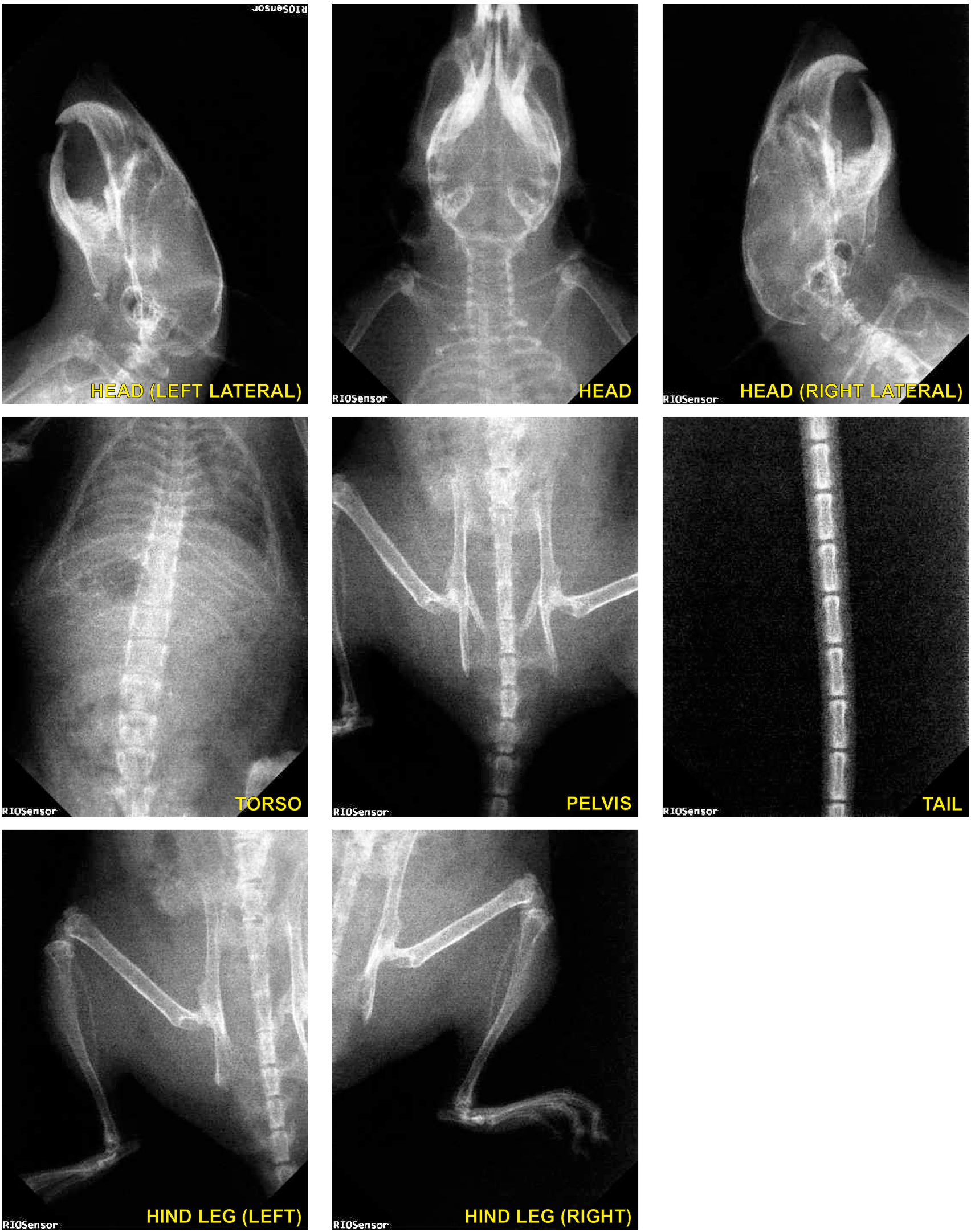
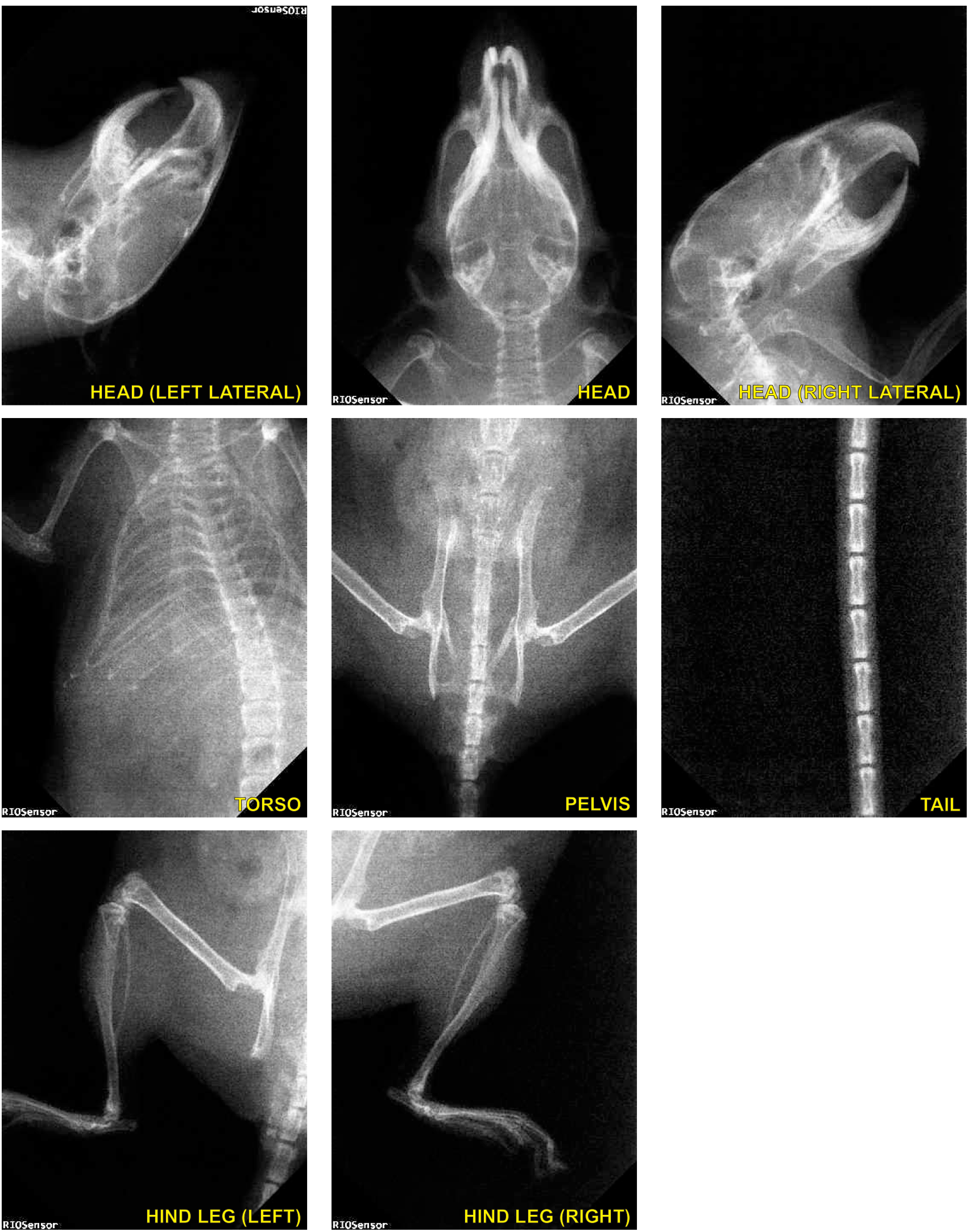
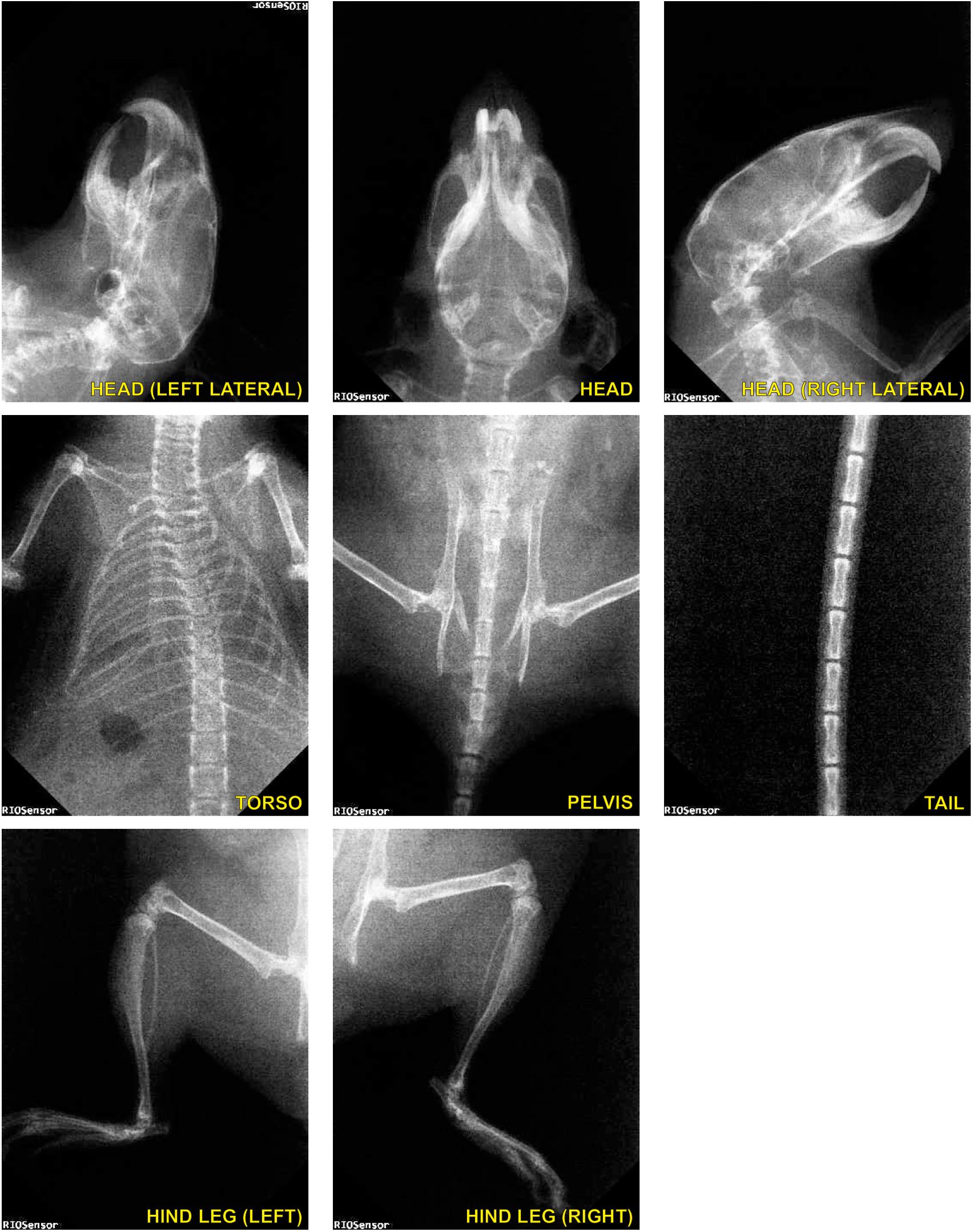
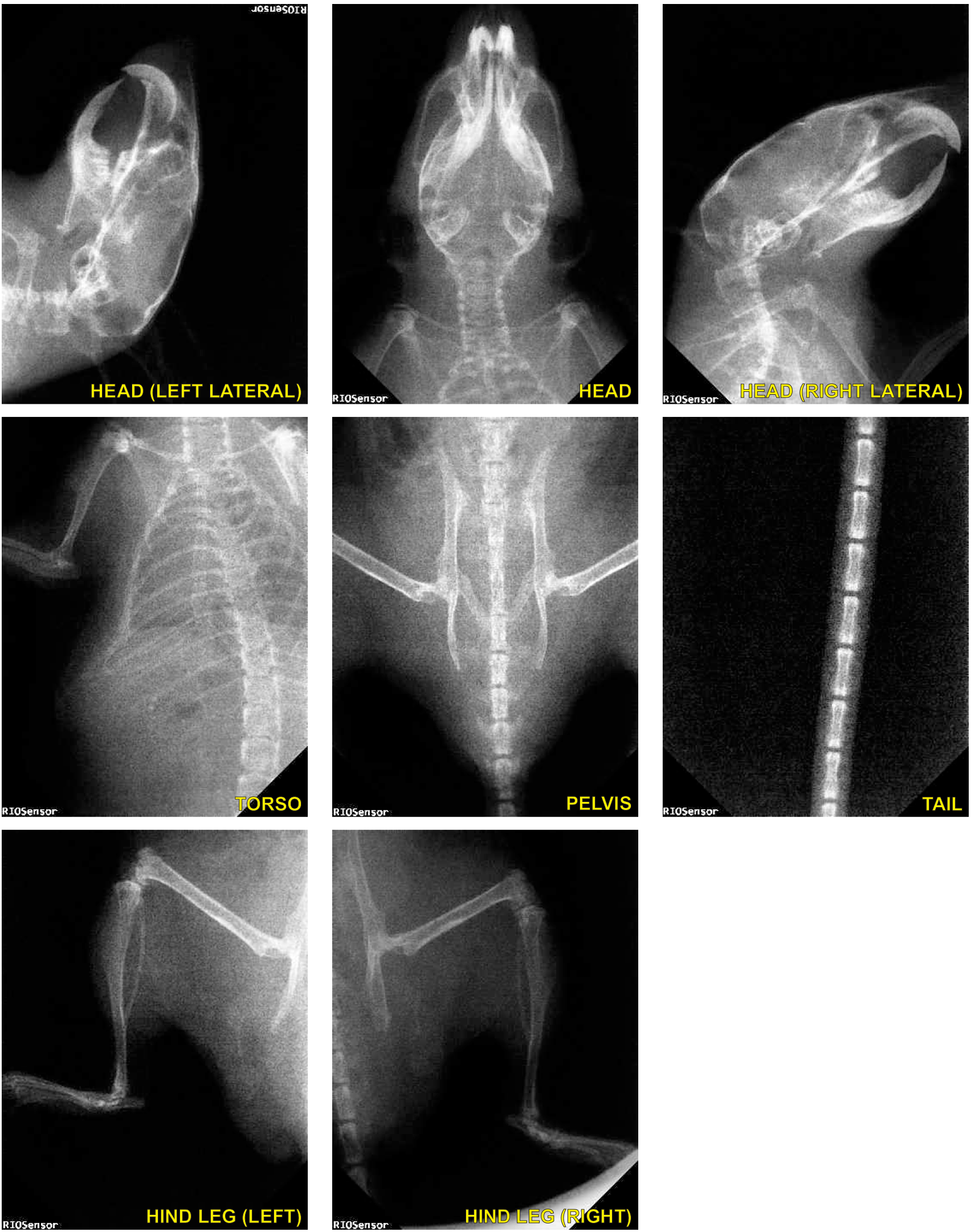
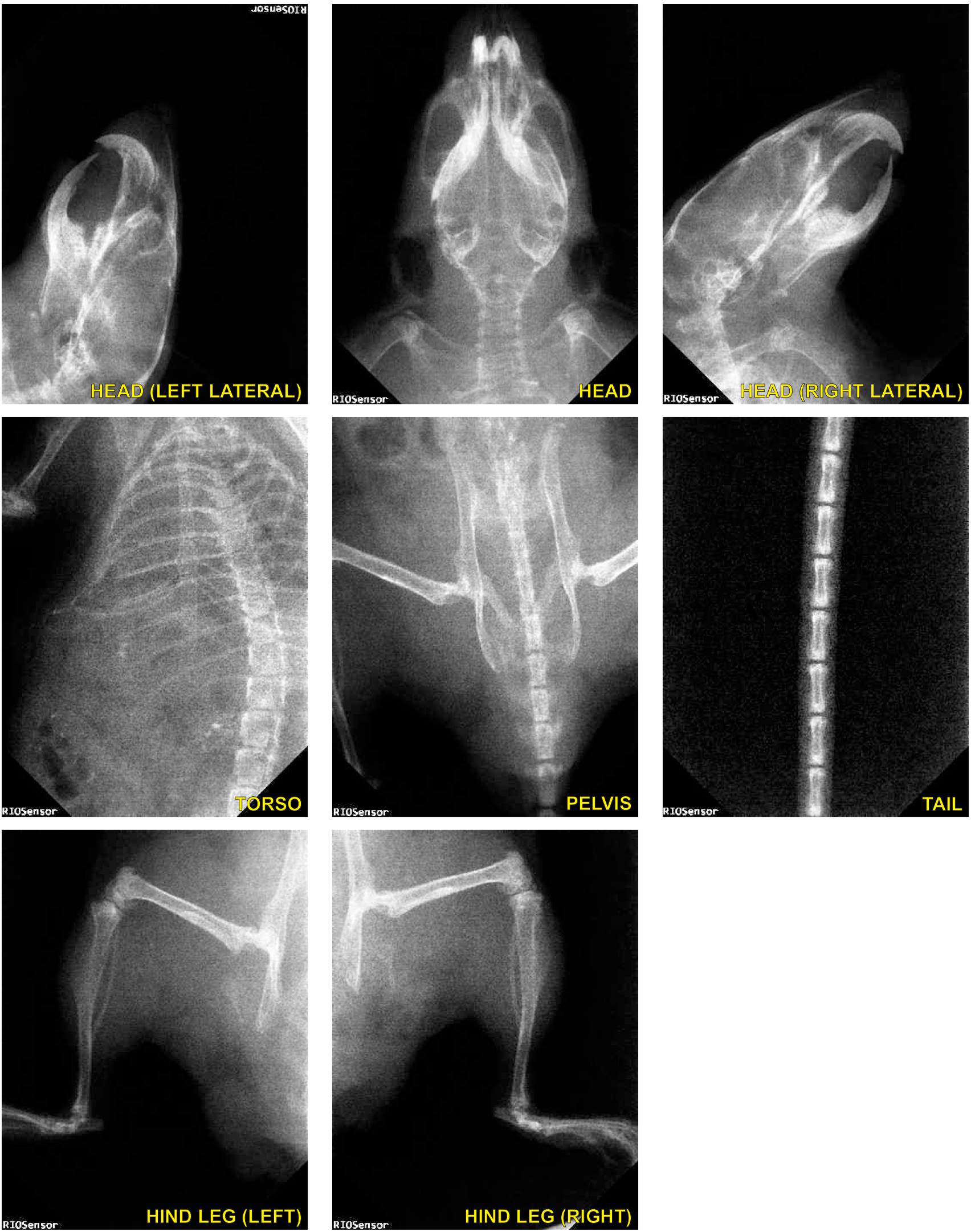

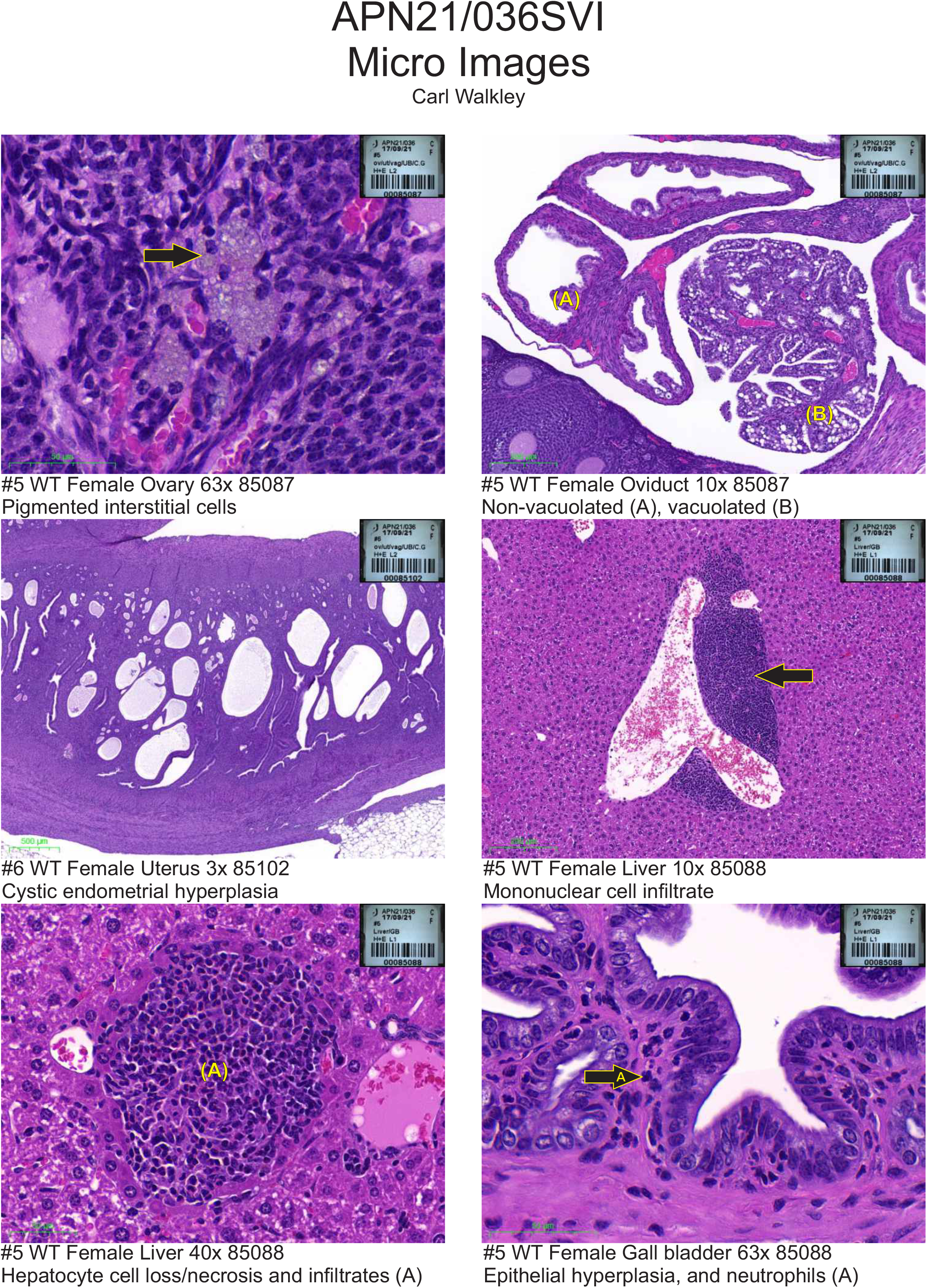
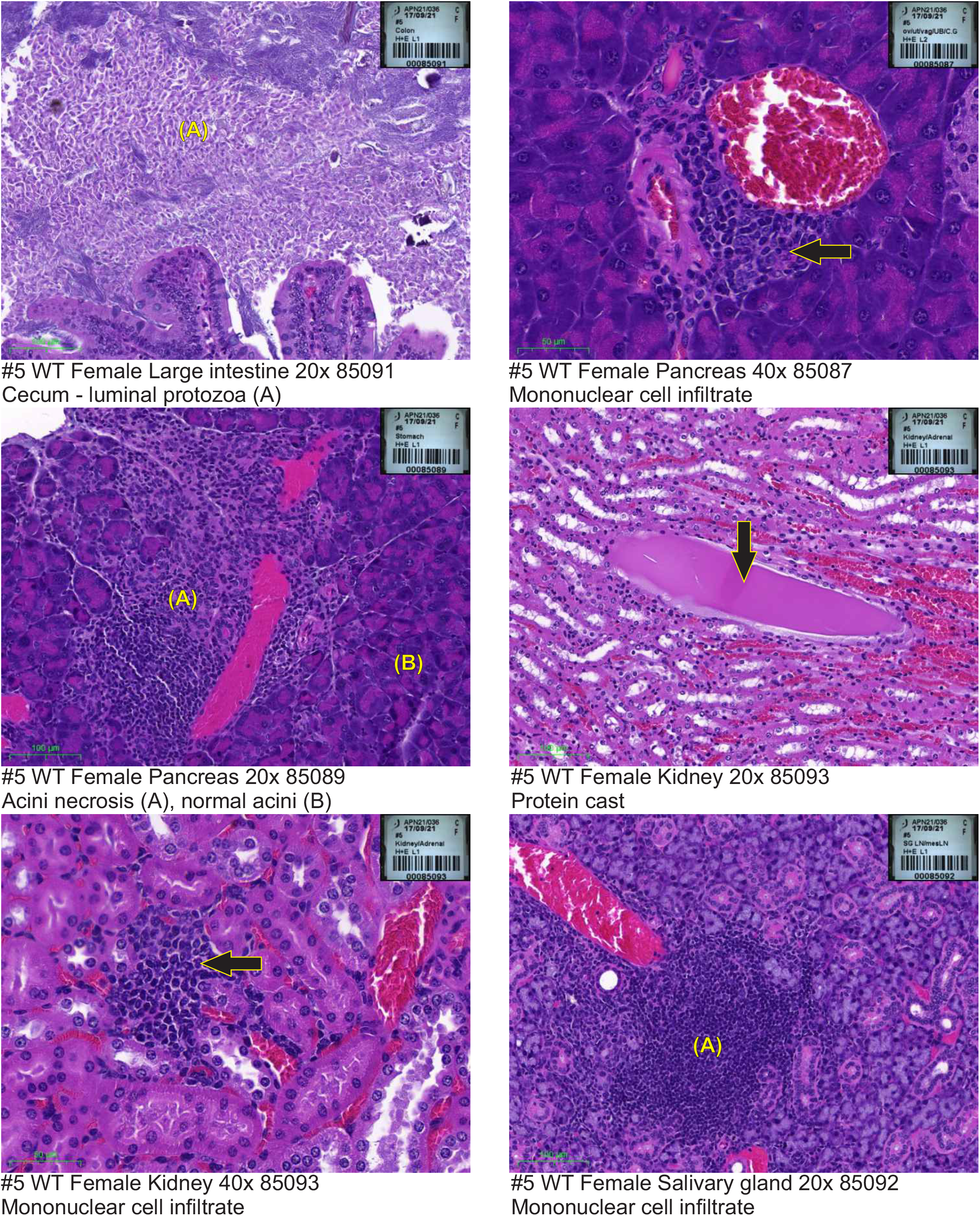
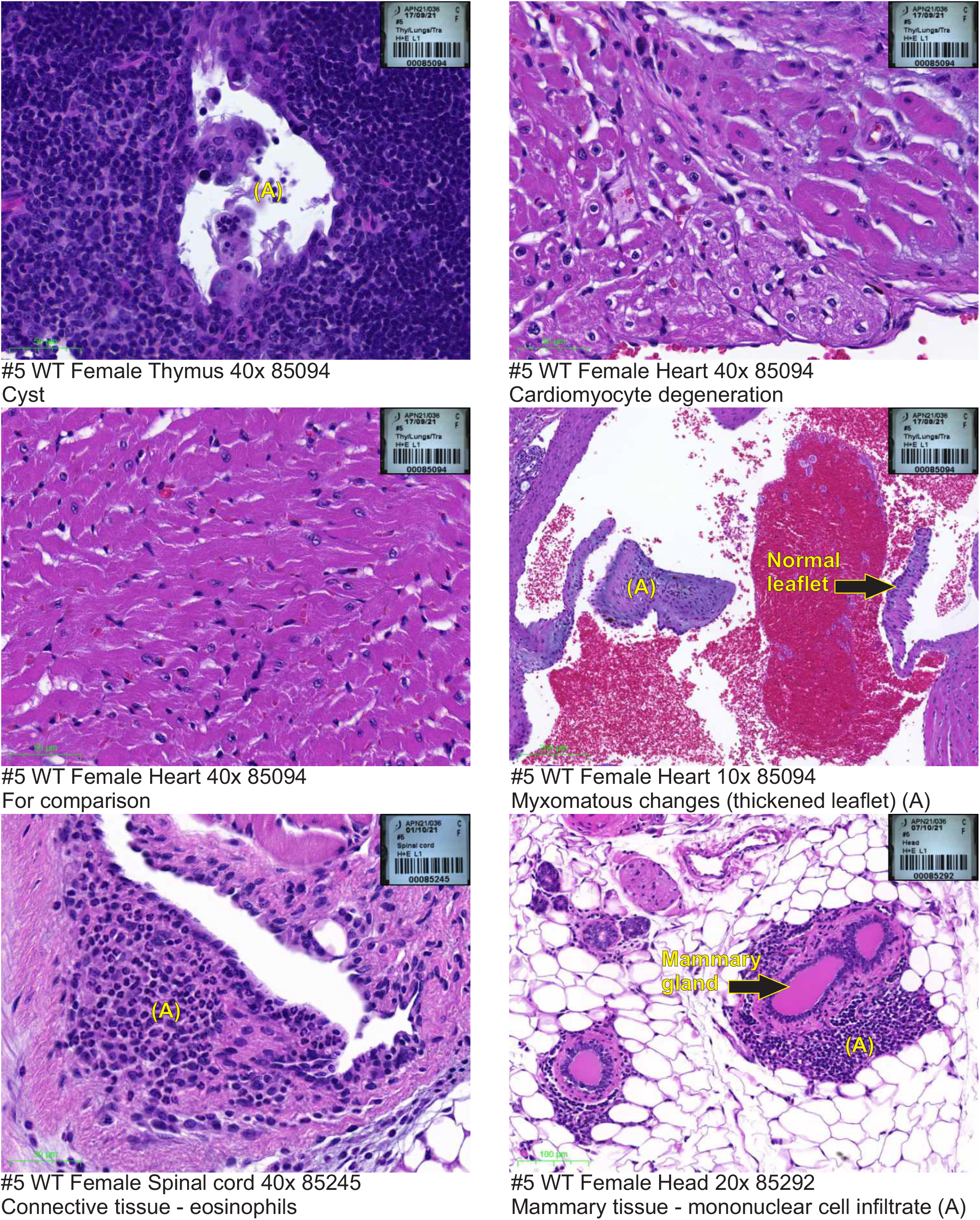
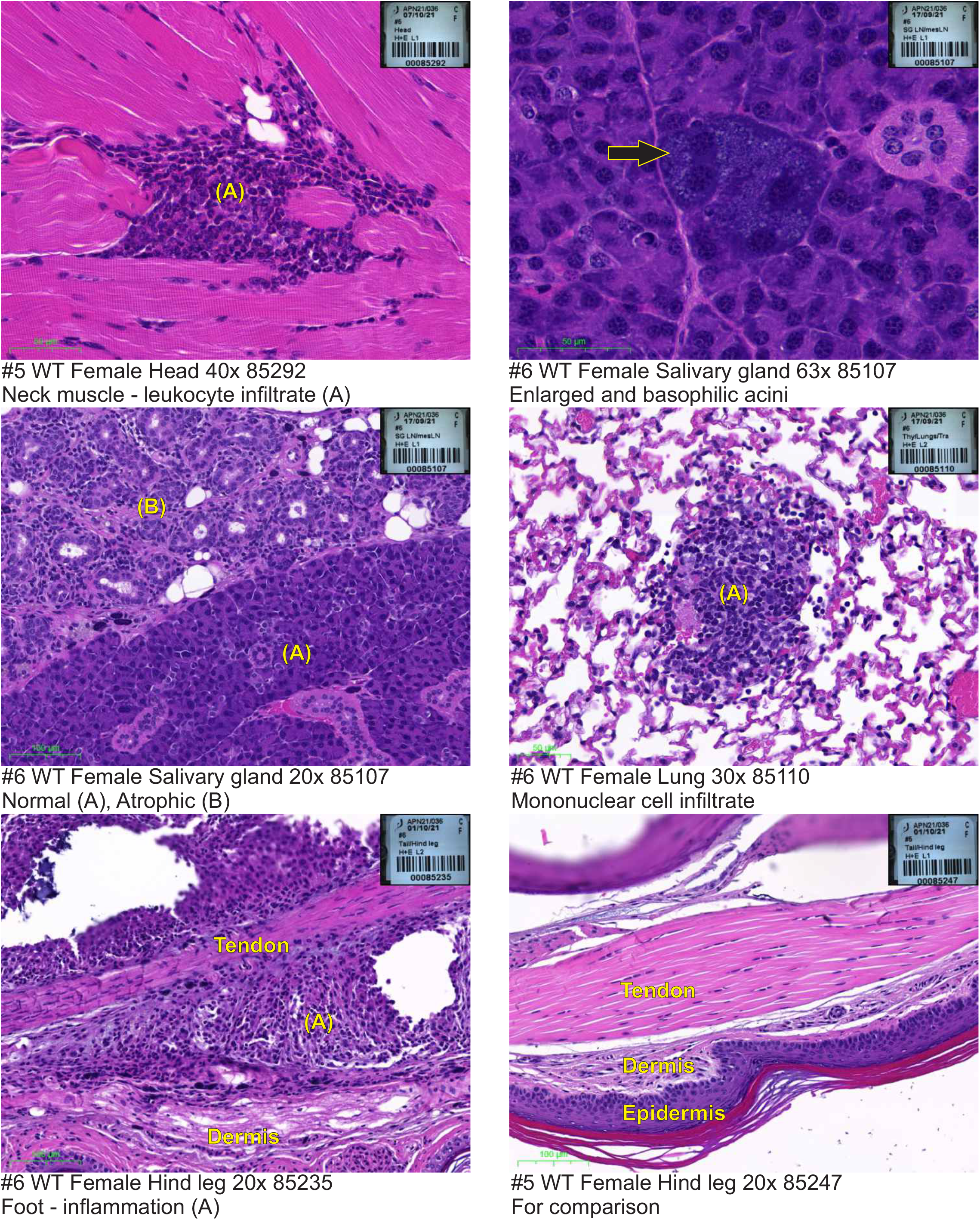
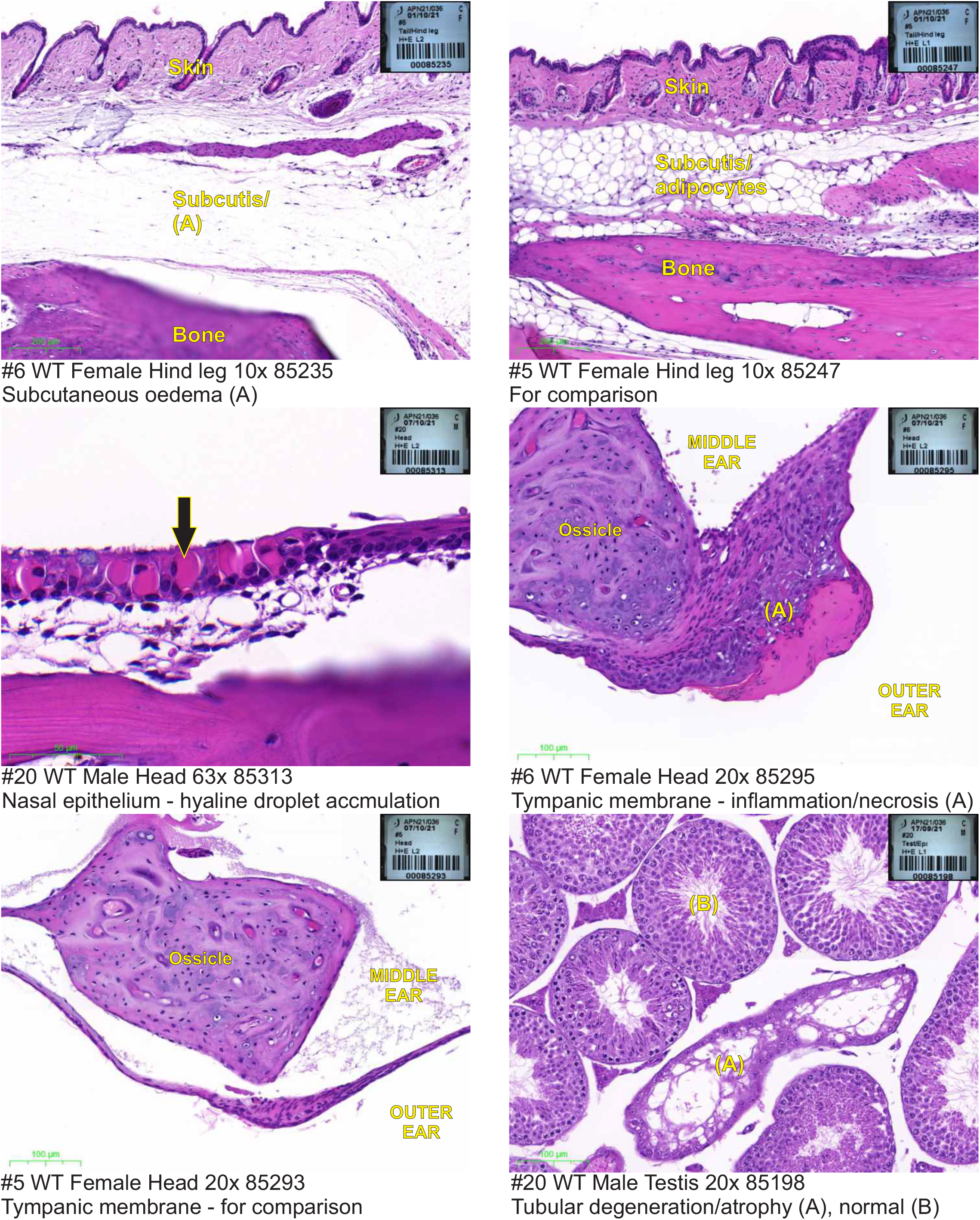
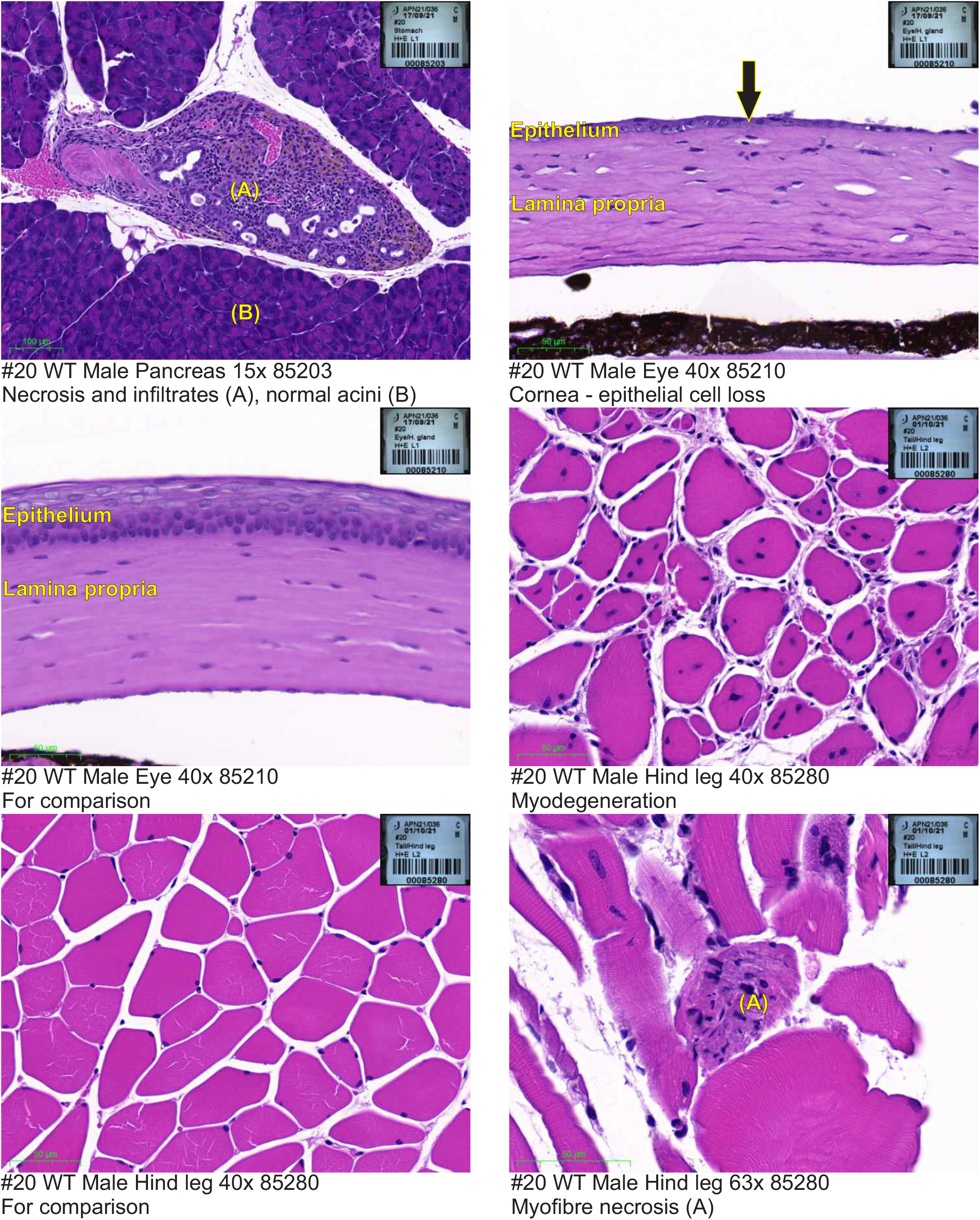
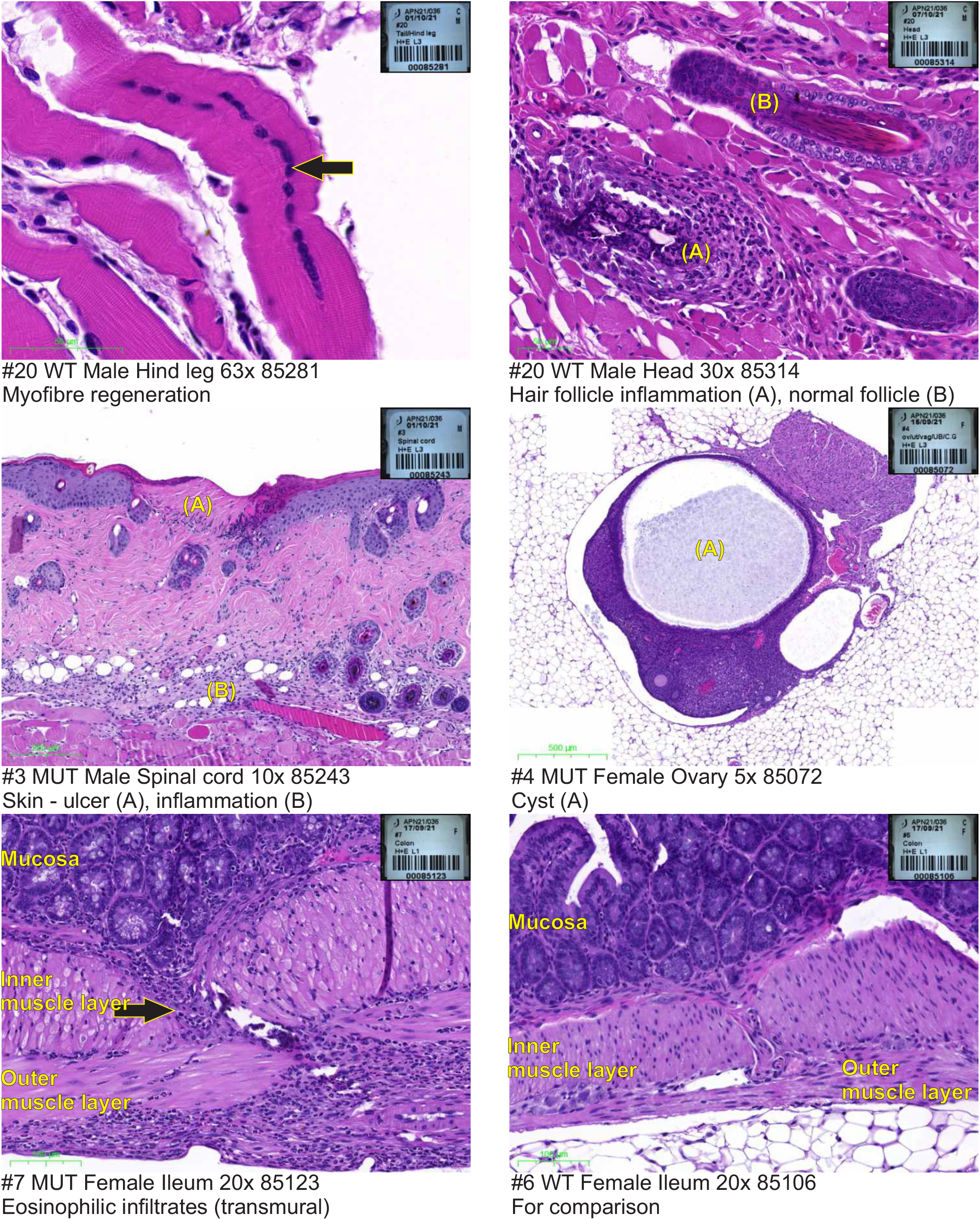
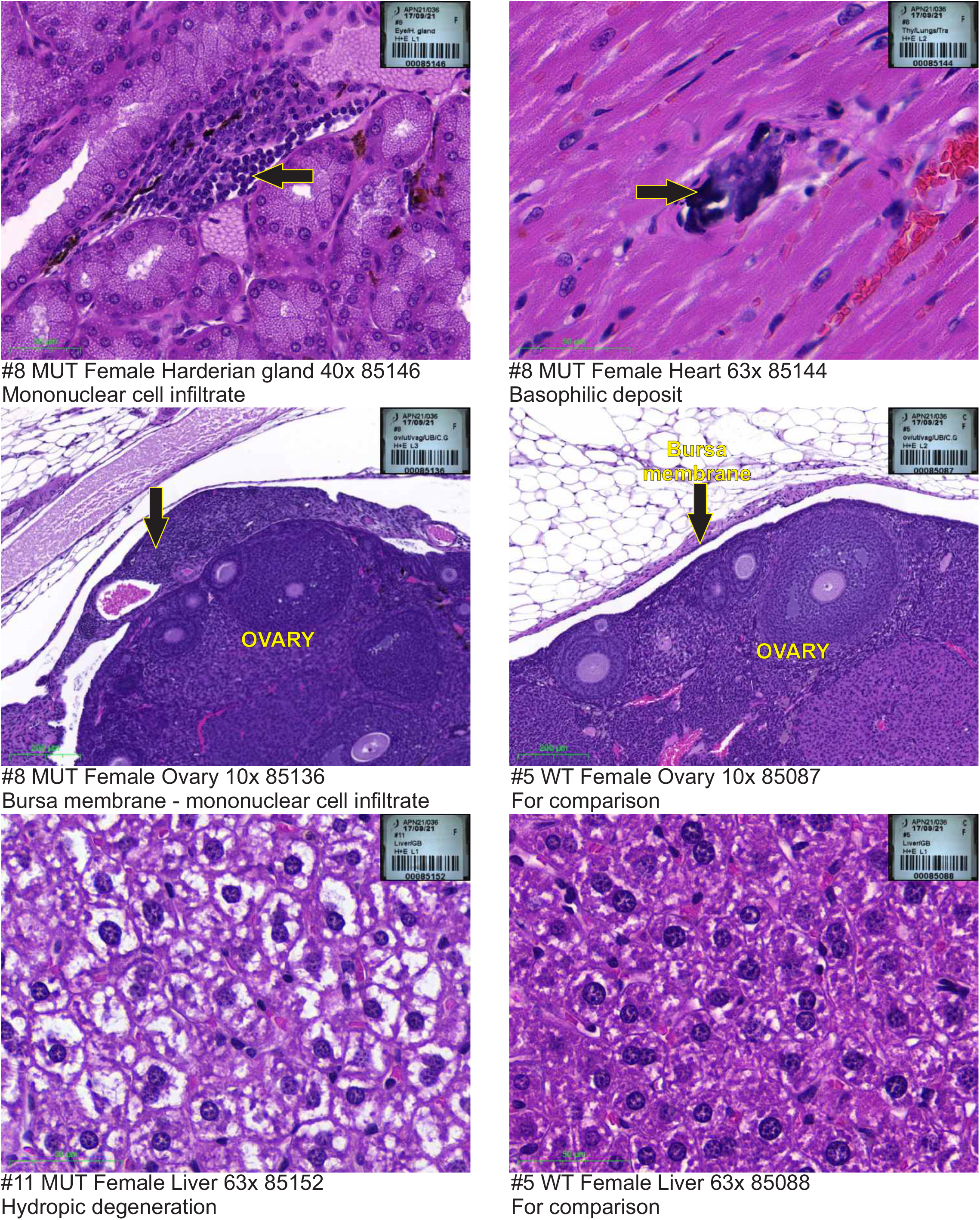
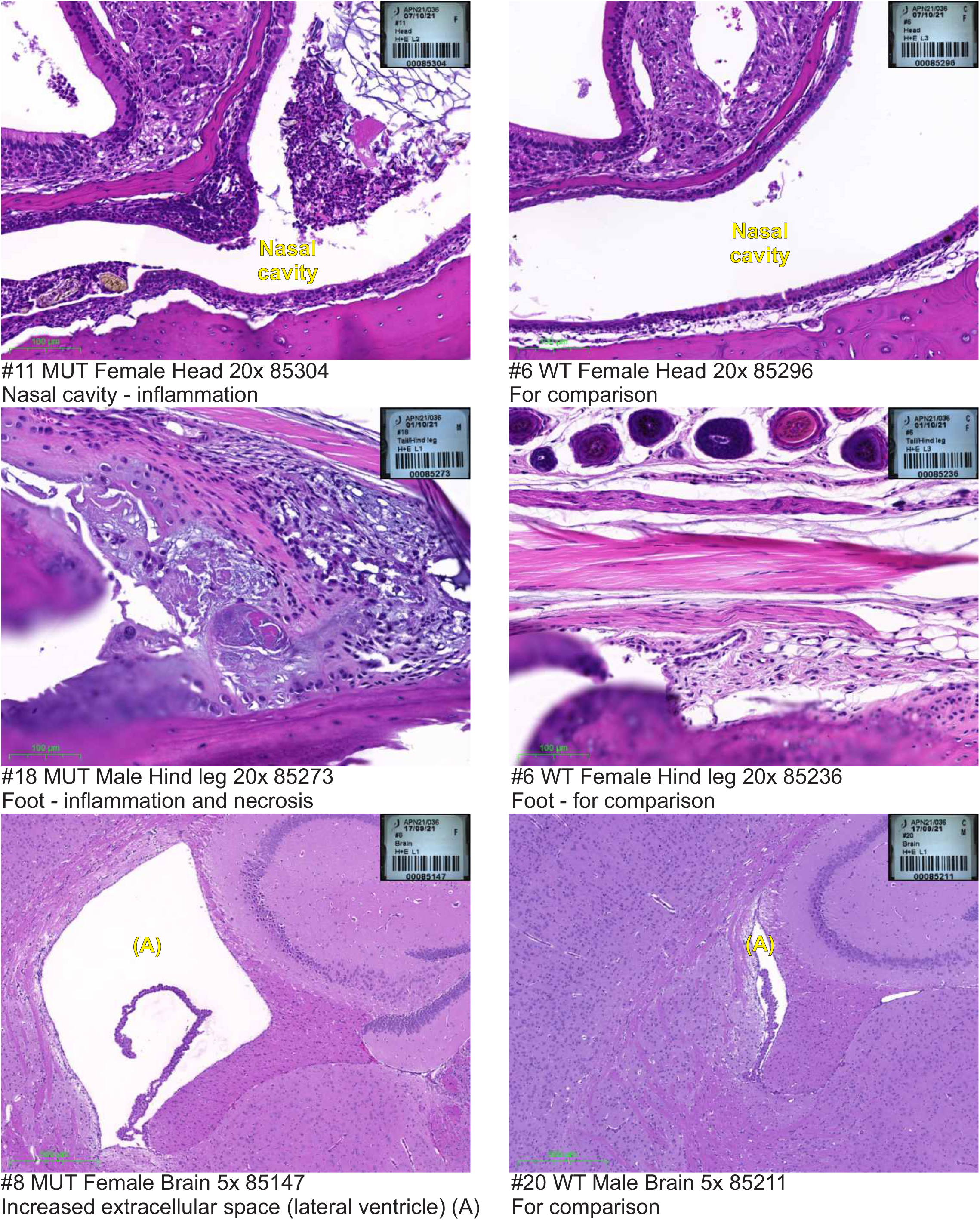

## 9.1 Histopathology Report

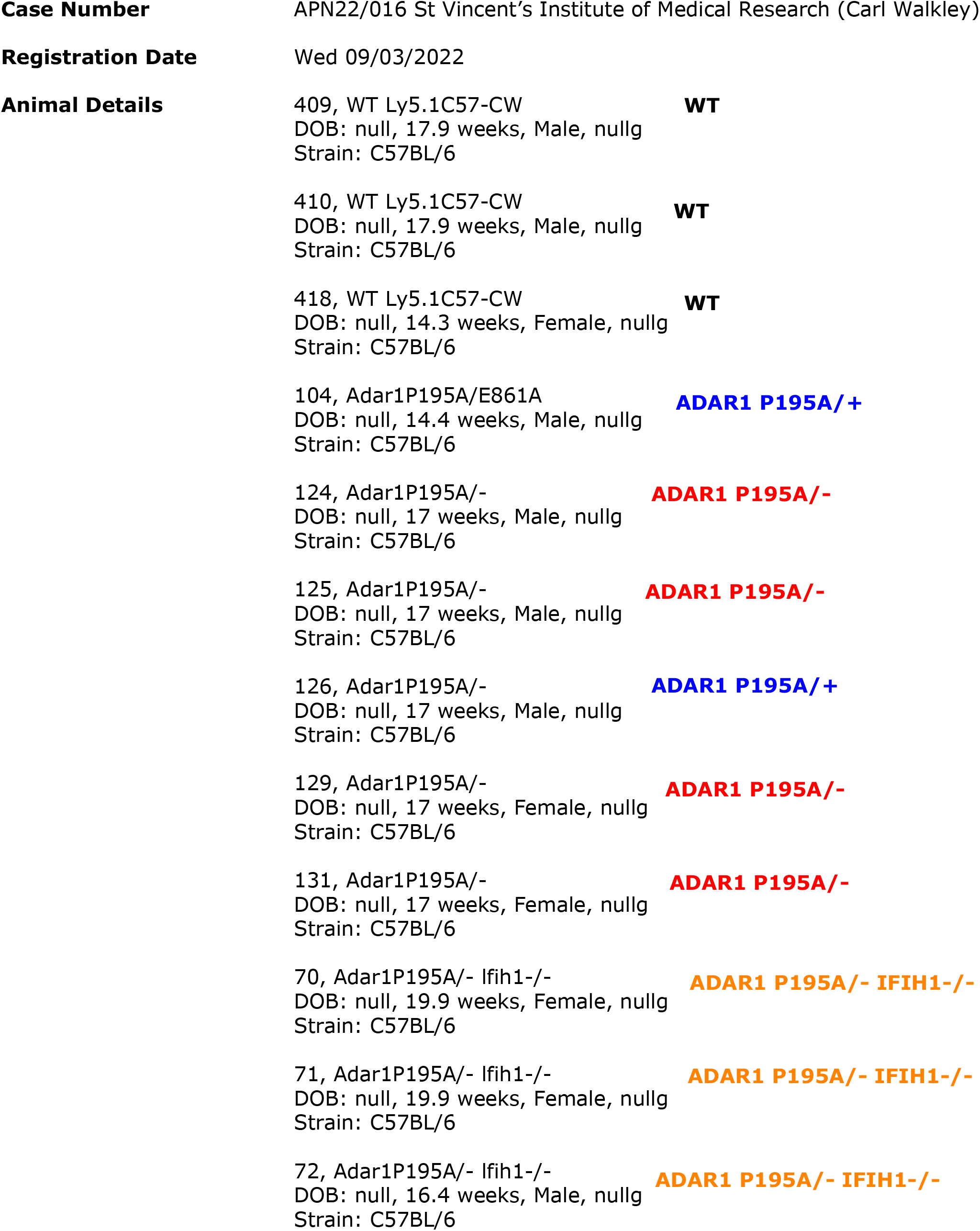
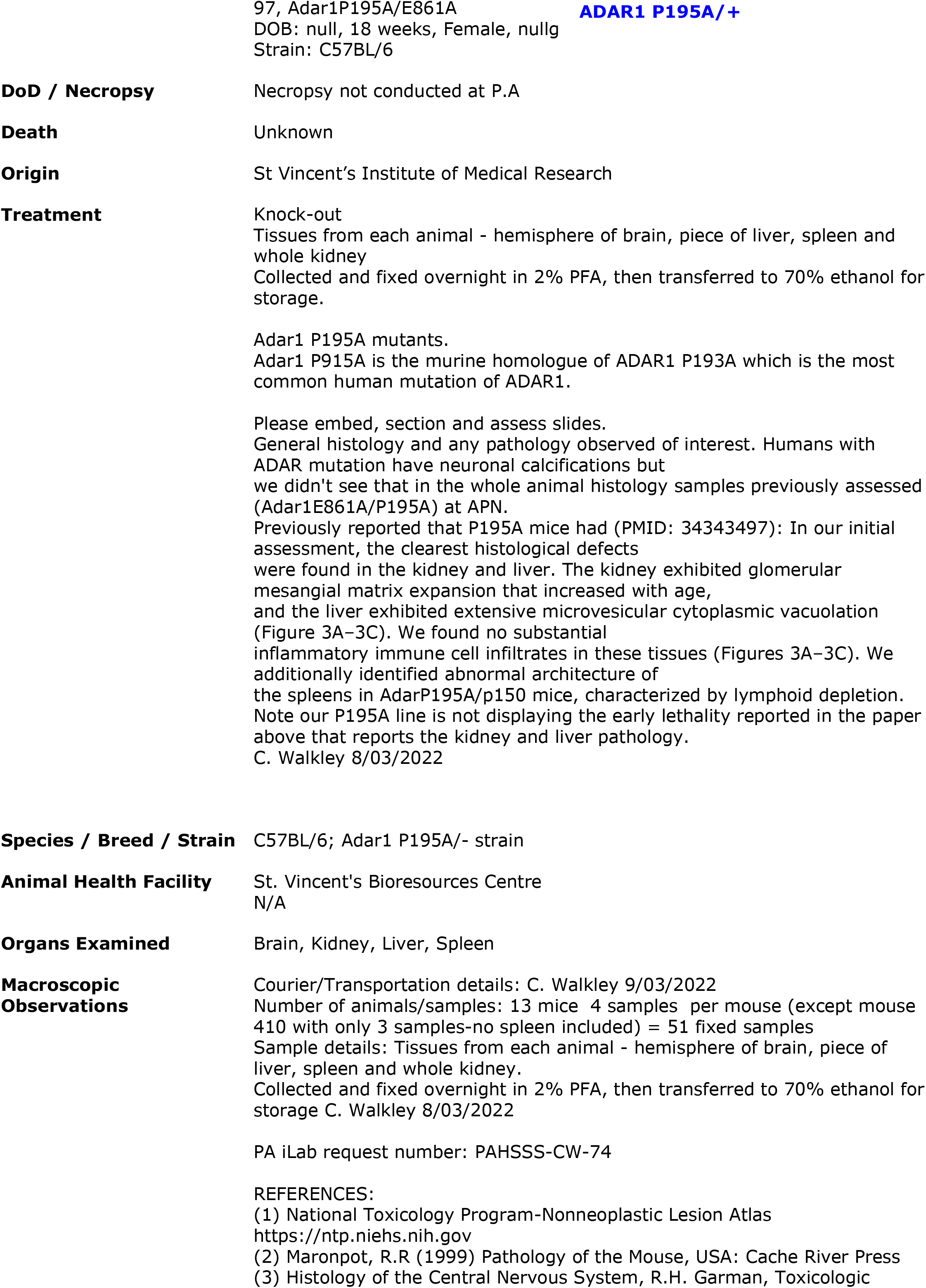
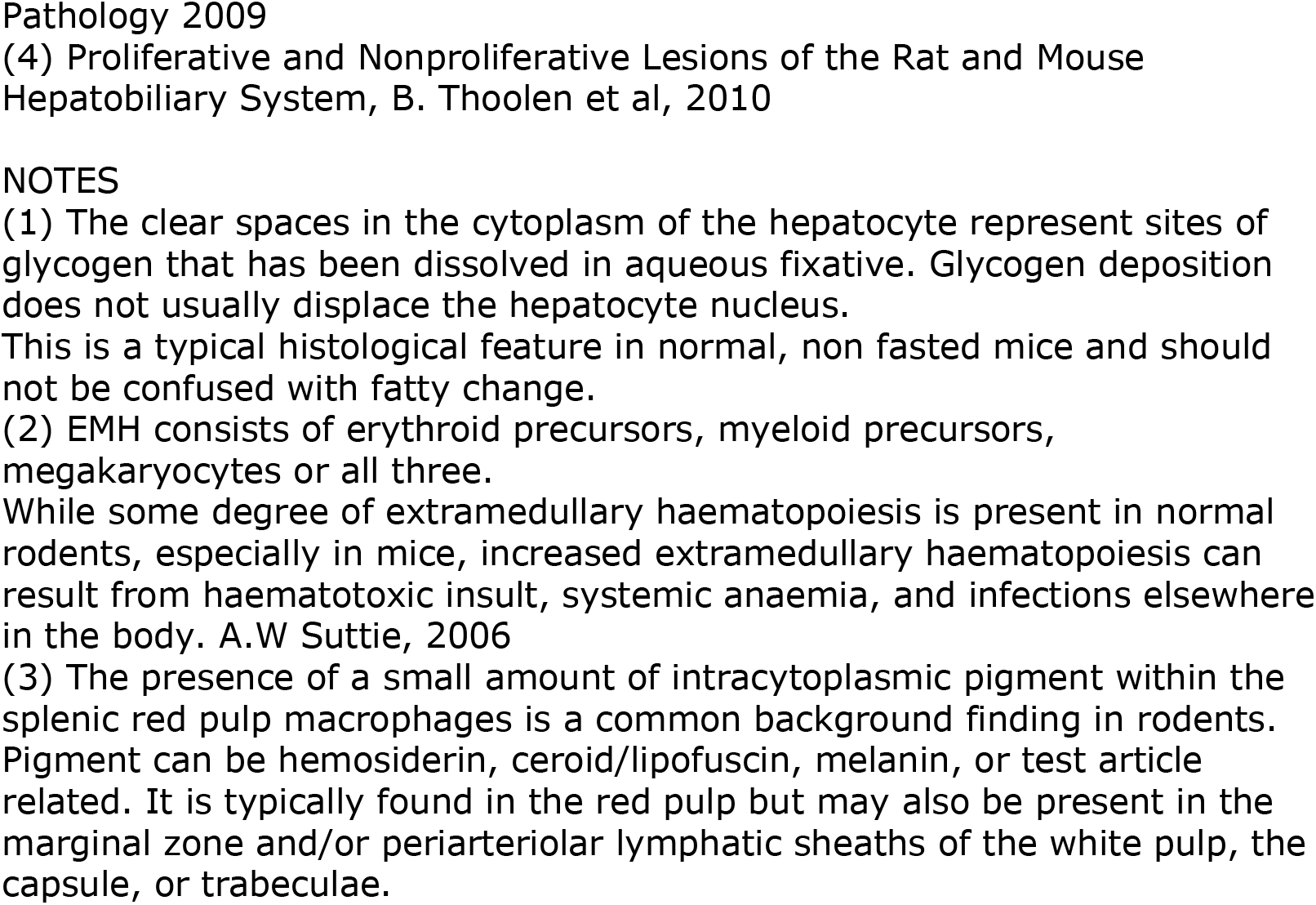

### Microscopic Observations

#### SUMMARY

70 Brain (87505)-cellularity/lamination of cerebral cortex or plane of section. Kidney (87508)- Tubule protein cast in medulla.

71 kidney (87504)-interstitial inflammatory cell infiltrates

72 Kidney (87500)- inflammatory cell infiltrate (lymphocytic)

97 NLS Liver(87495)-Mild multifocal inflammatory cell aggregate. Brain (87493)-NLS

104 NLS Brain (87489)-NLS

124 NLS Brain(87481)-NLS

125 NLS Brain(87477)-NLS

126 NLS Spleen (87463)-query less white pulp. Likely to be plane of section.

Brain (87461, 87465)-NLS

129 Brain (87473)-cellularity/lamination of cerebral cortex, likely to be plane of section.

Liver (87474)- inflammatory cell infiltrate. Spleen(87475)-moderate EMH

131 Liver (87487)-periportal inflammatory cell aggregates with hepatocyte loss.

Brain -NLS (87485)

409 NLS Brain (87469)-NLS

410 NLS Brain (87466)-NLS

418 NLS Brain (87457)-NLS

NLS= No lesions of significance

#### 409 (control)

Macro Observations

Fixed spleen, liver, brain & kidney
Kidney 11×6×4mm
No macroscopic lesions identified.

Micro Observations

WT Ly5.1C57-CW

Liver

Section shows typical liver parenchyma including hepatocytes, Kupffer cells, portal triads and central veins. A small aggregation of Kupffer cells with trivial hepatocyte loss.
No lesions of significance
(87470)

Spleen

Unremarkable follicular pattern identified with typical red and white pulp micromorphology.
No lesions of significance
(87471)

Kidney

H&E section show cortex, medulla and papilla regions of the kidney.
Typical glomeruli, many with open capillary lumens. The expected cuboidal epithelium lining the
Bowman’s capsule is apparent in many glomeruli.
The interstitium and tubules is unremarkable.
No lesions of significance
(87472)

Brain

Slide shows sagittal sections of one hemisphere demonstrating, meninges, olfactory lobe, cerebral cortex with typical lamination, corpus callosum, ventricles and choroid plexi, hippocampus, thalamus, hypothalamus and hind brain (cerebellum, pons). Unremarkable cerebellum with typical architecture and Purkinje cells. No obvious neuronal loss, and the myelination appears normal. Readily identified dark neuron fixative artifact, a common feature.
No lesions of significance
(87469)

#### 410 (control)

Macro Observations

Fixed liver, brain & kidney
Kidney 11×6×4mm
No macroscopic lesions identified.

Micro Observations

WT Ly5.1C57-CW

Liver

Section shows typical liver parenchyma including hepatocytes, Kupffer cells, portal triads and central veins. Unremarkable Gall bladder
No lesions of significance
(87467)

Kidney

H&E section show cortex, medulla and papilla regions (and ureter) of the kidney.
Typical glomeruli, many with open capillary lumens. The expected cuboidal epithelium lining the Bowman’s capsule is apparent in many glomeruli.
The interstitium and tubules is unremarkable.
No lesions of significance (87468)

Brain

Slide shows sagittal sections of one hemisphere demonstrating, meninges, small portion of the olfactory lobe, cerebral cortex with typical lamination, corpus callosum, ventricles and choroid plexi, hippocampus, thalamus, hypothalamus and hind brain (cerebellum). Unremarkable cerebellum with typical architecture and Purkinje cells. No obvious neuronal loss, and the myelination appears normal. Readily identified dark neuron fixative artifact, a common feature.
No lesions of significance
(87466)

#### 418 (control)

Macro Observations

Fixed spleen, liver, brain & kidney
Kidney 9×5×3mm
No macroscopic lesions identified.

Micro Observations

WT Ly5.1C57-CW

Liver

Section shows typical liver parenchyma including hepatocytes, Kupffer cells, portal triads and central veins. A small aggregation of Kupffer cells with trivial hepatocyte loss.
No lesions of significance
(87460)

Spleen

Unremarkable follicular pattern identified with typical red and white pulp micromorphology.
No lesions of significance
(87458)

Kidney

H&E section show cortex, medulla and papilla regions of the kidney.
Typical glomeruli, many with open capillary lumens. The expected cuboidal epithelium lining the Bowman’s capsule is apparent in many glomeruli.
The interstitium and tubules is unremarkable.
No lesions of significance (87459)

Brain

Slide shows sagittal sections of one hemisphere demonstrating, meninges, olfactory lobe, cerebral cortex with typical lamination, corpus callosum, ventricles and choroid plexi, hippocampus, thalamus, hypothalamus and hind brain (cerebellum, pons). Unremarkable cerebellum with typical architecture and Purkinje cells. No obvious neuronal loss, and the myelination appears normal. Readily identified dark neuron fixative artifact, a common feature. Section includes small portion of exocrine tissue (pancreas) judged to be cross contamination
No lesions of significance
(87457)

#### 104

Macro Observations

Fixed spleen, liver, brain & kidney
Kidney 12×7×4mm
No macroscopic lesions identified.

Micro Observations

Adar1P195A/E861A

Liver

Section shows typical liver parenchyma including hepatocytes, Kupffer cells, portal triads and central veins.
No lesions of significance
(87491)

Spleen

Unremarkable follicular pattern identified with typical red and white pulp micromorphology.
Mild extramedullary haematopoiesis identified in the red pulp, a common feature in the mouse spleen.
No lesions of significance (87490)

Kidney

H&E section show cortex, medulla and papilla regions of the kidney.
Typical glomeruli, many with open capillary lumens. The expected cuboidal epithelium lining the Bowman’s capsule is apparent in many glomeruli.
The interstitium and tubules is unremarkable.
No lesions of significance (87492)

Brain

Slide shows sagittal sections of one hemisphere demonstrating, meninges, cerebral cortex with typical cellular lamination, corpus callosum, ventricles and representative choroid plexus, hippocampus, subfornical organ, thalamus, hypothalamus and cerebellum. Unremarkable cerebellum with typical architecture and Purkinje cells. No obvious neuronal loss, and the myelination appears normal.
No lesions of significance
(87489)

#### 124

Macro Observations

Fixed spleen, liver, brain & kidney
Kidney 13×7×5mm
No macroscopic lesions identified.

Micro Observations

Adar1P195A/−

Liver

Section shows typical liver parenchyma including hepatocytes, Kupffer cells, portal triads and central veins.
No lesions of significance
(87482)

Spleen

Unremarkable follicular pattern identified with typical red and white pulp micromorphology.
Mild extramedullary haematopoiesis identified in the red pulp as well as brown pigment, likely to be haemosiderin.
No lesions of significance
(87483)

Kidney

H&E section show cortex, medulla and papilla regions of the kidney.
Typical glomeruli, many with open capillary lumens. The expected cuboidal epithelium lining the Bowman’s capsule is apparent in many glomeruli.
The interstitium and tubules is unremarkable.
No lesions of significance
(87484)

Brain

Slide shows sagittal sections of one hemisphere demonstrating, meninges, small portion of the olfactory lobe, cerebral cortex with typical cellular lamination, corpus callosum, ventricles and choroid plexi, hippocampus, thalamus, hypothalamus and hind brain (cerebellum). Unremarkable cerebellum with typical architecture and Purkinje cells. No obvious neuronal loss, and the myelination appears normal. Readily identified dark neuron fixative artifact, a common feature.
No lesions of significance
(87481)

#### 125

Macro Observations

Fixed spleen, liver, brain & kidney
Kidney 13×8×4mm
No macroscopic lesions identified.

Micro Observations

Adar1P195A/−

Liver

Section shows typical liver parenchyma including hepatocytes, Kupffer cells, portal triads and central veins.
No lesions of significance
(87479)

Spleen

Unremarkable follicular pattern identified with typical red and white pulp micromorphology. Moderate extramedullary haematopoiesis identified in the red pulp as well as brown pigment, likely to be haemosiderin. No lesions of significance (87478)

Kidney

H&E section show cortex, medulla and papilla regions of the kidney.
Typical glomeruli, many with open capillary lumens. The expected cuboidal epithelium lining the Bowman’s capsule is apparent in many glomeruli.
The interstitium and tubules is unremarkable.
No lesions of significance (87480)

Brain

Slide shows sagittal sections of one hemisphere demonstrating, meninges, small portion of the olfactory lobe, cerebral cortex with typical cellular lamination, corpus callosum, ventricles and choroid plexi, hippocampus, thalamus, hypothalamus and hind brain (cerebellum, pons).
Unremarkable cerebellum with typical architecture and Purkinje cells. No obvious neuronal loss, and the myelination appears normal. Readily identified dark neuron fixative artifact, a common feature.
No lesions of significance
(87477)

#### 126

Macro Observations

Fixed spleen, liver, brain & kidney
Kidney 13×7×5mm -mechanical artefact
No macroscopic lesions identified.
Micro Observations Adar1P195A/−

Liver

Section shows typical liver parenchyma including hepatocytes, Kupffer cells, portal triads and central veins.
No lesions of significance
(87462)

Spleen

Discernible red and white pulp micromorphology.
Less white pulp, judged to be plane of section.
Mild extramedullary haematopoiesis identified in the red pulp. (87463)
*Comments:*
*Pathology to comment*

Kidney

H&E section show cortex, medulla and papilla regions of the kidney.
Typical glomeruli, many with open capillary lumens. The expected cuboidal epithelium lining the Bowman’s capsule is apparent in many glomeruli.
The interstitium and tubules is unremarkable.
No lesions of significance (87464)

Brain

Section shows several pieces of non-intact brain. Limited evaluation due to mechanical artefact. Discernible and unremarkable cerebellum and olfactory bulb region.
No observable lesions. Cellularity and lamination appears unremarkable. (87461, 87465)
*Comments:*
*Neuropathology to comment*

#### 129

Macro Observations

Fixed spleen, liver, brain & kidney
Kidney 11×6×4mm
No macroscopic lesions identified.

Micro Observations

Adar1P195A/−

Liver

Section shows liver parenchyma including hepatocytes, Kupffer cells, portal triads and central veins.
Mild, multifocal inflammatory cell aggregate with some hepatocyte loss. (87474)
*Comments:*
*Pathology to comment*

Spleen

Unremarkable follicular pattern identified with typical red and white pulp micromorphology. Moderate extramedullary haematopoiesis identified in the red pulp as well as brown pigment, likely to be haemosiderin.
(87475)

Kidney

H&E section show cortex, medulla and papilla regions of the kidney.
Typical glomeruli, many with open capillary lumens. The expected cuboidal epithelium lining the Bowman’s capsule is apparent in many glomeruli.
The interstitium and tubules is unremarkable.
No lesions of significance
(87476)

Brain

Slide shows sagittal sections of one hemisphere demonstrating, meninges, olfactory lobe, cerebral cortex, corpus callosum, ventricles and choroid plexi, thalamus, hypothalamus and hind brain (cerebellum, pons). Unremarkable cerebellum with typical architecture and Purkinje cells.
Irregular cellular lamination of the cerebral cortex, judged to be plane of section. (87473)
*Comments:*
*Neuropathology to comment*

#### 131

Macro Observations

Fixed spleen, liver, brain & kidney
Kidney 12×6×4mm
No macroscopic lesions identified.

Micro Observations

Adar1P195A/−

Liver

Section shows typical liver parenchyma including hepatocytes, Kupffer cells, portal triads and central veins.
Mild, multifocal centrilobular and periportal inflammatory cell aggregate with some hepatocyte loss.
(87487)
*Comments:*
*Pathology to comment*

Spleen

Unremarkable follicular pattern identified with typical red and white pulp micromorphology. Mild extramedullary haematopoiesis identified in the red pulp, a common feature in the mouse spleen.
No lesions of significance
(87486)

Kidney

H&E section show cortex, medulla and papilla regions of the kidney.
Typical glomeruli, many with open capillary lumens. The expected cuboidal epithelium lining the Bowman’s capsule is apparent in many glomeruli.
The interstitium and tubules is unremarkable.
No lesions of significance
(87488)

Brain

Slide shows sagittal sections of one hemisphere demonstrating, meninges, small portion of the olfactory lobe, cerebral cortex with typical cellular lamination, corpus callosum, ventricles and representative choroid plexus, hippocampus, subfornical organ, thalamus, hypothalamus, cerebellum and pons. Unremarkable cerebellum with typical architecture and Purkinje cells. No obvious neuronal loss, and the myelination appears normal. Readily identified dark neuron fixative artifact, a common feature.
No lesions of significance
(87485)

#### 70

Macro Observations

Fixed spleen, liver, brain & kidney
Kidney 10×5×3mm
No macroscopic lesions identified.

Micro Observations

Adar1P195A/− lfih1−/−

Liver

Section shows typical liver parenchyma including hepatocytes, Kupffer cells, portal triads and central veins.
Unremarkable Gall bladder. No lesions of significance
(87507)

Spleen

Unremarkable follicular pattern identified with typical red and white pulp micromorphology. Mild extramedullary haematopoiesis identified in the red pulp, a common feature in the mouse spleen. No lesions of significance
(87506)

Kidney

H&E section show cortex, medulla and papilla regions of the kidney.
Typical glomeruli, many with open capillary lumens. The expected cuboidal epithelium lining the Bowman’s capsule is apparent in many glomeruli.
Tubule protein cast in medulla.
The remaining interstitium and tubules is unremarkable.
(87508)
*Comments:*
*Pathology to comment*

Brain

Slide shows sagittal sections of one hemisphere demonstrating, meninges, cerebral cortex, corpus callosum, ventricles and representative choroid plexus, subfornical organ, hippocampus, thalamus, hypothalamus and cerebellum. Query cellularity/lamination of cerebral cortex or plane of section.
Unremarkable cerebellum with typical architecture and Purkinje cells. No obvious neuronal loss, and the myelination appears normal.
(87505)
*Comments:*
*Neuropathology to comment*

#### 71

Macro Observations

Fixed spleen, liver, brain & kidney
Kidney 11×6×4mm
No macroscopic lesions identified.

Micro Observations

Adar1P195A/− lfih1−/−

Liver

Section shows typical liver parenchyma including hepatocytes, Kupffer cells, portal triads and central veins.
No lesions of significance
(87503)

Spleen

Unremarkable follicular pattern identified with typical red and white pulp micromorphology.
Mild extramedullary haematopoiesis identified in the red pulp, a common feature in the mouse spleen.
No lesions of significance (87502)

Kidney

H&E section show cortex, medulla and papilla regions of the kidney.
Typical glomeruli, many with open capillary lumens. The expected cuboidal epithelium lining the Bowman’s capsule is apparent in many glomeruli.
Mild, multifocal interstitial inflammatory cell infiltrate. The remaining interstitium and tubules is unremarkable.
(87504)
*Comments:*
*Pathology to comment*

Brain

Slide shows sagittal sections of one hemisphere demonstrating, meninges, cerebral cortex with typical cellular lamination, corpus callosum, ventricles and representative choroid plexus, subfornical organ, thalamus, hypothalamus, cerebellum and pons. Unremarkable cerebellum with typical architecture and Purkinje cells. No obvious neuronal loss, and the myelination appears normal.
No lesions of significance
(87501)

#### 72

Macro Observations

Fixed spleen, liver, brain & kidney
Kidney 10×6×4mm
No macroscopic lesions identified.

Micro Observations

Adar1P195A/− lfih1−/−

Liver

Section shows typical liver parenchyma including hepatocytes, Kupffer cells, portal triads and central veins.
No lesions of significance
(87499)

Spleen

Unremarkable follicular pattern identified with typical red and white pulp micromorphology. Mild extramedullary haematopoiesis identified in the red pulp, a common feature in the mouse spleen.
Section also includes a small portion of unremarkable pancreatic exocrine tissue.
No lesions of significance
(87498)

Kidney

H&E section show cortex, medulla and papilla regions of the kidney.
Typical glomeruli, many with open capillary lumens. The expected cuboidal epithelium lining the Bowman’s capsule is apparent in many glomeruli.
Focal inflammatory cell aggregate (lymphocytic).
The remaining interstitium and tubules is unremarkable.
(87500)
*Comments:*
*Pathology to comment*

Brain

Slide shows sagittal sections of one hemisphere demonstrating, meninges, cerebral cortex with typical cellular lamination, corpus callosum, ventricles and representative choroid plexus, hippocampus, subfornical organ, thalamus, hypothalamus, cerebellum and pons. Unremarkable cerebellum with typical architecture and Purkinje cells. No obvious neuronal loss, and the myelination appears normal.
No lesions of significance (87497)

#### 97

Macro Observations

Fixed spleen, liver, brain & kidney
Kidney 10×5×4mm
No macroscopic lesions identified.

Micro Observations

Adar1P195A/E861A

Liver

Section shows liver parenchyma including hepatocytes, Kupffer cells, portal triads and central veins.
Mild, multifocal inflammatory cell aggregate, resembling extramedullary haematopoiesis. (87495)
*Comments:*
*Pathology to comment*

Spleen

Unremarkable follicular pattern identified with typical red and white pulp micromorphology. Mild extramedullary haematopoiesis identified in the red pulp, a common feature in the mouse spleen.
No lesions of significance
(87494)

Kidney

H&E section show cortex, medulla and papilla regions of the kidney.
Typical glomeruli, many with open capillary lumens. The expected cuboidal epithelium lining the Bowman’s capsule is apparent in many glomeruli.
The interstitium and tubules is unremarkable.
No lesions of significance
(87496)

Brain

Slide shows sagittal sections of one hemisphere demonstrating, meninges, portion of olfactory lobe, cerebral cortex with typical cellular lamination, corpus callosum, ventricles and representative choroid plexus, hippocampus, subfornical organ, thalamus, hypothalamus and cerebellum. Unremarkable cerebellum with typical architecture and Purkinje cells. No obvious neuronal loss, and the myelination appears normal.
No lesions of significance(87493)

#### Comment / Plan

Case APN22/016SVI(C. Walkley) to has been referred to text removed CW090522
28th March, 2022

### Supplementary Pathology Report

#### 409 (control)

No abnormalities detected

#### 410 (control)

No abnormalities detected

#### 418 (control)

No abnormalities detected

#### 104

No abnormalities detected

#### 124

No abnormalities detected

#### 125

No abnormalities detected

#### 126

126(87463)- spleen-normal follicular pattern (unusual plane of section)
No other abnormalities detected

#### 129

129(87475)- spleen-normal follicular pattern (unusual plane of section)
129(87474)-liver-focal hepatocellular necrosis with PMN and Kupffer cell reaction (common incidental finding); accentuation of acinar pattern due to mild hydropic degeneration in periacinar and mid zonal hepatocytes.
No other abnormalities detected.

#### 131

131(87487)-liver-focal hepatocellular necrosis with PMN and Kupffer cell reaction (common incidental finding); accentuation of acinar pattern due to mild hydropic degeneration in periacinar and mid zonal hepatocytes.
No other abnormalities detected

#### 70

70(87508) kidneys - tubular protein casts (common incidental finding) No other abnormalities detected

#### 71

71(87504) kidneys - Glomerular tuft hypercellular with basement membrane thickening and capsular thickening. Interstitial mononuclear cell infiltrate with mesenchymal-like cells with hyperchromatic, elongated nuclei.
Tubular protein casts (common incidental finding)
No other abnormalities detected

#### 72

72(87500) kidneys - single periarteriolar mononuclear cell cuff.
No other abnormalities detected

#### 97

97(87495)-liver- focal hepatocellular necrosis with PMN and Kupffer cell reaction (common incidental finding); accentuation of acinar pattern due to mild hydropic degeneration in periacinar and mid zonal hepatocytes
No other abnormalities detected

### Summary

Please see individual samples in the above report for comments.
PMN=Polymorphonuclear leukocytes
text removed CW090522
30th March, 2022

### Supplementary Neuropathology Report

#### 409 (control)

No abnormalities detected

#### 410 (control)

No abnormalities detected

#### 418 (control)

No abnormalities detected

#### 104

No abnormalities detected

#### 124

No abnormalities detected

#### 125

No abnormalities detected

#### 126

No abnormalities detected

#### 129

No abnormalities detected

#### 131

No abnormalities detected

#### 70

No abnormalities detected

#### 71

No abnormalities detected

#### 72

No abnormalities detected

#### 97

No abnormalities detected

#### Summary

No abnormalities detected. Cerebrum; often much mechanical damage in the form of numerous dark neurons; cerebellum appeared normal Brain-The samples presented in this case bear no lesions of significance.
text removed CW090522
30th March, 2022
Phenomics Australia advises all research groups that images or results obtained through the services are to be acknowledged in resultant publications. Example acknowledgment: “This study utilised the Phenomics Australia Histopathology and Slide Scanning Service, University of Melbourne.”

